# Unearthing the rhizosphere microbiome recruited by ancestral bread wheat landraces

**DOI:** 10.1101/2025.05.08.652585

**Authors:** Maria C. Hernandez-Soriano, Frederick J. Warren, Falk Hildebrand, Luzie U. Wingen, Anthony J. Miller, Simon Griffiths

**Affiliations:** John Innes Centre, Norwich Research Park, Colney Ln, Norwich NR4 7UH, UK; Quadram Institute Bioscience, Norwich Research Park, Rosalind Franklin Rd, Norwich NR4 7UQ, UK; Earlham Institute, Norwich Research Park, Norwich, Norfolk, NR4 7UQ, UK

**Keywords:** wheat, landrace, root traits, rhizosphere, soil, microbiome

## Abstract

Crop root traits that modulate the soil microbiome can turn the tide of reduced fertility in intensively farmed land by optimising nutrient availability and resilience to environmental stresses. Advantageous genetic diversity that allows adaptation to nutrient availability is present in historic crop genotypes. The Watkins collection of bread wheat landraces is an unexploited resource, carrying untapped phenotypic traits. Here, we show that the rhizosphere microbiome assembly of these landraces is distinct compared to elite varieties, specifically those that come from ancestral groups (AGs) not used in modern breeding.

We used 16S rRNA sequencing to identify changes in microbial communities of rhizosphere soil collected from 81 landraces and two elite varieties. We found high similarity in microbiome recruitment between the elite cultivars and the two AGs genetically closest to the elite. The rhizosphere microbiome of five AGs genetically distant from the elite cultivars showed significant differences in the abundance of taxa involved in nitrogen and carbon turnover, keystone taxa and associations within the microbial network. Our findings suggest that genes to recruit or suppress microbial taxa in the rhizosphere are shared by landraces from these AGs. Selective breeding for traits to control microbial functions can enhance soil productivity and crop performance.

## Introduction

Recent wheat genomic research and our capacity to identify traits of interest has made a major leap forward by whole-genome re-sequencing of the A.E. Watkins collection (Wingen et al. 2014) of landrace cultivars (LCs) and over 200 elite varieties ^1^. This research has advanced our knowledge of the genetic and phenotypic diversity of the Watkins LCs and categorised them into seven ancestral groups (AGs, individual groups designated as AG1 to AG7). Five of the AGs (AG1, AG3, AG4, AG6 and AG7) are phylogenetically distant from elite cultivars and provide a reservoir of unexplored functional diversity to support ecosystem functions ^1–3^, while AG2 and AG5 are genetically close to modern cultivars.

Wheat domestication and modern breeding have been keystones for the provision of food to humanity ^4^, but have resulted in a substantial reduction of diversity in the wheat genome ^5,6^, decreasing crop resilience and capability to control key soil processes.

Studies of trait variation have long been biased towards above-ground traits ^7^, but the identification of root traits to enhance soil productivity and plant resilience to abiotic and biotic stresses was overlooked and has now become a major research focus ^8,9^. Besides the fundamental role of roots in plant water and nutrient acquisition, they are also crucial for the active recruitment of microorganisms through the release of specialised root exudates. This strongly contributes to plant fitness through modulation and control of the rhizosphere microbiome composition and associated functions ^10,11^. Tailoring soil microbiome assembly allows plants to control key soil ecosystem functions such as the provision of nutrients, carbon (C) storage and physical stability, suppression of pathogens ^12,13^ and general support of healthy growth ^14^. However, our understanding of how plant genetics controls microbiome assembly is still developing, with key knowledge gaps remaining regarding the specific genes and pathways involved in microbiome interactions and how this cross-talk impacts plant health and resilience ^15–17^.

Soil microbiome composition and dynamics are driven by physical, chemical, and biological processes through complex ecological interactions, particularly plant-microbiome feedback and microbial interaction networks ^11,18^. Plants can alter soil physicochemical properties and modulate microbial activity in response to environmental changes ^8,19^ through the release of organic acids, amino acids, proteins, sugar, phenolics and specialised secondary metabolites ^10,13,20^.

Historic varieties of crops hold still uncharacterized, likely genotype-specific root traits to efficiently control the microbial communities in the rhizosphere^21^. There is potential for selection of soil microbiome-based traits in plant breeding, but identifying these root traits remains challenging due to the complexity of the interactions between roots, soil and environmental conditions ^9,22^.

Targeting root traits related to N and C dynamics is instrumental to the future of sustainable crop systems ^23^. Particularly, the capability of certain crop genotypes to release biological nitrification inhibitors (BNIs) into the rhizosphere to suppress the microbial transformation of ammonium into nitrate has been investigated for its potential to increase N use efficiency (NUE) ^24^. However, to date, this trait has only been tested using pure cultures of nitrifiers such as *Nitrosomonas europaea* or enzymatic assays ^25,26^. Nitrogen (N) cycling in soil, its availability for plant uptake, and the resulting impact on the environment involve complex networks of microbes and their metabolic pathways ^27,28^. Therefore, phenotyping this trait requires a holistic understanding of BNI impact on N cycling guilds ^29^ using high-throughput sequencing to study the microbiome composition of rhizosphere soil.

Similarly, several root traits have been reported to impact C stabilisation in soil ^30^. Root exudates are a significant source of C input into the soil, but they also influence the role of microbial communities in C cycling, potentially leading to increased soil C sequestration or organic matter decomposition and nutrient release ^31,32^. However, phenotyping these traits and guiding their delivery into new cultivars remains challenging.

Root traits are particularly relevant for staple crops such as wheat. Breeding programs focused on increased yield have reduced the ability of wheat to control nutrient cycling ^33^. Although modern, elite cultivars are highly competent at taking up N in controlled supply systems, they are less efficient at managing losses, perform ineffectively on N-deprived soils ^33–35^, and often have a lower seed protein content than LCs ^36^.

Several studies have addressed wheat-soil microbiome interactions. However, research addressing the relevance of wheat domestication history and implications for wheat microbial ecology remains scarce ^37^. Generally, wheat historic LCs present wider genetic diversity than their elite relatives ^5^, and might recruit more complex microbial community networks in their rhizosphere than post-Green Revolution cultivars ^6^. Recent research has also reported significant shifts in bacterial and fungal communities for wild and domesticated tetraploid wheat ^38^.

Here, we aim to link the genetic composition of wheat genotypes to the preferential recruitment of soil microbiomes by analysing the rhizosphere of 81 LCs. We found that genetically closely related wheat genotypes present more comparable microbiome assemblies in the rhizosphere than genetically distant genotypes, as evidenced by differences in microbiome diversity, abundance of taxa involved in N transformation and C turnover and microbiome network properties.

Our results pave the way to explore other unexploited root traits controlling specialised microbiome functions.

## Results

### Genetic diversity among wheat genotypes is consistent with the microbiome diversity in their rhizosphere

Firstly, we investigated differences in microbiome assembly in the rhizosphere soil (Supplementary Table 1) across the wheat genotypes according to their organisation into genetically similar groups (AGs and elite cultivars, Supplementary Table 2). We found marked dissimilarities in beta diversity between elite cultivars and AG1, AG3 and AG4 (visualised by multidimensional scaling in Fig. 1a). The results of PERMANOVA analysis using Bray-Curtis and Jaccard distances are summarised in Supplementary Table 3. The coefficients and p-values adjusted for multiple comparisons using FDR (q-values) indicate significant differences (q<0.1) in rhizosphere microbiome composition between modern cultivars and AG1 according to the Jaccard index (F=1.8807, q=0.056) and Bray-Curtis dissimilarity (F=2.3791, q = 0.084). The results also suggest a trend in dissimilarity for AG3 and AG4, considering the high coefficient values and low q values (jaccard: AG3, F=1.4101, q = 0.151; AG4, F=1.6124, q = 0.151; bray: AG3, F=1.5906, q = 0.203; AG4, F=1.7983, q = 0.241).

**Figure 1.**
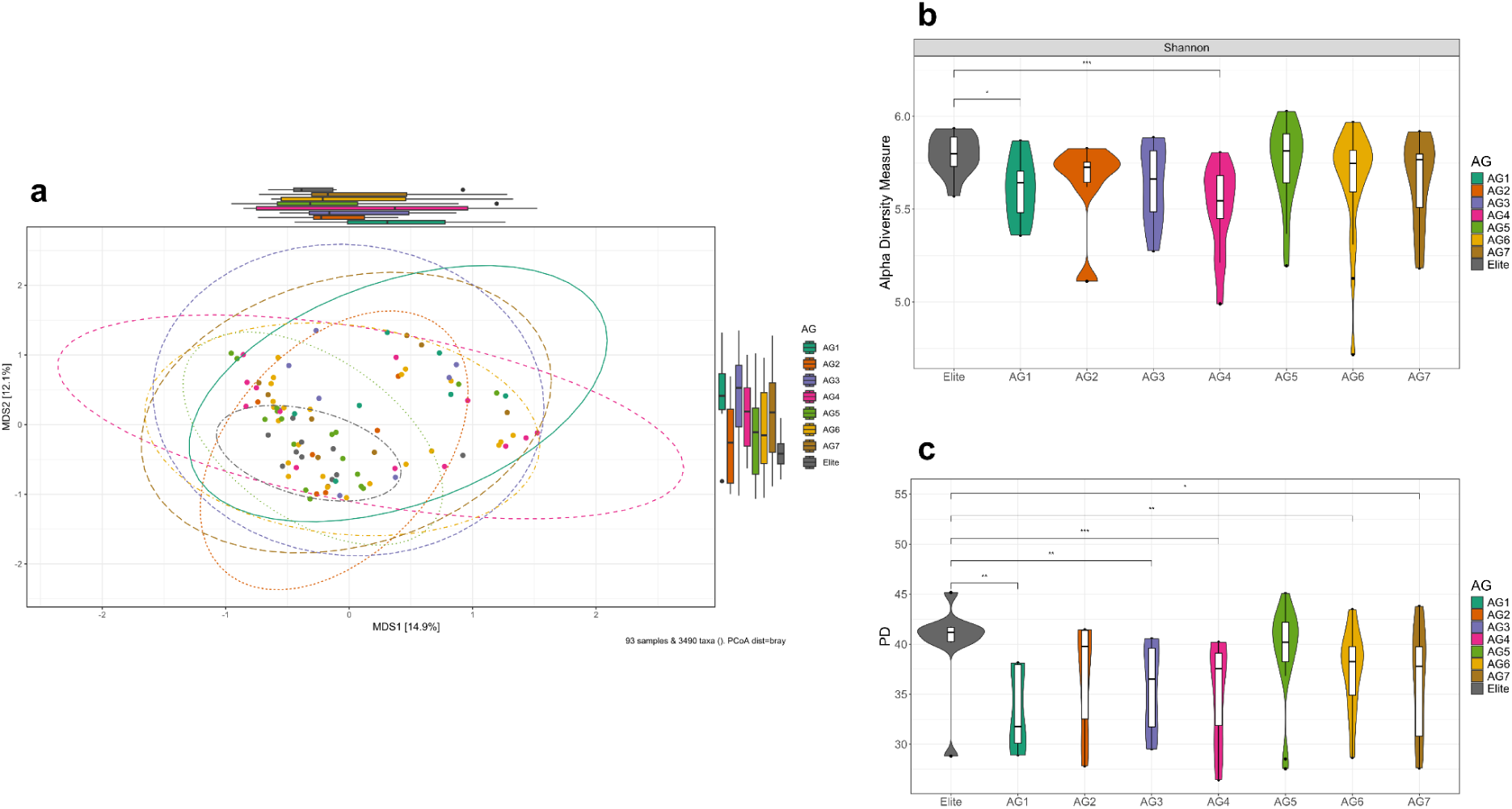
Microbial diversity in the rhizosphere soil of wheat landraces and elite cultivars. **(a)** PCoA of the beta diversity based on Bray-Curtis distance of ASV relative abundance data from rhizosphere soil collected from 83 genotypes of hexaploid wheat grown in the field (R package microviz ^1^). Colour code for the classification of the wheat genotypes into ancestral groups (AG) ^2^, closely related genetically, and elite (modern) cultivars. Ellipses were drawn based on a 95% confidence interval and represent samples from each wheat genotype group. **(b)** Alpha-diversity index (Shannon index) and **(c)** Faith’s phylogenetic diversity (PD) index across wheat genotypes, grouped into ancestral groups (AG) ^2^, closely related genetically, and elite (modern) cultivars. Significant differences in Shannon diversity and PD were calculated by pairwise comparison using the Wilcoxon test and the Benjamini-Hochberg method for the adjusted p-value (*p ≤ 0.05, **p ≤ 0.01, ***p ≤ 0.001).

The analysis indicates that AG2 and AG5 present the least different microbiome recruitment compared to elite cultivars, according to the high q ( q>0.6) and F values (Jaccard: AG2, F=0.972; AG5, F=1.095; Bray: AG2, F=0.9329; AG5, F=1.1271). This similarity in rhizosphere microbiome recruitment is consistent with the reported genetic overlap of the LCs within these AG groups and elite cultivars ^1^.

To further investigate the diversity of the microbial community composition in the rhizosphere soil of the AGs compared to that of elite varieties, we used alpha diversity (Shannon index) and Faith’s phylogenetic diversity (PD) measurements. The rhizosphere communities from LCs as grouped within AG1 and AG4 showed significantly lower diversity (p < 0.01 and p < 0.001, respectively) than those from the elite cultivars (Fig. 1b). The other AGs showed a similar rhizosphere community diversity to the elite cultivars, although individual LCs from those AGs also displayed a low diversity (see Supplementary Fig. 1-5 and Supplementary Table 4).

We found significant PD differences between the rhizosphere of the elite cultivars and all AGs except for AG2 and AG5 according to Faith’s PD measurement (Fig. 1c).

### Genetic proximity among wheat genotypes aligns with the microbiome recruitment profile in their rhizosphere

The core microbiome in the rhizosphere soil across all the wheat genotypes assessed (Supplementary Fig. 6) represents the top 50 taxa at the genus level and their prevalence at increasing abundance thresholds. Focussing on the top 15 genera and their contrasting abundance among AG and elite group soils (Fig. 3), we found that the most abundant phyla were Proteobacteria (33.2-36.2%), Actinobacteriota (23.9-35.2%), Thermoproteota (4-11.2%), Acidobacteriota (6.8-10%) and Chloroflexota (4.7-6.1%) (Fig. 2a and Supplementary Table 5). We observed a distinctive lower abundance of the ammonia-oxidising archaea (AOA) *TA-21* (Thermoproteota) in the rhizosphere soil of AG1, AG3 and AG7 compared to the elite cultivars. Subsetting the key nitrifier taxa found in the rhizosphere soil (Nitrososphaerales, Nitrospirales, Gemmatales) confirmed the dominance of the AOA *TA-21* (Fig. 2b). Also, several genera were only found in the rhizosphere soil of specific groups (abundance threshold > 0.01). For instance, *Ketobacter* was only found in AG7, while *Terrimicrobium* was only detected in AG2 and AG7.

**Figure 2.**
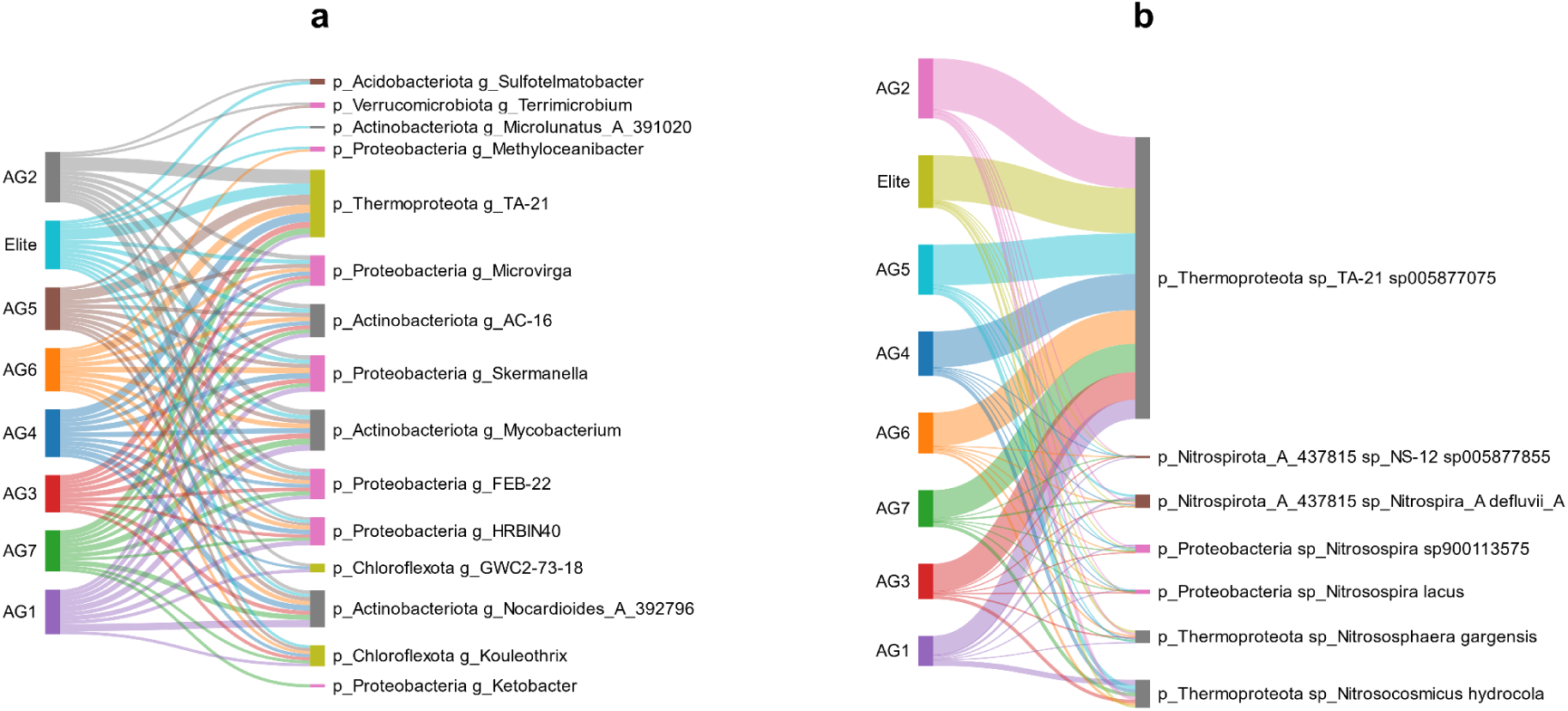
Differences in core microbiome and abundance of nitrogen cycling guilds in rhizosphere soil between wheat landraces and elite cultivars. **(a)** Distribution of the top 15 most abundant taxa identified in the rhizosphere soil collected from 83 genotypes of hexaploid wheat grown in the field. Colour code for the classification of the wheat genotypes into ancestral groups (AG) ^2^ closely related genetically, and elite (modern) cultivars, ranked by taxonomic group phylum and genus, and decreasing abundance; **(b)** distribution of taxa belonging to the order Nitrososphaerales, Nitrospirales and Gemmatales confirmed the dominance of the *AOA TA-21* in the rhizosphere soil collected from 83 genotypes of hexaploid wheat grown in the field ^3^. Colour code for the classification of the genotypes into ancestral groups (AG) ^2^, closely related genetically, and elite cultivars, ranked by taxonomic group, phylum and species, and decreasing abundance. The relative abundance of Thermoproteota is provided in Supplementary Table 5. The species are categorised into the order Nitrospirales, Nitrososphaerales and Burkholderiales. The abundance of ammonia-oxidising archaea (phylum Thermoproteota) was significantly lower in the rhizosphere of AG1 (*TA-21*) and AG7 (*Nitrososphaera*) (Supplementary Tables 8 and Supplementary Fig. 7a). On the left band, the Sankey diagrams show the wheat genotype group, which splits into phylum and genus or phylum and species (right). The widths of the bands are linearly proportional to the relative abundance in the rhizosphere of the wheat genotypes. Sankey diagrams were created using SankeyMATIC (https://sankeymatic.com/).

**Figure 3.**
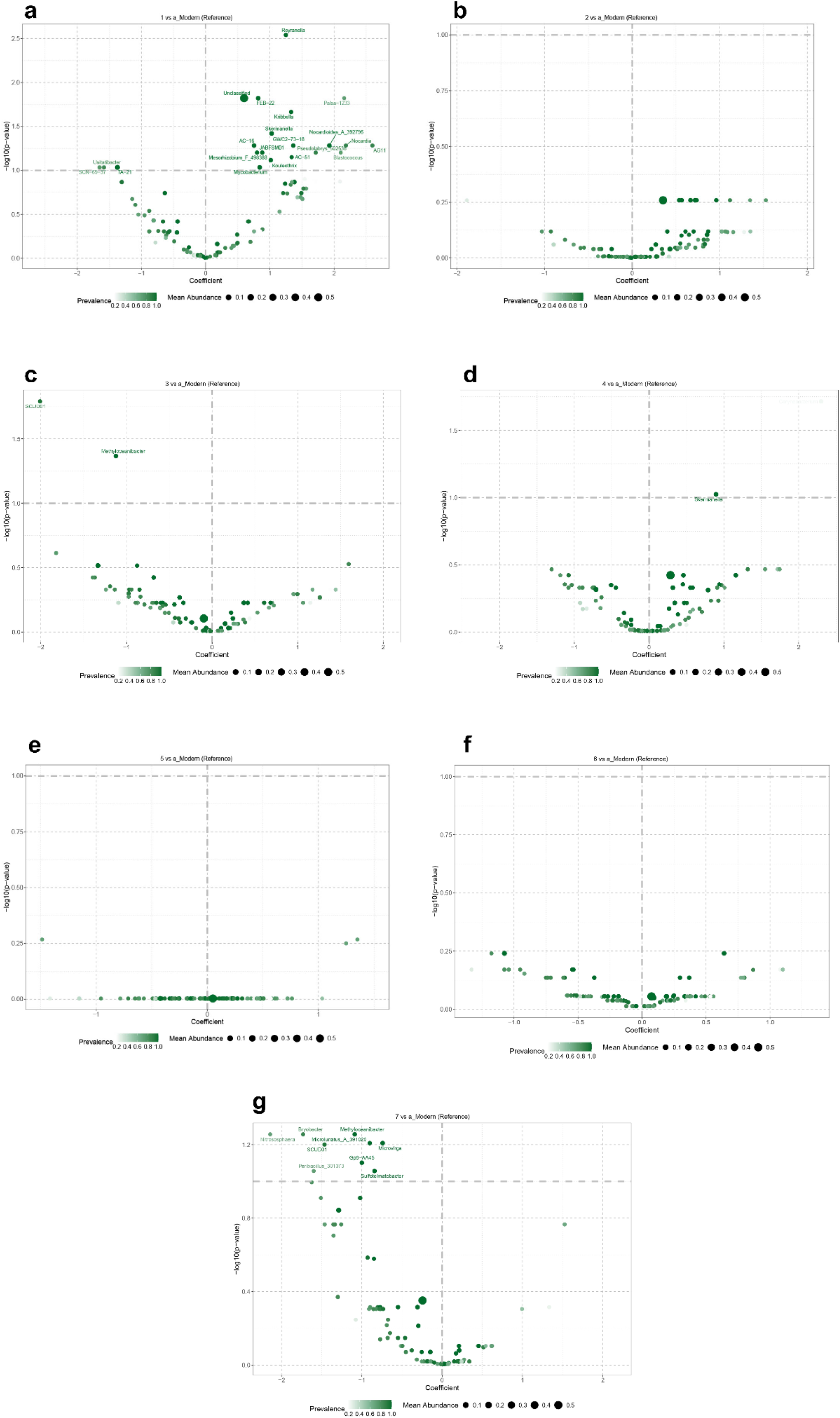
Differential abundance of microbial taxa in the rhizosphere soil of wheat landraces and elite cultivars. Volcano plots showing differential abundance of taxa at the genus level between elite (modern) cultivars of hexaploid wheat and historic genotypes grouped into ancestral groups (AG) ^2^, AG1 **(a)**, AG2 **(b)**, AG3 **(c)**, AG4 **(d)**, AG5 **(e)**, AG6 **(f)** and AG7 **(g)**. The position along the x-axis represents the abundance fold change. The dashed line shows the threshold of significant differential taxa (|log 10 (FC)| > 1). Taxa located on the left side are less abundant in the rhizosphere of the AGs compared to the elite cultivar (p-value, Supplementary Table 8). Taxa located on the right side are more abundant in the rhizosphere of the AG. Volcano plots and linear models for differential abundance analysis (LindDA) at phylum and order levels are provided in Supplementary Figure 7 and Supplementary Tables 6 and 7. The LinDA analysis was performed using the R package microbiomeStat ^4^.

To further explore differences in the microbiome recruitment across wheat genotypes, we performed differential abundance univariate testing (LinDA). This confirmed the similarity in microbial recruitment in the rhizosphere soil between elite cultivars and AG2 and AG5 at the genus level (Fig 3, plot b and e), and at the phylum and order level (Supplementary Fig. 7, Supplementary Tables 6-8).

Compared to the rhizosphere of the elite cultivars, several taxa presented differential abundance in the rhizosphere of other AGs (Fig 3, plot a, c, d, f and g, Supplementary Fig. 3, Supplementary Tables 6-8). Generally, at the phylum level, Verrucomicrobiota, Thermoproteota, Firmicutes, Nitrospirota, Acidobacteria and Desulfobacterota were significantly less abundant in AG1 compared to the elite cultivars. Proteobacteria, Chloroflexota, Actinobacteria and Gemmatimonadota appeared significantly more abundant in AG4 compared to elite cultivars (see Supplementary Fig. 7).

The statistical association between the genetic diversity of AGs and elite cultivars and the rhizosphere microbial composition according to the LinDA analysis was further confirmed by the multivariate association analysis performed with MaAsLin2 (phylum level: Supplementary Fig. 8, Supplementary Table 9 and genus level: Supplementary Fig. 9, Supplementary Table 10). This confirms the significant differences in microbiome recruitment in the rhizosphere between elite cultivars and AG1, AG3 and AG4. The q-values indicate that two unresolved genera were significantly enriched in AG1 and AG2 soils compared to the elite cultivars (Supplementary Fig. 10). Consistent with the results from the LinDA analysis, AG2 and AG5 present comparable microbiome recruitment to the elite varieties. In contrast, AG1 and AG4 presented marked differences with the elite varieties in the abundance of several taxa recruited in the rhizosphere. Particularly, the MaAslin2 analysis confirmed a significant increase in Proteobacteria and a substantial decrease in Thermoproteota for AG1 compared to the elite cultivars, and a reduction in Acidobacteria for AG1 and AG4 compared to the elite cultivars.

Using imputed ecological functions associated with the genera that were differentially abundant between AGs and elite cultivars, we found organic matter decomposition, ammonia-oxidation and potential N fixation, nutrient cycling, and plant growth promotion being most often different between AGs and elite cultivars (Supplementary Table 11).

Pairwise comparison of the predicted functional annotations for N metabolism using the elite cultivars as reference (Supplementary Fig. 11, Supplementary Table 12) indicated lower abundance of genes involved in ammonia oxidation and complete nitrification in the rhizosphere of AG1, consistent with the lower abundance of the AOA *TA-21* in the rhizosphere (Fig. 2, Fig. 3, Supplementary Table 8).

For genes involved in C fixation, pairwise analysis using the elite cultivars as reference indicated lower abundance of genes involved in C fixation pathways in AG1 (Arnon-Buchanan, 3-Hydroxypropionate bi-cycle), AG3 (Arnon-Buchanan) and AG4 (Calvin cycle), and higher abundance of genes involved in the conversion of acetyl-CoA to acetate for AG3 and AG6 (Supplementary Fig. 12, Supplementary Table 13).

### Microbiome association networks confirm the link between wheat genetic diversity and microbiome recruitment in the rhizosphere

Next, we used microbial network analysis to investigate associations in rhizosphere microbiome taxa across the wheat genotypes. The microbial co-occurrence network in the rhizosphere soil had low modularity (0.03) and moderate clustering (0.63) (Supplementary Table 14). Three clusters of microbial taxa were identified across the wheat genotype rhizospheres (Fig. 4a), with 72 nodes at the order level and 129 at the genus level. The most influential and well-connected taxa at the order level were Acidimicrobiales, Actinomycetiales, Geminicoccales, Mycobacteriales and Propionibacteriales, identified as hub taxa according to the centrality measures (Supplementary Table 14). Chitinophagales, Mycobacteriales, Geminicoccales and Burkholderiales co-occurred with multiple clusters, acting as bridges (high betweenness).

**Figure 4.**
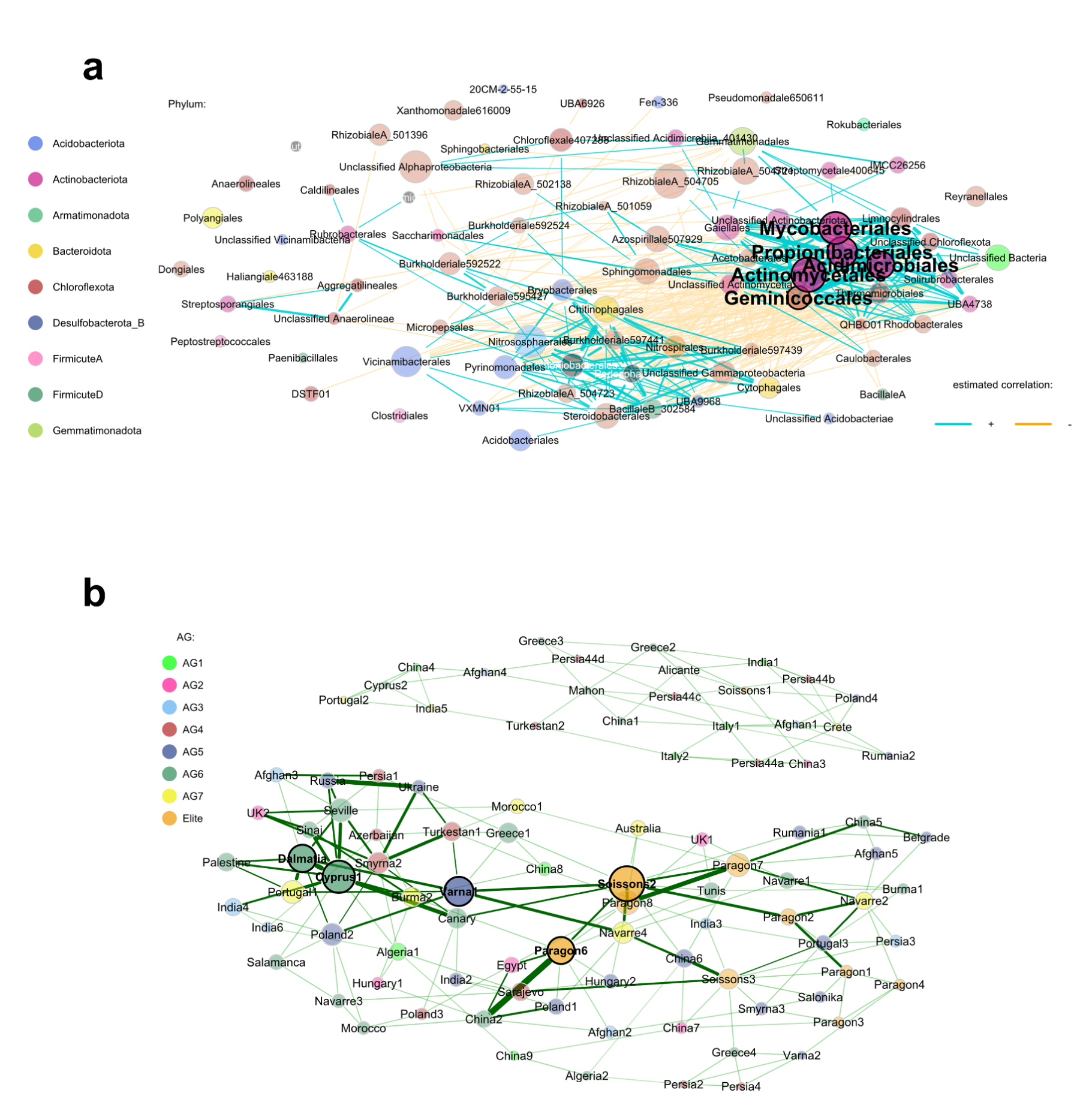
Microbial co-occurrence network of the rhizosphere soil across wheat genotypes and dissimilarity network. **(a)** Microbial co-occurrence network ^5^. Nodes are taxa at the order level, colour-coded to represent phylum. Node size indicates relative abundance. Associations between taxa (Pearson correlations) are displayed as turquoise (positive) and orange (negative) lines. **(b)** Dissimilarity network ^5^, nodes are wheat genotypes, colour-coded to represent whether they are categorised into ancestral groups (AGs) ^2^ or elite cultivars. Node size corresponds to eigenvector centrality, which was used to identify wheat genotypes that are hubs in the network. Similarity between nodes (wheat genotypes) is indicated as green lines (Aitchinson distance).

Cluster A gathered taxa involved in N metabolism, including the most prevalent nitrifiers (Thermoproteota, Nitrososphaerales and Nitrospirales). Cluster B shows a dominance and prevalence of Actinomycetales, Acidimicrobiales, Mycobacteriales, Propionobacterales, Geminicoccales and several taxa within the Chloroflexota phylum. Taxa within both clusters were positively correlated, while taxa between clusters were generally negatively correlated. Cluster C was relatively small, gathering only five taxa (phyla Acidobacteria and Chloroflexota) with negative or no correlations with the other clusters. Several taxa were disconnected, including Reyranellales, Rokubacteriales, Pseudomonadales and Clostridiales.

To explore the differences in microbiome network structure between the elite cultivar and each AGs we constructed differential (Supplementary Fig. 13) and association (Supplementary Fig. 14) networks. The network comparison analyses (Supplementary Tables 15 to 21) also indicated low modularity (<0.05). The Jaccard indices derived from the comparison analyses (summarized in Supplementary Table 22) confirmed that AG1, AG3 and AG7 networks significantly differ from the elite cultivars network for several centrality measures. The AG2, AG4, and AG6 networks didn’t present any significant differences for any of the measures compared to the elite cultivars, and the AG5 network only presented significantly different closeness. Particularly, we found higher positive associations between taxa (positive edge percentage) for AG1, AG4 and AG6 networks than for the elite cultivars network. The positive associations were lower for the AG5 network than for the elite cultivars, while AG2, AG3 and AG7 presented no differences.

Although the Jaccard index indicates no significant differences for the hub nodes within each comparison, the hub nodes corresponded to different taxa among the network comparisons (Supplementary Fig. 13 and 14 and Supplementary Table 23).

In particular, several differences in degree were identified for hub taxa across the different networks. We found a higher degree for the AOA *Nitrososphaeraceae* genus and a lower degree for the AOA *TA-21* in the AG1, AG5 and AG6 networks compared to the elite cultivars. Both genera presented a higher degree for the AG3 network. *TA-21* also presented a higher degree for the AG7 network. These genera were not identified as hubs for the AG2 and AG4 networks when compared to the elite cultivars, while we observed contrasting differences in degree for taxa such as *Burkholderiaceae* (AG2, AG4), *Skermanella* and *Micrococcaceae* (AG4).

Additionally, according to the topological measures of the global comparison networks, the positive edge percentage indicates higher network complexity in the rhizosphere of AG1, AG4 and AG6 compared to elite cultivars.

To gain further insights into the differences in taxa associations between the elite cultivars and each AG we examined the connections and relationships between nodes in the association (Supplementary Fig. 13) and differential netwoks (Supplementary Fig. 14). These connections can reveal patterns, clusters, and the flow of information or influence within the network. We observed shifts in abundance and associations for several taxa between the elite cultivars and the AGs rhizosphere. Particularly, *Nitrososphaeraceae* and *TA-21* are positively associated in the rhizosphere of the elite cultivars but not associated in the rhizosphere of AG1 or AG3. We also observed a shift from no association in the rhizosphere of the elite cultivar to negative association between *Nitrososphaeraceae, TA-21* and taxa involved in organic matter decomposition and nutrient cycling (*Ketobacter, Skermanella*, *Micrococcaceae* and *Alphaproteobacteria*) in the rhizosphere of AG1 and AG3. The differential networks feature the similarity between the associations in the rhizosphere microbiome of the elite cultivars and AG2 and AG5.

Finally, to assess the similarity and relationship of the AGs based on their rhizosphere microbiome assembly, we constructed a dissimilarity network where the nodes were individual genotypes. The network presented a moderate level of community structure, with a modularity of 0.43 and a clustering coefficient of 0.36. Most of the genotypes were identified as nodes (77 nodes at the order level, 75 at the genus level). Overall, elite cultivars and some accessions within the AG5, AG6 and AG7 were the most influential genotypes within the network. The analysis gathered the genotypes into three clusters (Fig. 5b, Supplementary Table 24). The network showed strong correlations between the elite cultivars and most LCs within the AG2 and AG5 groups. Accordingly, 57% of LCs within AG2 and AG5 (13 LCs) were found in the same cluster as the elite cultivars (cluster 1), three LCs (13%) within cluster 2, and 30% (7 LCs) in cluster 3. Clusters one (29 genotypes) and three (35 genotypes) showed strong correlations between genotypes. Cluster two (23 genotypes) was disconnected from the other clusters and gathered LCs from all AGs, with just one sample from the elite cultivars, one LC from AG2 and two LCs from AG5.

## Discussion

The capacity of particular wheat genotypes to control soil microbiome assembly and functions by releasing specialised metabolites into the rhizosphere can improve soil productivity and crop performance ^39^. The Watkins landrace collection of bread wheat is a germplasm resource bearing unexploited assets of these functional root traits that can be identified and introduced into modern varieties by breeding programs ^1,40^. Traits related to wheat control of microbial N and C metabolism are highlighted in this study for their importance in agricultural systems.

Differences in the root microbiome recruitment among crop genotypes have been reported for several crops, although most studies only considered a limited number of genotypes. Shifts in rhizosphere microbiome composition have been reported between two genotypes of barley grown in rhizoboxes ^41^, among four inbred lines of pearl millet grown in pots ^42^, among three genotypes of rice grown in a field trial ^43^ and 19 genotypes grown in pots ^44^, among eight genotypes of hexaploid wheat grown in the field ^6^, and among 44 genotypes of tetraploid wheat grown in a field trial ^38^. While these studies have shed light on the potential of root traits to modulate soil microbiome functions, the phenotyping of these traits remains unexplored.

Here, we examined the bacterial and archaeal communities in the rhizosphere soil collected from 81 genetically diverse LCs and two elite cultivars of hexaploid wheat using amplicon sequencing (16S). We linked the microbiome fingerprint to the genetic diversity of the wheat genotypes, according to the classification of the LCs in AG groups established for the Watkins landrace collection of bread wheat ^1^.

The lower diversity observed in the rhizosphere soil of the LCs compared to elite cultivars (Fig. 1) is consistent with previous research on wild genotypes of rice, maize and hexaploid wheat ^6,44,45^, while a study on tetraploid wheat reported the opposite trend ^38^. These contrasting results may be explained by the influence of other factors on microbiome recruitment, particularly differences in root architecture among genotypes and soil environment ^46^. Previous research has indicated that microbiome diversity decreases with increasing availability of resources ^13^.

Accordingly, we postulate that LCs might support an environment richer in nutrients compared to the elite cultivars. The interpretation of this trend should also consider that associations, rather than richness, determine the complexity of the microbiome assembly in the rhizosphere ^18,47,48^.

The core microbiome of the rhizosphere soil across the wheat genotypes (Fig. 2 and Supplementary Fig. 6) was consistent with previous field studies ^6,13,49–51^, reporting the prevalence of phyla involved in decomposition of organic matter, nutrient cycling and potential N fixation, Proteobacteria, Actinobacteriota, Acidobacteriota and Chloroflexota. The high abundance and prevalence of Thermoproteota is consistent with the low dose of N-fertiliser applied in this study and the field soil properties ^52^.

Exploring the microbiome communities at finer taxonomic resolution confirmed the ubiquity and dominance of the AOA genus *TA-21* ^53^ (Fig. 2b), plant-growth-promoting bacteria *Skermanella* and *Microvirga* and bacteria involved in C and N metabolism, particularly *Mycobacterium* and *Kouleothrix* (Fig. 2a, Supplementary Table 11).

Several genera related to organic matter decomposition and nutrient cycling (Supplementary Table 11) were AG or elite genotype-specific, including *Terrimicrobium*, *Microlunatus* and *Ketobacter*. These taxa, particularly *TA-21,* can be proposed as candidates to be indicator taxa ^47,53^ since their prevalence is selective to specific habitats such as crop rhizosphere soils, and land management strategies ^52^. These results prompt further research on the genetic advantage of modulating these taxa to enhance crop resilience and fitness.

Differential abundance analysis further evidenced that wheat genetics drives plant microbiome assembly. The similarity in rhizosphere microbiome composition among the elite cultivars and genetically closely related AGs ^1^ (Fig.2, Fig.3b and 2e and Supplementary Fig. 7) aligns with previous research reporting comparable microbiome recruitment across modern cultivars ^49^, underpinning the role of wheat genetics in microbiome recruitment in the rhizosphere. The genetic relatedness of AG2 and AG5, which originate mainly from Western and Central European regions, with the elite cultivars ^1^, is consistent with a comparable rhizosphere microbiome assembly. The AGs less genetically related to the elite cultivars and originated mainly from China (AG1), South /Central Asia (AG3), the Balkans and greater Iran (AG4) and South Europe/Asia (AG7) ^1^, differed in microbiome recruitment compared to the elite cultivars (Fig. 2a, 2c, 2d and 2g, Fig. 3 and Supplementary Fig. 7). The genetic makeup of LCs results from adaptation to local conditions driven by human intervention, and explains the divergence between wheat genotypes from different AGs ^1,54^. The difference in the abundance of several taxa in the rhizosphere microbiome found for AG1 and AG7 (Fig 3a and 3g) compared to elite cultivars suggests that LCs within these AGs bear traits to suppress the generally ubiquitous soil AOA genera *TA-21* and *Nitrosophaera.* We postulate that these LCs carry genes for releasing BNIs into the rhizosphere, expanding on previous research that reported the BNI activity of several wheat genotypes against pure cultures of ammonia-oxidising bacteria ^25^. Moreover, these wheat genotypes could also impact AOA by altering soil aggregation since these communities have been described to proliferate as biofilms in aggregates ^53^. The substantial enrichment of several bacterial genera involved in C and N metabolism in the rhizosphere of AG1 and suppression of several genera involved in nutrient cycling in the rhizosphere of AG7, compared to the elite cultivars (Fig. 3a and 3g and Supplementary Table 11), further highlights the genotype-specific strategy for microbiome recruitment and the potential to explore the associated advantages of these LCs under different crop system scenarios.

While exploring the differences in rhizosphere microbiome recruitment across wheat genotypes at the genus level provided fine taxonomic resolution, surveying the order level allows for a broader comparison and gives insights into genotype strategies to soil microbiome assembly. The enrichment in Rhizobiales, associated with N-fixation, Actinomycetales, Gemmatimonadales and Mycobacteriales, known for their role in the decomposition of organic matter, humification, soil aggregation and water retention (Supplementary Fig. 7 and Supplementary Table 11) in the rhizosphere of AG1 and AG4, compared to the elite cultivars, suggests an advantage for these LCs to modulate C and N turnover in soil to improve soil structure and provision of nutrients. This is further supported by the suppression of Acidobacteriota, Nitrososphaerales and Nitrospirota in the rhizosphere of AG1 compared to elite cultivars and the enrichment in Chloroflexota, reported to increase C fixation, P and N availability, in the rhizosphere of AG4 (Fig. 7 and Supplementary Table 11). The genotype-specific strategy to control microbial N metabolism and C turnover is also underpinned by the differences in the predicted microbial functions for N (Supplementary Fig.11) and C cycling (Supplementary Fig.12). Particularly, we observed increased microbial assimilation of ammonium in the rhizosphere of AG1 by nitrate reduction combined with a decrease of ammonia oxidation compared to the rhizosphere of the elite cultivars. This suggests that LCs belonging to AG1 contribute to a “closed” N cycle ^52^, likely through the release of exudates that support the recycling of N within the rhizosphere environment, preventing losses to the atmosphere or groundwater. Moreover, an increase in assimilatory nitrate reduction, also found for the rhizosphere of AG3 compared to the elite cultivars, contributes to plant growth and NUE ^53^. The increase in dissimilatory nitrate reduction observed in the rhizosphere of AG2 compared to elite cultivars contributes to N retention in soil and reduction of nitrous oxide emissions ^52,55^. Generally, nitrification is rather limited in the rhizosphere environment ^13^, but our results indicate that the ability to support recycling of N within the rhizosphere environment is limited to specific wheat genotypes. Key functions related to C fixation were decreased in the rhizosphere of AG1 and AG3 compared to elite cultivars, while functions associated with the degradation of organic matter were increased in the rhizosphere of AG3 and AG6. These results suggest that these LCs are more efficient at recruiting taxa involved in C turnover, promoting effective nutrient cycling and ready availability.

The microbial co-occurrence and dissimilarity networks constructed for the rhizosphere soil provided further insights into the wheat genotype-specific impact on microbial recruitment. Firstly, we hypothesise that the low modularity of the co-occurrence microbial network constructed for all samples reflects the susceptibility of the rhizosphere community to the influence of different wheat genotypes. Low modularity suggests a less structured, potentially more dynamic community with greater plasticity in responses to environmental changes ^13,18^, or more flexible interactions between taxa ^56,57^. Taxa with high betweenness, considered as “gatekeepers” (Chitinophagales, Mycobacteriales, Geminicoccales and Burkholderiales) often co-occurred within multiple clusters, indicating an important role as active symbiont to diverse ecotypes. These taxa actively interact with other microbes and/or the plant host, potentially benefiting both parties. Symbionts within a microbiome play a crucial role in shaping its diversity, stability, and resilience, ultimately influencing host health and adaptation to the environment.

The moderate modularity of the dissimilarity network constructed for the wheat genotypes implies a balance between specialisation (within modules) and generalist interactions (between modules), which might reflect the similarities in microbial recruitment across related wheat genotypes. The dissimilarity network identified several LCs disconnected from the elite cultivars and other LCs, suggesting a marked shift in the community composition and structure for those wheat genotypes.

The association and differential networks performed for each AG against the elite cultivars indicated significant differences in the structure of the microbial networks across all comparisons. Consistent with the trend observed from our results, we observed high similarity between the networks obtained for the elite cultivars and the AGs genetically closely related (AG2 and AG5) and distinct differences with the networks for AGs genetically less related. The shift in keystone taxa between AGs and the elite cultivars further evidences the genotype-specific ability to recruit or suppress particular taxa and influence the structure of the community network in the rhizosphere.

The hubs identified in the co-occurrence network indicate a general dominance of taxa involved in organic matter decomposition and N cycling, as expected for agricultural soils ^6,59^. The shift to antagonistic interaction between nitrifiers and taxa involved in C turnover from the elite cultivars to AG1 and AG3 networks (Supplementary Fig xx) indicates the capability of the LCs within these AGs to support microbes involved in C cycling while suppressing nitrifiers. Particularly, we observed that AOA shifted from a positive association in the elite rhizosphere to no association in the AG1 and AG3 rhizosphere. Also, we observed a shift to negative associations between AOA and genera involved in organic matter degradation and nutrient supply (*Skermanella*, *Ketobacter*, *Micrococcaceae*). This strategy reduces the competition for limited resources like oxygen, N and C, fostering the supply of nutrients from the breakdown of organic matter, while suppressing rapid transformation of N into soluble forms that can be easily lost to the environment ^52,60^. The shifts in network structure between elite cultivars and the AGs suggest the potential of LCs to prompt a more structured, clustered network, potentially more resilient to the impact of disturbances in the environment, such as fertiliser addition.

Network complexity is instrumental to support soil functions, since interactions in ecological networks are crucial for nutrient provision to crops ^48,56^

We postulate that the identification of nitrifiers as indicator taxa reflects the differential response of LCs and elite cultivars to changes in the environment, particularly their capability to suppress the enrichment in nitrifiers following application of N-fertiliser in crop systems. This can have important implications for reducing losses of N to the environment and maintaining an optimal balance in the microbiome network composition and interactions that support efficient cycling and the availability of natural nutrient resources. We propose that wheat traits that allow plant control of specific taxa, such as the AOA *TA-21,* in response to the addition of N-fertiliser can increase NUE and support natural microbial activity that supplies N to plants by supporting the balance in the competition for the utilisation of resources.

Overall, our results indicate that LCs genetically distant from elite cultivars drive plant-microorganisms interactions in the rhizosphere that might have been lost in modern elite cultivars. This likely occurs through the release of specialised metabolites in the rhizosphere, a trait that can be introduced in new wheat varieties. In this study, we focused on the legacy of different wheat genotypes on rhizosphere microbiome assembly. Future research will explore microbiome assembly at key plant developmental stages and the response of the microbial community in the rhizosphere to the application of N-fertiliser.

## Conclusions

Wheat genotype-specific root traits can harness differentiated ecological roles of root-associated microbiomes, likely through the production and release of specialised metabolites to modulate below-ground nutrient and carbon cycles and plant growth. To the best of our knowledge, our results provide the first evidence linking wheat genetic diversity to genotype-specific impact on microbial diversity, differential abundance and network structure in the rhizosphere soil.

Following on previous research that classified wheat genotypes according to genetic relatedness, we report the alignment of this classification with the strategies to recruit microbial communities in the plant rhizosphere.

Our results indicate that certain wheat LCs from the Watkins collection recruit a distinctive microbiome profile in their rhizosphere and foster beneficial associations in the microbiome network. This capability to modulate the soil microbiome and its associated functions seems limited in the elite cultivars examined. Particularly, LCs from the AG1, AG3 and AG7 categories, genetically distant from the elite cultivars, presented a significant reduction of nitrifier communities and associated genes for N metabolism. Our findings provide valuable insights into the mechanisms underlying plant-microbiome interactions and how to harness the rhizosphere microbiome for crop improvement in sustainable agriculture.

Future research will look into breeding these root traits into new wheat cultivars, with particular attention to the ability of plants to control nitrogen and carbon microbial metabolisms.

## Materials and methods

### Field trial

The Watkins core set (119 LCs) and two elite cultivars, Paragon and Soissons, were grown in 2020 in randomised field plots of 1 m × 1m (three replicates). The soil was a sandy loam, pH 8.4, with 2.6 % w/w organic matter content. Further details on the soil properties are provided in Supplementary Table 1. The plants were treated with xxx and supplied with N-fertilizer at 48 kg N ha^-1^ at tillering stage to avoid lodging.

### Rhizosphere soil samples

Rhizosphere soil samples were collected from 81 wheat LCs and the two elite cultivars following harvest. Composite samples were obtained by collecting three plants per plot and stored at –20 °C until analysis. The 81 LCs (Supplementary Table 2) were selected from the core set ^40^. The selection of the LCs was based on previous studies that reported differences among the LCs in the capability of their exudates to inhibit nitrification by *Nitrosomonas europaea* ^25^.

### Soil DNA extraction and bioinformatics used for the microbiome analysis

The total DNA from each composite rhizosphere soil sample was separately extracted using the FastDNA SPIN Kit for Soil (MP Biomedicals, CA, USA) according to the manufacturer’s instructions.

Amplicon sequencing was performed on the Illumina MiSeq platforms at Novogene Bioinformatics Technology Co., Ltd, Beijing, China, using primers 341F and 806R. Complete data sets are submitted to the NCBI Short Read Archive under accession no. PRJNA1243908.

Raw demultiplexed forward and reverse reads were processed using the methods implemented in QIIME2 version 2024.2.0 with default parameters unless otherwise stated ^61^. Deblur was used for quality filtering, denoising and calling Amplicon Sequence Variants (ASV’s) using the QIIME deblur denoise-paired method ^62^. Paired reads were trimmed to 250bp in length before denoising. The ASV’s were aligned and used to calculate a phylogenetic tree using the QIIME function phylogeny aligned to tree ^63,64^. Taxonomic assignment of the ASVs was performed using the QIIME naive Bayesian classifier Scikit-learn using the Greengenes2 full-length OTU database ^65–67^.

The raw sequencing data had a median of 14,380 reads per sample. Following filtering, denoising and merging, this was reduced to 5,504 reads per sample. A total of 3,600 ASV’s were identified across all the samples. Samples with fewer than 1,000 reads were excluded from subsequent analysis, and the data were rarefied to 2,430 reads. Picrust2 was used to predict metagenome functions from the 16S sequencing outputs ^68^. The pipeline was run with the default settings. KEGG modules were predicted from the Picrust2 output ^69,70^.

### Statistical analysis

Statistical analyses were conducted using R software version 4.3.2 ^71^.

The microbial diversity across the rhizosphere of the different genotypes was assessed by the Whittaker index alpha diversity, to determine the species richness for individual samples and beta diversity to estimate differences in composition between samples ^72^. Shannon’s, Chao1 and Inverse Simpson indexes (alpha diversity) and Evenness were calculated using the package phyloseq ^72^, both for individual accessions and the AGs. Faith’s phylogenetic diversity, which considers the evolutionary relationships of species within the rhizosphere community ^73^, and species richness, were determined using the R packages phyloseq and picante and the phylogenetic tree derived from the QIIME analysis. The association between alpha and phylogenetic diversities and the AGs were tested using the package microbiomeStat ^74^. Significant differences were calculated with the geom_pwc function for pairwise comparison from the ggpubr R package using the Wilcoxon test and the Benjamini-Hochberg method for the adjusted p-value.

The core microbiome was obtained using the microbiome R package for the top 50 taxa at the genus level. Prevalence is presented at increasing abundance thresholds across the wheat genotypes assessed. Sankey diagrams were built for the top 15 genera using an abundance threshold of 0.01. Additionally, a Sankey diagram was built after subsetting the key nitrifier taxa (Nitrososphaerales, Nitrospirales, Gemmatales) identified in the rhizosphere soil of the wheat genotypes.

The relationships between the elite cultivars and AGs based on their beta diversity were examined by multidimensional scaling (principal coordinates analysis) using the Bray-Curtis dissimilarity index, package microViz ^75^.

We used the R package pairwiseAdonis2 for multilevel pairwise comparison (PERMANOVA) to assess differences between the AG groups and the elite cultivars. The p-values were adjusted using the false discovery rate (FDR) method to control the number of errors associated with pairwise comparisons.

Differential abundance of the microbiome taxa between elite cultivars and AGs was calculated using linear regression for differential abundance analysis (LinDA) and visualised using volcano plots generated with the microbiomeStat R package ^74^.

Multivariate association between microbial composition and the AGs and elite cultivars was determined to identify differential abundances between the wheat groups using the MaAsLin2 package ^76^ and visualised using the geom_raster function of the ggplot2 R package.

An association network to model the interactions between microorganisms in the rhizosphere microbiome was constructed using the package NetCoMi ^58^. Taxa were aggregated at the order level. The network was constructed using a frequency threshold of 0.65, centred log ratio normalisation, the Pearson method to estimate the associations and dissimilarities and the cluster_fast_greedy clustering method. We then constructed a differential network to assess changes in microbiome interactions in response to the wheat genotypes using the Aitchison method and the hierarchical method for clustering. For these networks, we decided to represent microbiomes at the order level to provide a balance between taxonomic resolution and data manageability.

Next, we used the NetCoMi functions for network comparisons and construction of association and differential networks to compare network structure and metrics between the elite cultivars and each AG by selecting hubs with the highest degree and eigenvector centrality. For these analyses, taxa were aggregated at the genus level, and comparisons were performed using the Pearson and cluster_fast_greedy methods. Jaccard’s indices were calculated for the centrality measures, degree, betweenness, closeness, eigenvector and hub taxa with a default quantile threshold of 0.75. Keystone taxa were identified according to the highest degree and highest closeness centrality, and the lowest betweenness centrality scores ^77^. Visual comparison of the networks was performed using the “union” layout to facilitate the visual comparison of the networks by positioning nodes in a way that maximises their correspondence across the networks being compared. This helps identify shared nodes, edges, or other network structural elements. Node colours represent clusters. Differential networks were also constructed to highlight differences in association between nodes.

## Declaration

The authors declare no competing interests.

## Germplasm availability

The Watkins single seed-derived accessions and their predecessor landrace populations and elite cultivars are all available from the John Innes Centre Germplasm Resources Unit (https://www.seedstor.ac.uk/)

## Data availability

Datasets are available in the Supplementary Material. Complete sequencing data sets are submitted to the NCBI Short Read Archive under accession no.

PRJNA1243908.

## Acknowledgements

MCHS thanks the Daphne Jackson Trust and the Biotechnology and Biological Sciences Research Council (BBSRC) for a postdoctoral fellowship. We thank the germplasm resource unit at the John Innes Centre for providing the seeds. MCHS and AJM were funded by the BBSRC WISH-Roots 21EJP Soil: Tuning the wheat root microbiome to improve soil health and optimise rhizosphere nitrogen cycling and availability (grant no. BB/X003000/1). SG, FJW and LUW were funded by the BBSRC Institute Strategic Programme Delivering Sustainable Wheat (grant no.

BB/X011003/1). FJW gratefully acknowledges the support of the BBSRC; this research was funded by the BBSRC Institute Strategic Programme Food Microbiome and Health BB/X011054/1 and its constituent project BBS/E/F/000PR13631. AJM was funded by the BBSRC Institute Strategic Programmes: Harnessing Biosynthesis For Sustainable Food and Health (grant no. BB/X01097X/1) and Advancing Plant Health (grant no. BB/X010996/1). MCHS thanks Miss Maya Warren, Miss Amelia Lyons, Mr. Lewis Cocks and Miss Snowdrop Warren, for logistic support during the elaboration of this manuscript.

## Authors contributions

MCHS, FJW and LUW designed the experiment. MCHS carried out the experimental work. MCHS and FJW performed the data analysis. MCHS wrote the manuscript.

FH, LUW, AJM, FJW and SG reviewed the manuscript.

## References

1. Cheng, S. et al. Harnessing landrace diversity empowers wheat breeding. Nature 632, 823–831 (2024).

2. Davies, J. The business case for soil. Nature 543, 309–311 (2017).

3. Cappelli, S. L., Domeignoz-Horta, L. A., Loaiza, V. & Laine, A.-L. Plant biodiversity promotes sustainable agriculture directly and via belowground effects. Trends Plant Sci. 27, 674–687 (2022).

4. Gohar, S. et al. Domestication of newly evolved hexaploid wheat-A journey of wild grass to cultivated wheat. Front. Genet. 13, 1022931 (2022).

5. Lopes, M. S. et al. Exploiting genetic diversity from landraces in wheat breeding for adaptation to climate change. J. Exp. Bot. 66, 3477–3486 (2015).

6. Kavamura, V. N. et al. Wheat dwarfing influences selection of the rhizosphere microbiome. Sci. Rep. 10, 1452 (2020).

7. Weigelt, A. et al. An integrated framework of plant form and function: the belowground perspective. New Phytol. 232, 42–59 (2021).

8. Bardgett, R. D., Mommer, L. & De Vries, F. T. Going underground: root traits as drivers of ecosystem processes. Trends Ecol. Evol. 29, 692–699 (2014).

9. Freschet, G. T. et al. Root traits as drivers of plant and ecosystem functioning: current understanding, pitfalls and future research needs. New Phytol. (2020) doi:10.1111/nph.17072.

10. Pascale, A., Proietti, S., Pantelides, I. S. & Stringlis, I. A. Modulation of the root microbiome by plant molecules: The basis for targeted disease suppression and plant growth promotion. Front. Plant Sci. 10, 1741 (2019).

11. Santoyo, G. How plants recruit their microbiome? New insights into beneficial interactions. J. Adv. Res. 40, 45–58 (2022).

12. Toyota, K. & Shirai, S. Growing interest in microbiome research unraveling disease suppressive soils against plant pathogens. Microbes Environ. 33, 345–347 (2018).

13. Ling, N., Wang, T. & Kuzyakov, Y. Rhizosphere bacteriome structure and functions. Nat. Commun. 13, 836 (2022).

14. Eichmann, R., Richards, L. & Schäfer, P. Hormones as go-betweens in plant microbiome assembly. Plant J. 105, 518–541 (2021).

15. Trivedi, P., Leach, J. E., Tringe, S. G., Sa, T. & Singh, B. K. Plant-microbiome interactions: from community assembly to plant health. Nat. Rev. Microbiol. 18, 607–621 (2020).

16. Clouse, K. M. & Wagner, M. R. Plant Genetics as a Tool for Manipulating Crop Microbiomes: Opportunities and Challenges. Front Bioeng Biotechnol 9, 567548 (2021).

17. Xun, W. et al. Dissection of rhizosphere microbiome and exploiting strategies for sustainable agriculture. New Phytol. (2024) doi:10.1111/nph.19697.

18. Coyte, K. Z., Schluter, J. & Foster, K. R. The ecology of the microbiome: Networks, competition, and stability. Science 350, 663–666 (2015).

19. Bradshaw, A. Evolutionary significance of phenotypic plasticity in plants. Advances in Genetics 13, 115–155 (1965).

20. Wang, X., Zhang, J., Lu, X., Bai, Y. & Wang, G. Two diversities meet in the rhizosphere: root specialized metabolites and microbiome. J. Genet. Genomics 51, 467–478 (2024).

21. Trivedi, P., Mattupalli, C., Eversole, K. & Leach, J. E. Enabling sustainable agriculture through understanding and enhancement of microbiomes. New Phytol. 230, 2129–2147 (2021).

22. Baggs, E. M. et al. Exploiting crop genotype-specific root-soil interactions to enhance agronomic efficiency. *Front*. Soil Sci. 3, (2023).

23. Langholtz, M. et al. Increased nitrogen use efficiency in crop production can provide economic and environmental benefits. Sci. Total Environ. 758, 143602 (2021).

24. Coskun, D., Britto, D. T., Shi, W. & Kronzucker, H. J. Nitrogen transformations in modern agriculture and the role of biological nitrification inhibition. Nature Plants 3, 1–10 (2017).

25. O’Sullivan, C. A., Fillery, I. R. P., Roper, M. M. & Richards, R. A. Identification of several wheat landraces with biological nitrification inhibition capacity. Plant Soil 404, 61–74 (2016).

26. Bozal-Leorri, A. et al. Biological nitrification inhibitor-trait enhances nitrogen uptake by suppressing nitrifier activity and improves ammonium assimilation in two elite wheat varieties. Front. Plant Sci. 13, 1034219 (2022).

27. Kuypers, M. M. M., Marchant, H. K. & Kartal, B. The microbial nitrogen-cycling network. Nat. Rev. Microbiol. 16, 263–276 (2018).

28. Daims, H., Lücker, S. & Wagner, M. A new perspective on microbes formerly known as nitrite-oxidizing bacteria. Trends Microbiol. 24, 699–712 (2016).

29. Ghatak, A., Chaturvedi, P., Waldherr, S., Subbarao, G. V. & Weckwerth, W. PANOMICS at the interface of root-soil microbiome and BNI. Trends Plant Sci. 28, 106–122 (2023).

30. Poirier, V., Roumet, C. & Munson, A. D. The root of the matter: Linking root traits and soil organic matter stabilization processes. Soil Biol. Biochem. 120, 246–259 (2018).

31. Shabtai, I. A. et al. Root exudates simultaneously form and disrupt soil organo-mineral associations. Commun. Earth Environ. 5, 1–12 (2024).

32. Panchal, P., Preece, C., Peñuelas, J. & Giri, J. Soil carbon sequestration by root exudates. Trends Plant Sci. 27, 749–757 (2022).

33. Hawkesford, M. J. Reducing the reliance on nitrogen fertilizer for wheat production. J. Cereal Sci. 59, 276–283 (2014).

34. Fatholahi, S., Ehsanzadeh, P. & Karimmojeni, H. Ancient and improved wheats are discrepant in nitrogen uptake, remobilization, and use efficiency yet comparable in nitrogen assimilating enzymes capabilities. Field Crops Res. 249, 107761 (2020).

35. Cormier, F. et al. Breeding for increased nitrogen-use efficiency: a review for wheat (*T*. aestivumL.). Plant Breed. 135, 255–278 (2016).

36. Shewry, P. R. et al. Do modern types of wheat have lower quality for human health? Nutr. Bull. 45, 362–373 (2020).

37. Gruet, C., Muller, D. & Moënne-Loccoz, Y. Significance of the diversification of wheat species for the assembly and functioning of the root-associated microbiome. Front. Microbiol. 12, 782135 (2021).

38. Yue, H. et al. Plant domestication shapes rhizosphere microbiome assembly and metabolic functions. Microbiome 11, 70 (2023).

39. Luo, C., He, Y. & Chen, Y. Rhizosphere microbiome regulation: Unlocking the potential for plant growth. Curr. Res. Microb. Sci. 8, 100322 (2025).

40. Wingen, L. U. et al. Establishing the AE Watkins landrace cultivar collection as a resource for systematic gene discovery in bread wheat. Theor. Appl. Genet. 127, 1831–1842 (2014).

41. Kumar, A. et al. Shifts in plant functional trait dynamics in relation to soil microbiome in modern and wild barley. Plants People Planet (2024) doi:10.1002/ppp3.10534.

42. Ndour, P. M. S. et al. Pearl millet genotype impacts microbial diversity and enzymatic activities in relation to root-adhering soil aggregation. Plant Soil 464, 109–129 (2021).

43. Xiong, J. et al. Effect of rice (Oryza sativa L.) genotype on yield: Evidence from recruiting spatially consistent rhizosphere microbiome. Soil Biol. Biochem. 161, 108395 (2021).

44. Shenton, M., Iwamoto, C., Kurata, N. & Ikeo, K. Effect of wild and cultivated rice genotypes on rhizosphere bacterial community composition. Rice (N. Y.) 9, 42 (2016).

45. Huang, J. et al. The rhizospheric microbiome becomes more diverse with maize domestication and genetic improvement. J. Integr. Agric. 21, 1188–1202 (2022).

46. Barnes, C. J., Bahram, M., Nicolaisen, M., Gilbert, M. T. P. & Vestergård, M. Microbiome selection and evolution within wild and domesticated plants. Trends Microbiol. 33, 447–458 (2025).

47. Banerjee, S. et al. Agricultural intensification reduces microbial network complexity and the abundance of keystone taxa in roots. ISME J. 13, 1722–1736 (2019).

48. Yang, X., Hu, H.-W., Yang, G.-W., Cui, Z.-L. & Chen, Y.-L. Crop rotational diversity enhances soil microbiome network complexity and multifunctionality. Geoderma 436, 116562 (2023).

49. Simonin, M. et al. Influence of plant genotype and soil on the wheat rhizosphere microbiome: evidences for a core microbiome across eight African and European soils. FEMS Microbiol. Ecol. 96, (2020).

50. Zhou, Y. et al. The preceding root system drives the composition and function of the rhizosphere microbiome. Genome Biol. 21, 89 (2020).

51. Jaiswal, S. et al. Unveiling the wheat microbiome under varied agricultural field conditions. Microbiol. Spectr. 10, e0263322 (2022).

52. Prosser, J. I., Hink, L., Gubry-Rangin, C. & Nicol, G. W. Nitrous oxide production by ammonia oxidizers: Physiological diversity, niche differentiation and potential mitigation strategies. Glob. Chang. Biol. 26, 103–118 (2020).

53. Nelkner, J. et al. Abundance, classification and genetic potential of Thaumarchaeota in metagenomes of European agricultural soils: a meta-analysis. Environ Microbiome 18, 26 (2023).

54. Balfourier, F. et al. Worldwide phylogeography and history of wheat genetic diversity. Sci. Adv. 5, eaav0536 (2019).

55. L’Espérance, E., Bouyoucef, L. S., Dozois, J. A. & Yergeau, E. Tipping the plant-microbe competition for nitrogen in agricultural soils. iScience 27, 110973 (2024).

56. Kajihara, K. T. & Hynson, N. A. Networks as tools for defining emergent properties of microbiomes and their stability. Microbiome 12, 184 (2024).

57. Gao, C. et al. Co-occurrence networks reveal more complexity than community composition in resistance and resilience of microbial communities. Nat. Commun. 13, 3867 (2022).

58. Peschel, S., Müller, C. L., von Mutius, E., Boulesteix, A.-L. & Depner, M. NetCoMi: network construction and comparison for microbiome data in R. Brief. Bioinform. 22, bbaa290 (2021).

59. Banerjee, S. & van der Heijden, M. G. A. Soil microbiomes and one health. Nat. Rev. Microbiol. 21, 6–20 (2023).

60. Issifu, S., Acharya, P., Kaur-Bhambra, J., Gubry-Rangin, C. & Rasche, F. Biological nitrification inhibitors with antagonistic and synergistic effects on growth of ammonia oxidisers and soil nitrification. Microb. Ecol. 87, 143 (2024).

61. Bolyen, E. et al. Reproducible, interactive, scalable and extensible microbiome data science using QIIME 2. Nature biotechnology 37, 852–857 (2019).

62. Amir, A., et al. Deblur rapidly resolves single-nucleotide community sequence patterns. mSystems 2, (2017).

63. Katoh, K. & Standley, D. M. MAFFT multiple sequence alignment software version 7: improvements in performance and usability. Mol. Biol. Evol. 30, 772–780 (2013).

64. Price, M. N., Dehal, P. S. & Arkin, A. P. FastTree: computing large minimum evolution trees with profiles instead of a distance matrix. Mol. Biol. Evol. 26, 1641–1650 (2009).

65. McDonald, D. et al. Greengenes2 unifies microbial data in a single reference tree. Nat. Biotechnol. 42, 715–718 (2024).

66. Pedregosa, F. et al. Scikit-learn: Machine learning in Python. the Journal of machine Learning research 12, 2825–2830 (2011).

67. Bokulich, N. A. et al. Optimizing taxonomic classification of marker-gene amplicon sequences with QIIME 2’s q2-feature-classifier plugin. Microbiome 6, 90 (2018).

68. Douglas, G. M. et al. PICRUSt2 for prediction of metagenome functions. Nat. Biotechnol. 38, 685–688 (2020).

69. Kanehisa, M., Sato, Y., Kawashima, M., Furumichi, M. & Tanabe, M. KEGG as a reference resource for gene and protein annotation. Nucleic Acids Res. 44, D457–62 (2016).

70. Kanehisa, M., Sato, Y. & Morishima, K. BlastKOALA and GhostKOALA: KEGG tools for functional characterization of genome and metagenome sequences. J. Mol. Biol. 428, 726–731 (2016).

71. Core Team, R. A language and environment for statistical computing. *R Found*. Stat. Comput. (2018).

72. McMurdie, P. J. & Holmes, S. phyloseq: an R package for reproducible interactive analysis and graphics of microbiome census data. PLoS One 8, e61217 (2013).

73. Faith, D., Reid, C. & Hunter, J. Integrating phylogenetic diversity, complementarity, and endemism for conservation assessment. Conservation Biology 18, 255–261 (2004).

74. Zhou, H., He, K., Chen, J. & Zhang, X. LinDA: linear models for differential abundance analysis of microbiome compositional data. Genome Biol. 23, 95 (2022).

75. Barnett, D., Arts, I. & Penders, J. microViz: an R package for microbiome data visualization and statistics. J. Open Source Softw. 6, 3201 (2021).

76. Mallick, H. et al. Multivariable association discovery in population-scale meta-omics studies. PLoS Comput. Biol. 17, e1009442 (2021).

77. Berry, D. & Widder, S. Deciphering microbial interactions and detecting keystone species with co-occurrence networks. Front. Microbiol. 5, 219 (2014).

