## Supplementary Figures and Supplementary Tables Captions for "Unearthing the rhizosphere microbiome recruited by ancestral bread wheat landraces"

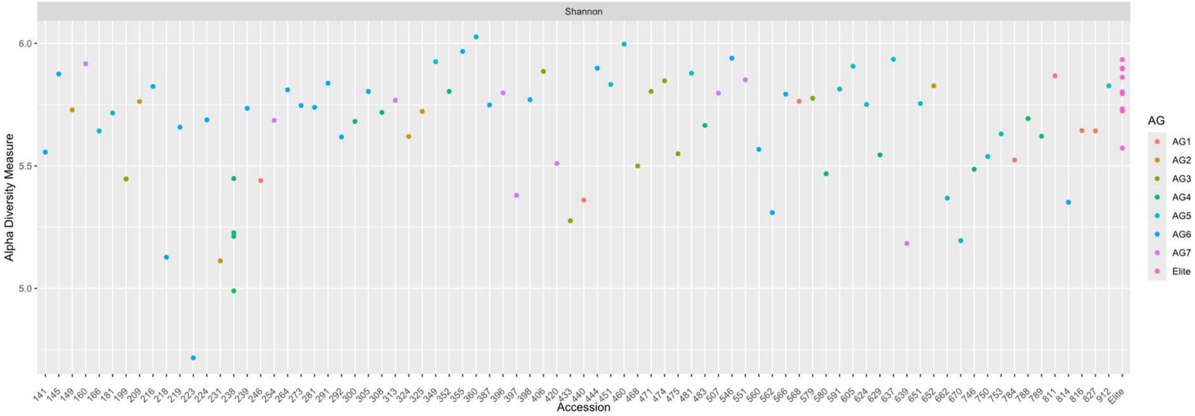


**Figure S1**. Alpha-diversity (Shannon index) across the rhizosphere soil of 83 genotypes of hexaploid wheat (two elite cultivars and 81 landraces). Colour code for the classification of the wheat genotypes into ancestral groups (AG) [2], closely related genetically, and elite (modern) cultivars. Shannon indices were calculated with the phyloseq [6] R package. Values are provided in Supplementary Table 4.

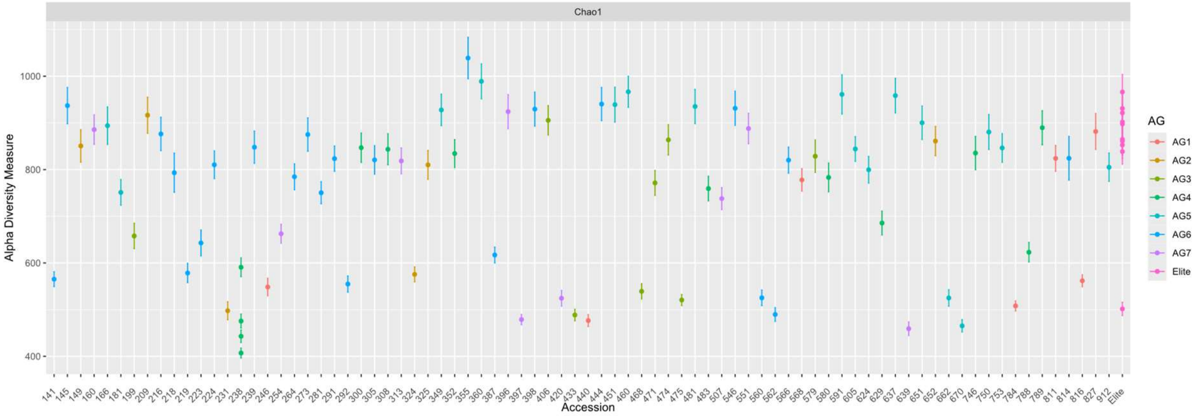


**Figure S2**. Alpha-diversity (Chao1 index) across the rhizosphere soil of 83 genotypes of hexaploid wheat (two elite cultivars and 81 landraces). Colour code for the classification of the wheat genotypes into ancestral groups (AG) [2], closely related genetically, and elite (modern) cultivars. Chao1 indices were calculated with the phyloseq [6] R package. Values and standard error of Chao's estimator of the number of unseen species are provided in Supplementary Table 4.

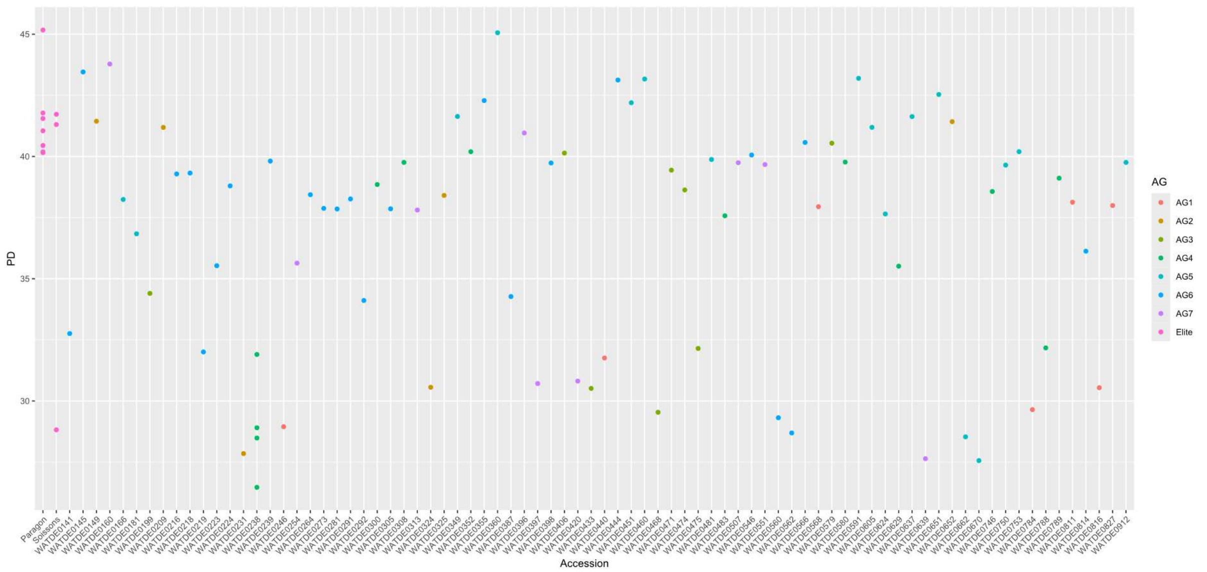


**Figure S3**. Faith's phylogenetic diversity (PD) across the rhizosphere soil of 83 genotypes of hexaploid wheat (two elite cultivars and 81 landraces). Colour code for the classification of the wheat genotypes into ancestral groups (AG) [2], closely related genetically, and elite (modern) cultivars. Faith's indices were calculated using the R packages phyloseq [6] and picante [7]. Values are provided in Supplementary Table 4.

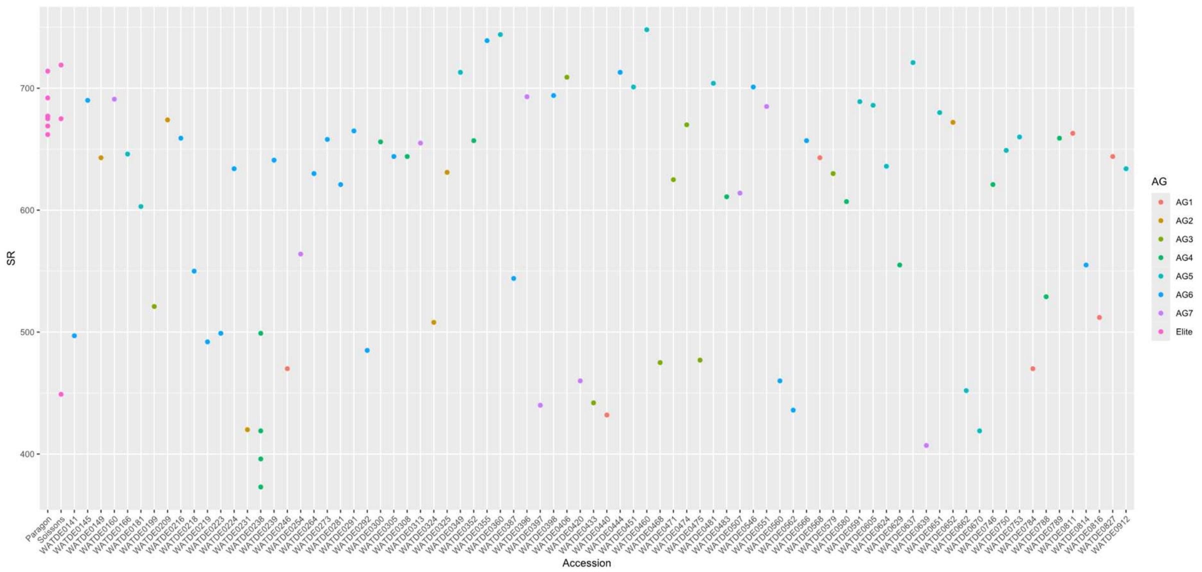


**Figure S4**. Species richness (SR) across the rhizosphere soil of 83 genotypes of hexaploid wheat (two elite cultivars and 81 landraces). Colour code for the classification of the wheat genotypes into ancestral groups (AG) [2], closely related genetically, and elite (modern) cultivars. The SR was calculated using the R packages phyloseq [6] and picante [7]. Values are provided in Supplementary Table 4.

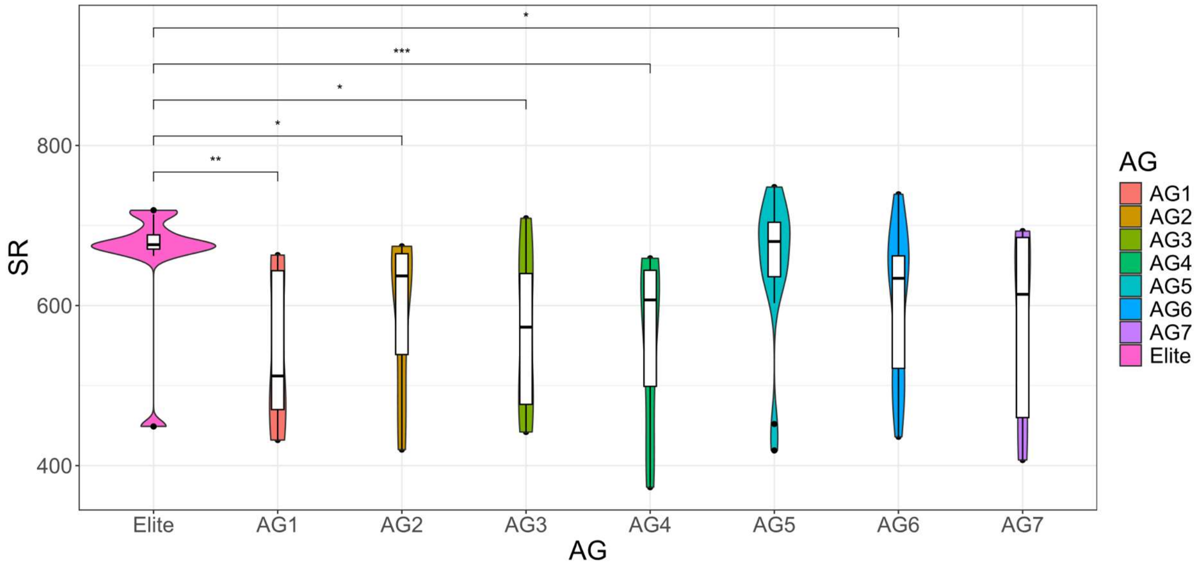


**Figure S5**. Species richness (SR) across 81 wheat genotypes grouped into ancestral groups (AG) [2], closely related genetically, and two elite (modern) cultivars. Significant differences in Shannon diversity and PD were calculated by pairwise comparison using the Wilcoxon test and the Benjamini-Hochberg method (R packages phyloseq [6], ggpubr [8] and ggplot2 [9]) for the adjusted p-value (*p ≤ 0.05, **p ≤ 0.01, ***p ≤ 0.001).

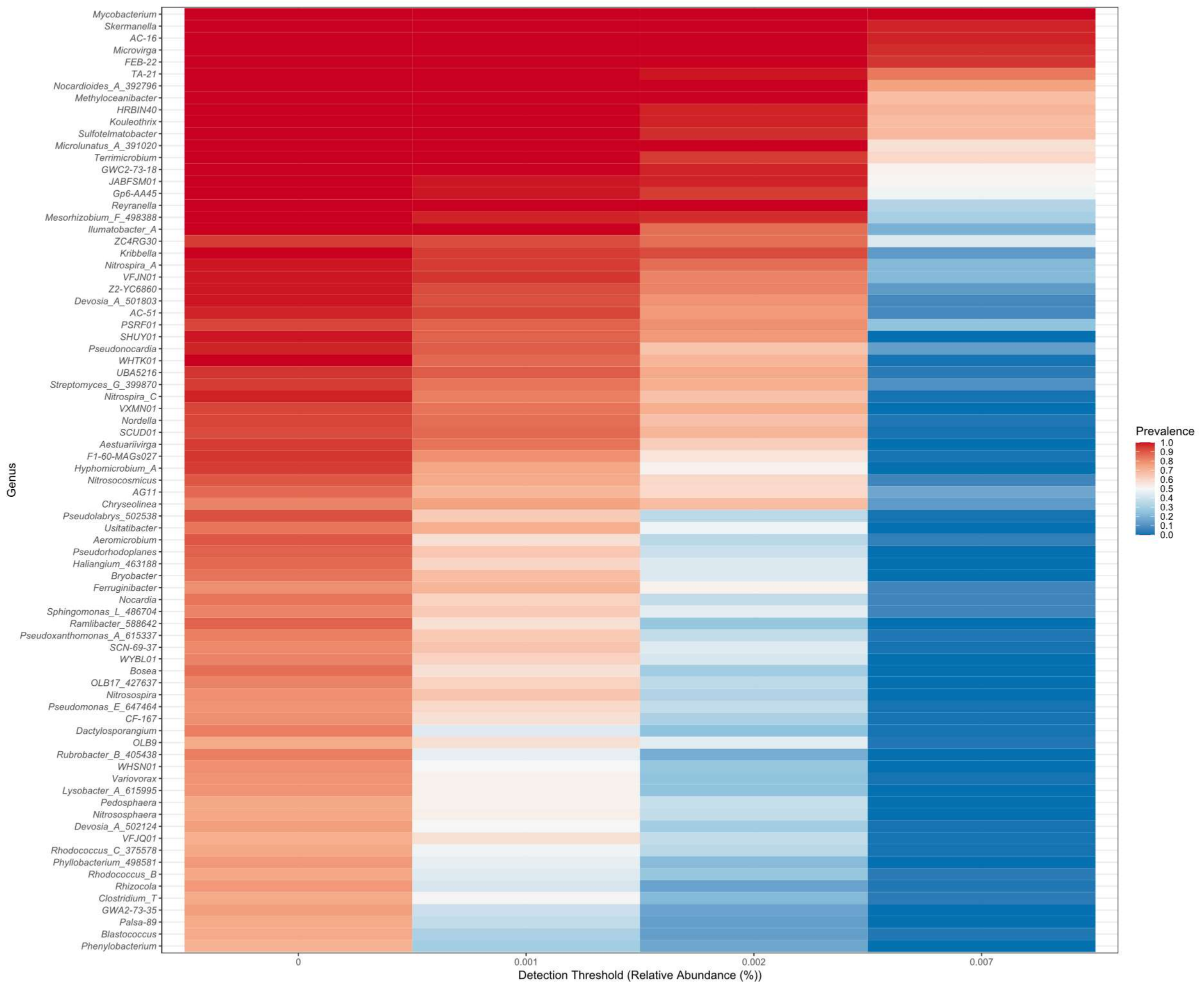


**Figure S6**. Core microbiome analysis based on relative abundance and sample prevalence of microbial taxa grouped by genus, from rhizosphere soil collected from 83 genotypes of hexaploid wheat (two elite cultivars and 81 landraces). Relative abundance was obtained using the R package microbiome [10] (transformation “compositional”).

**Figure S7**. Volcano plots showing differential abundance of microbial taxa at phylum (Supplementary Table 6), order (Supplementary Table 7) and genus level (Supplementary Table 8) between 81 wheat genotypes categorised into ancestral groups (AG) [2], closely related genetically, and two elite (modern) cultivars. Linear models for differential abundance analysis (LinDA) were obtained using the R package MicrobiomeStat [4].

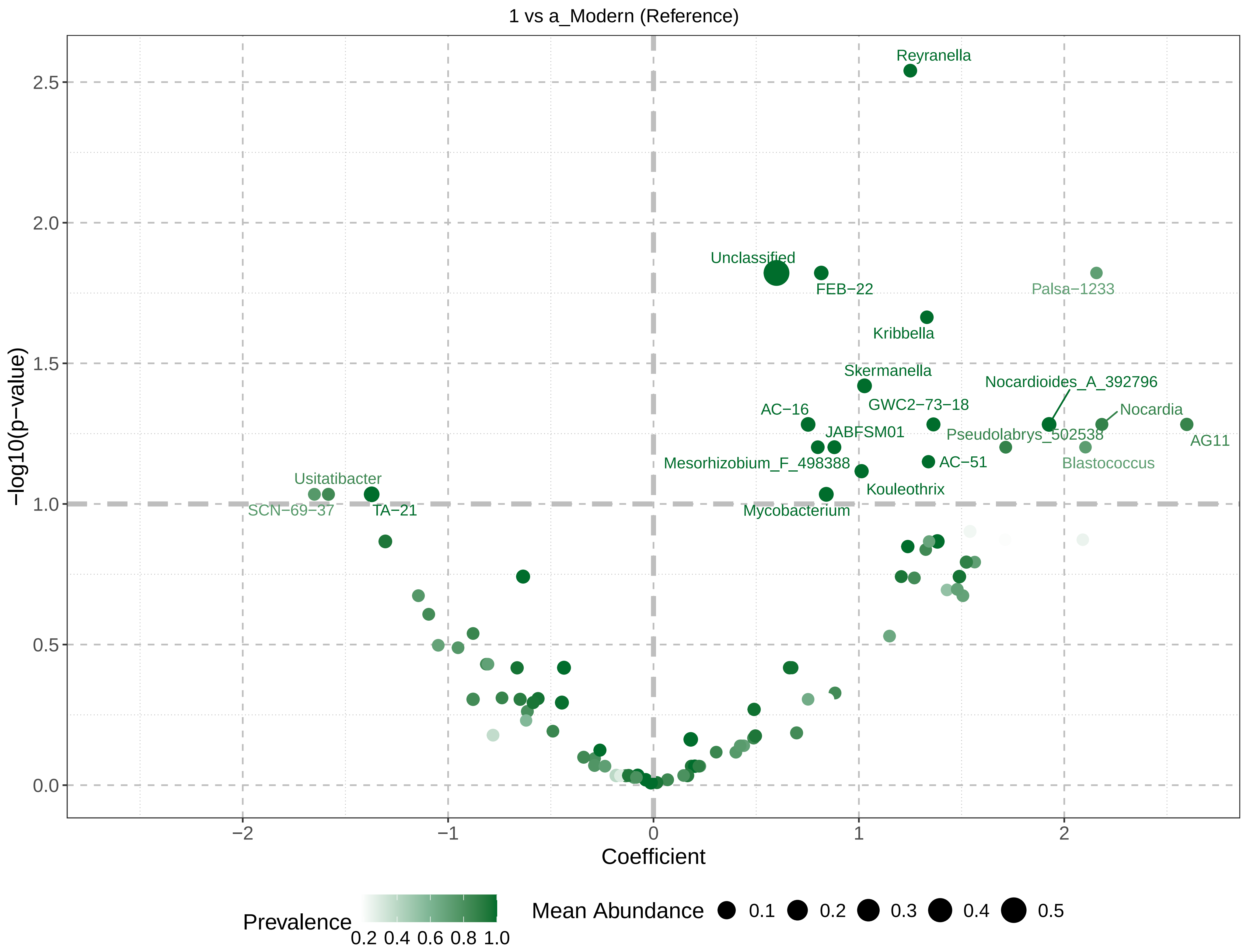
1 vs a_Modern (Reference)
Reyranella
Unclassified
FEB−22
Palsa−1233
Kribbella
Skermanella
Nocardioides_A_392796
GWC2−73−18
AC−16
Nocardia
JABFSM01
Pseudolabrys_502538
AG11
Mesorhizobium_F_498388
AC−51
Blastococcus
Usitatibacter
Kouleothrix
SCN−69−37
TA−21
Mycobacterium
−log10(p−value)
Coefficient
Prevalence
0.2 0.4 0.6 0.8 1.0
Mean Abundance
0.1
0.2
0.3
0.4
0.5

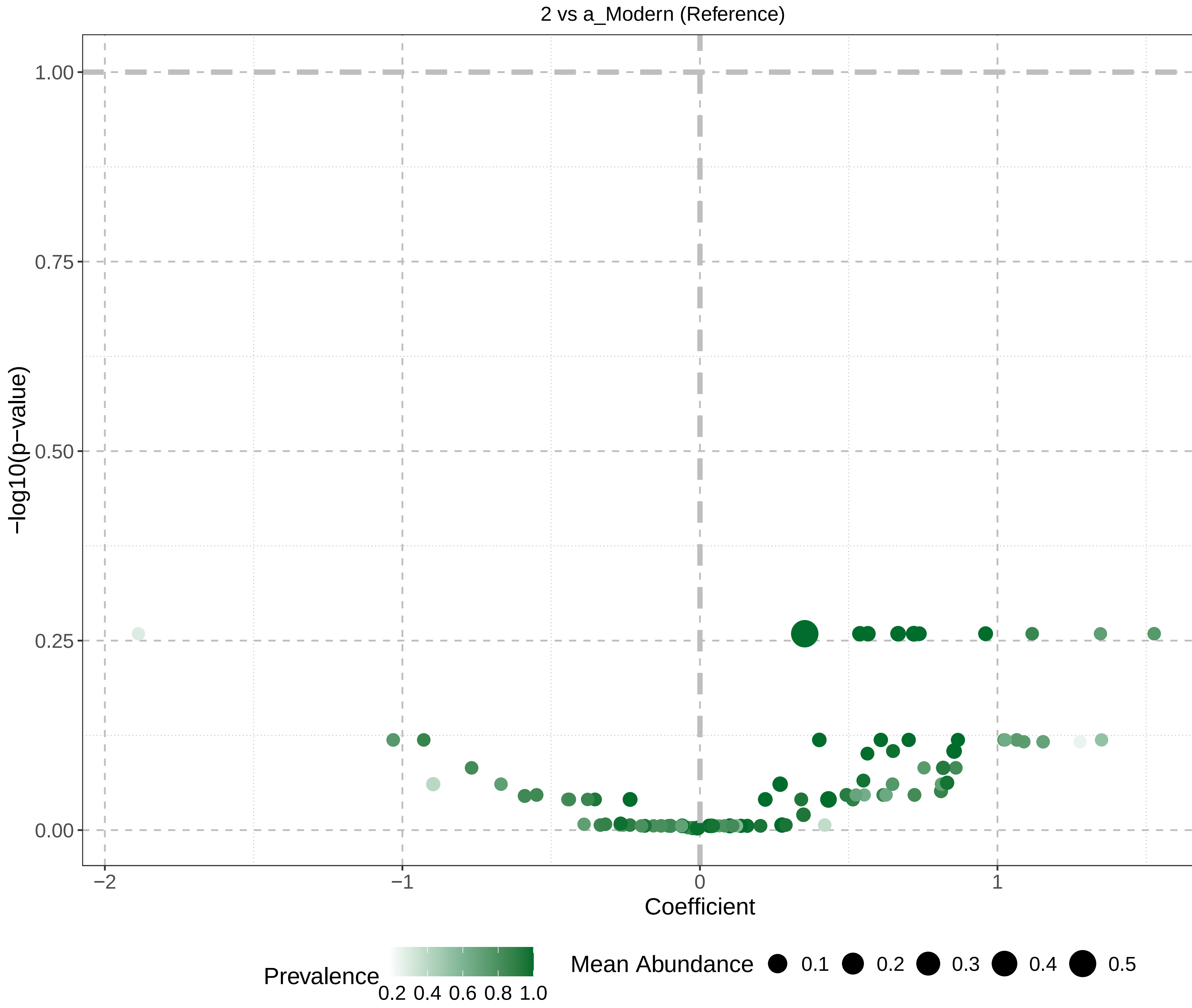

2 vs a_Modern (Reference)
−log10(p−value)
1.00
0.75
0.50
0.25
0.00
−2
−1
0
1
2
Coefficient
Prevalence
0.2 0.4 0.6 0.8 1.0
Mean Abundance
0.1
0.2
0.3
0.4
0.5

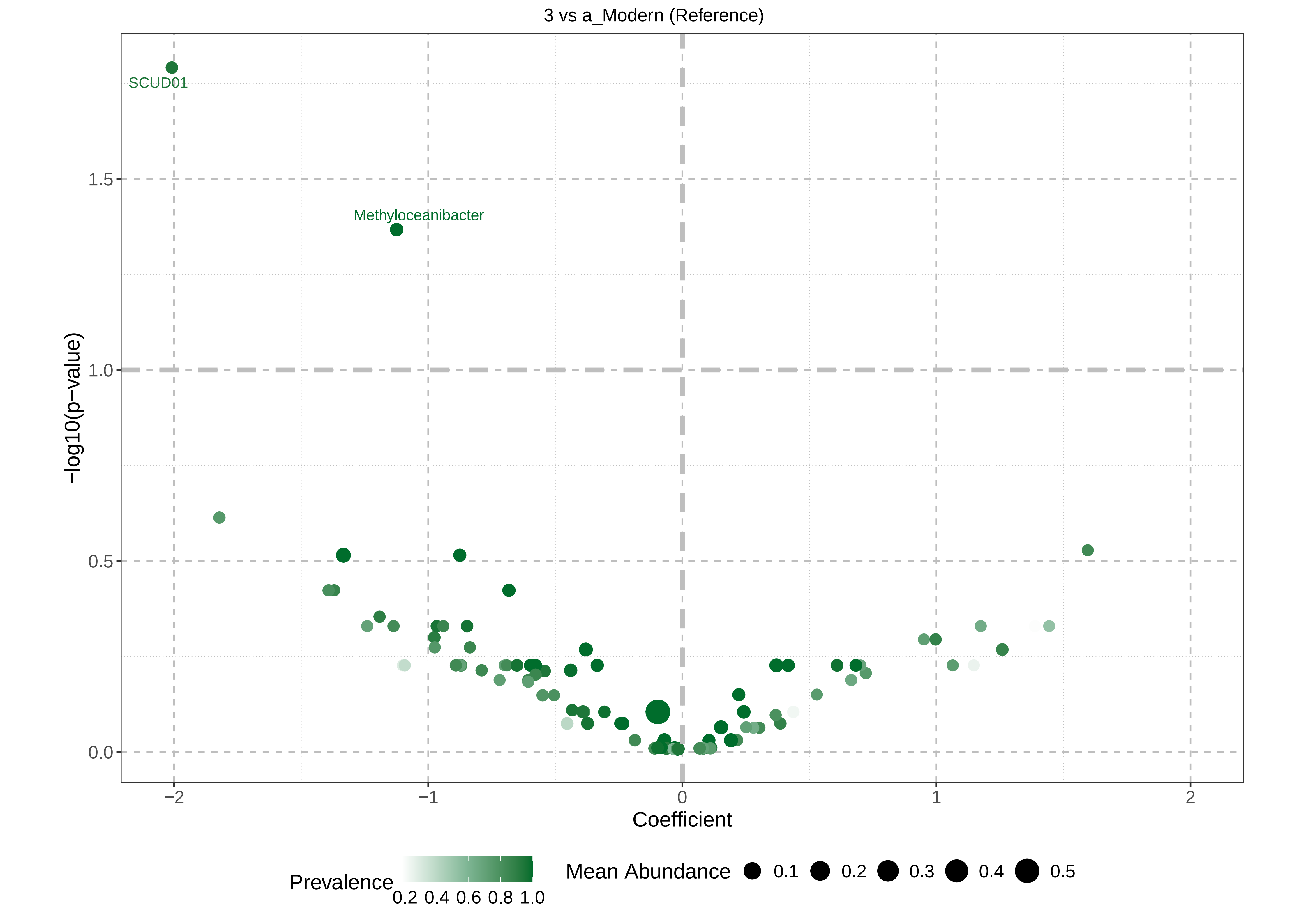

3 vs a_Modern (Reference)
SCUD01
Methyloceanibacter
−log10(p−value)
1.5
1.0
0.5
0.0
−2
−1
0
1
2
Coefficient
Prevalence
0.2 0.4 0.6 0.8 1.0
Mean Abundance
0.1
0.2
0.3
0.4
0.5

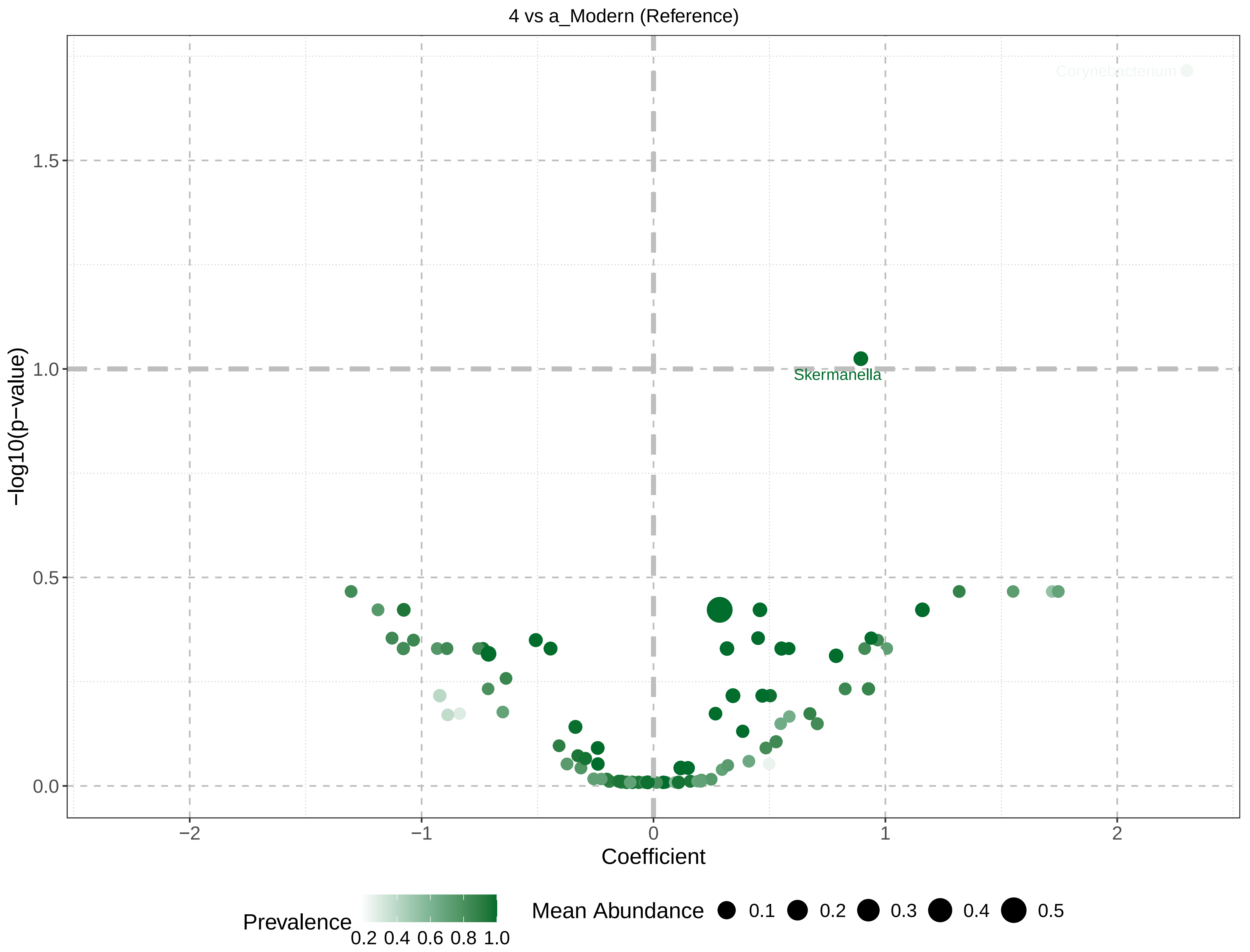

4 vs a_Modern (Reference)
Corynebacterium
Skermanella
1.5
1.0
0.5
0.0
−log10(p−value)
−2
−1
0
1
2
Coefficient
Prevalence
0.2 0.4 0.6 0.8 1.0
Mean Abundance
0.1
0.2
0.3
0.4
0.5

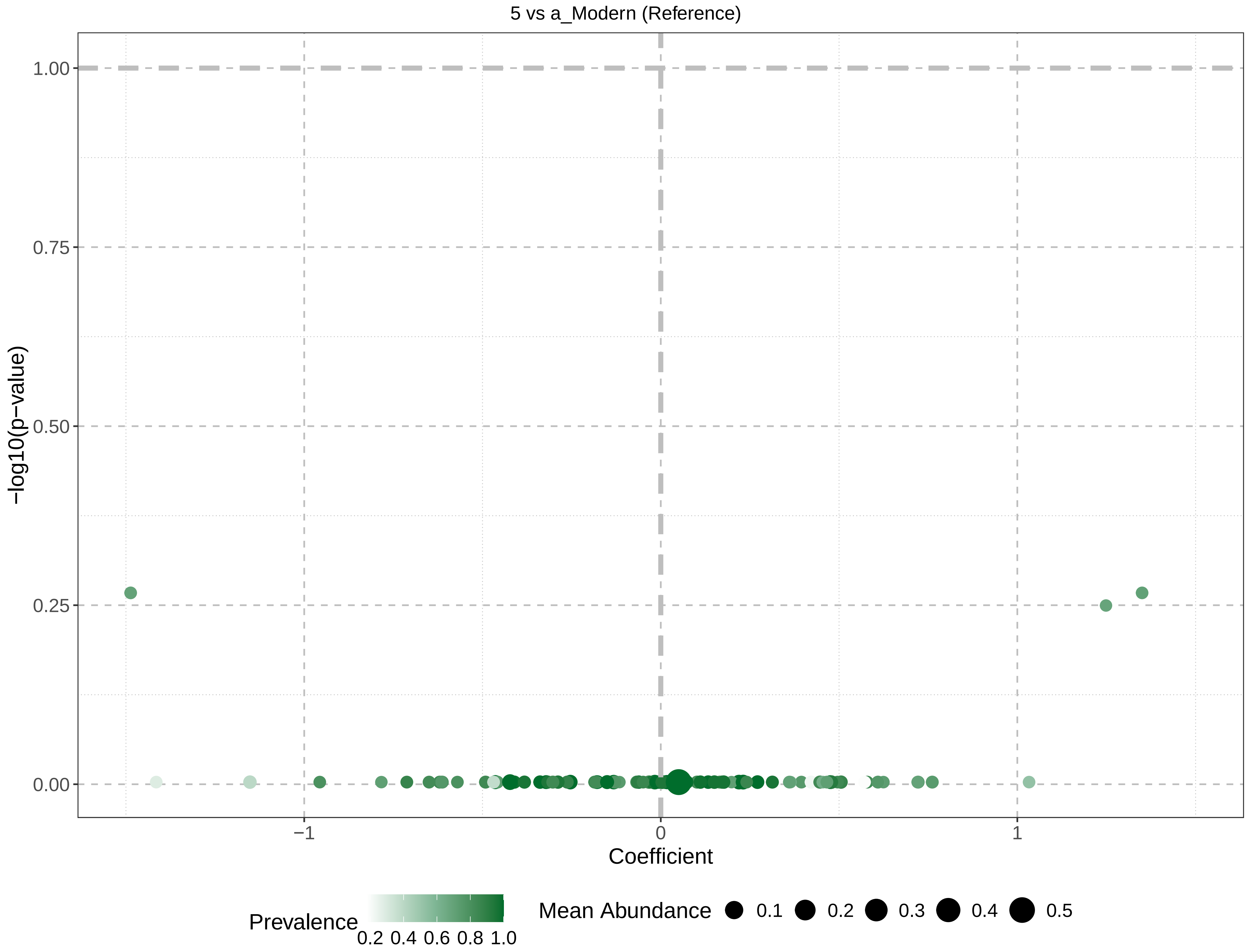

5 vs a_Modern (Reference)
−log10(p−value)
1.00
0.75
0.50
0.25
0.00
−1
0
1
Coefficient
Prevalence
0.2 0.4 0.6 0.8 1.0
Mean Abundance
0.1
0.2
0.3
0.4
0.5

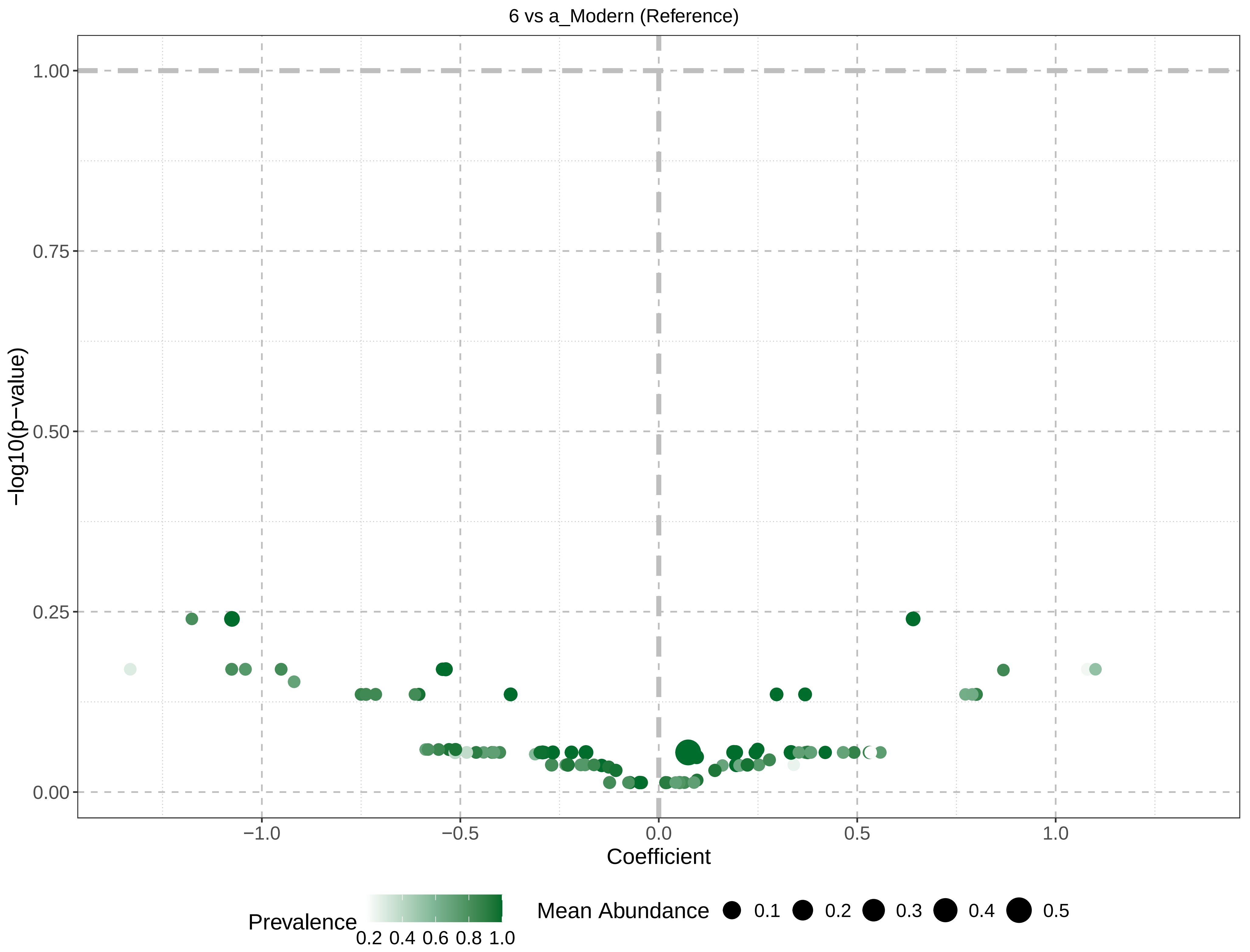

6 vs a_Modern (Reference)
1.00
0.75
0.50
0.25
0.00
−log10(p−value)
−1.0
−0.5
0.0
0.5
1.0
Coefficient
Prevalence
0.2 0.4 0.6 0.8 1.0
Mean Abundance
0.1
0.2
0.3
0.4
0.5

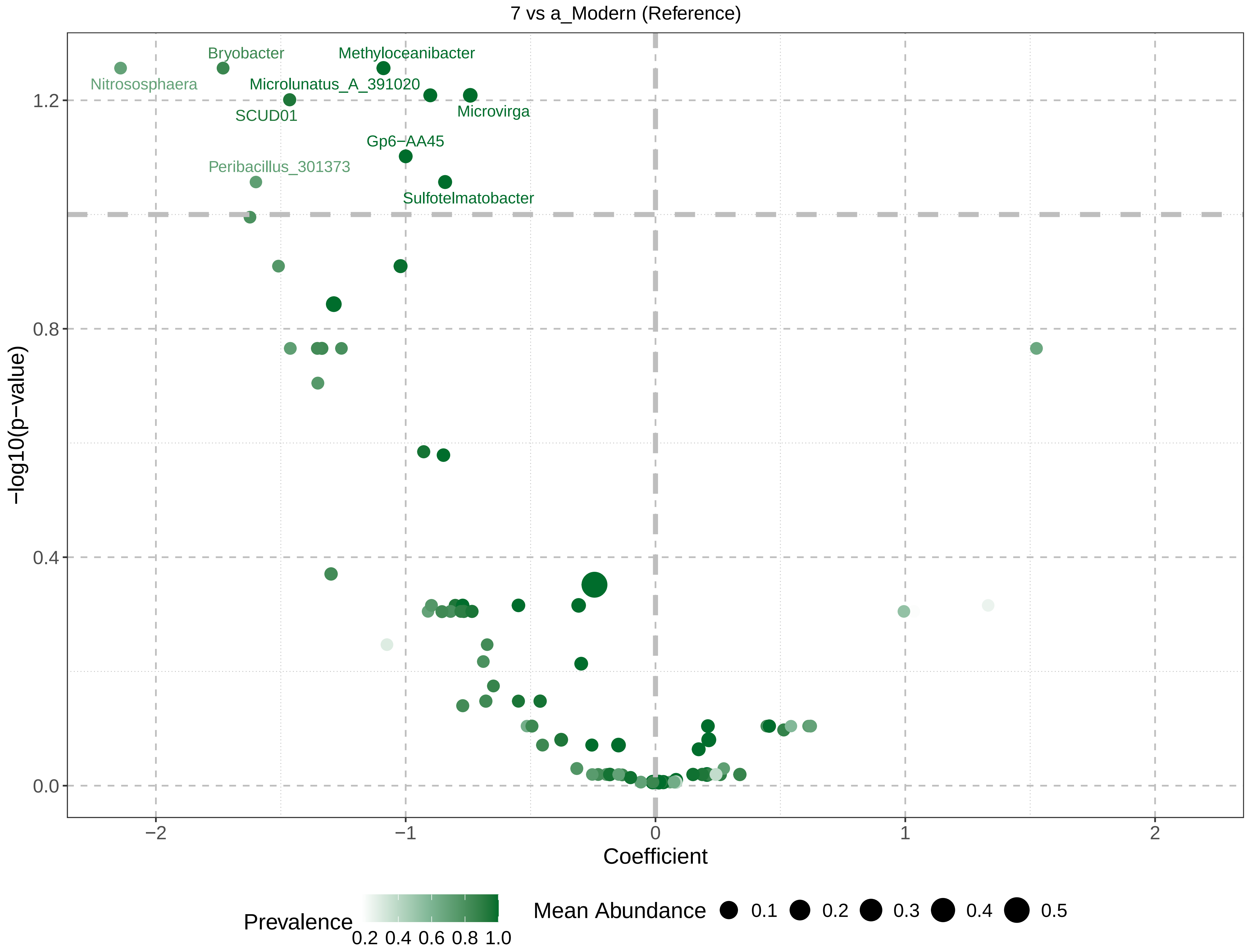
7 vs a_Modern (Reference)
Nitrososphaera
Bryobacter
Methyloceanibacter
Microlunatus_A_391020
SCUD01
Microvirga
Gp6−AA45
Peribacillus_301373
Sulfotelmatobacter
−log10(p−value)
1.2
0.8
0.4
0.0
−2
−1
0
1
2
Coefficient
Prevalence
0.2 0.4 0.6 0.8 1.0
Mean Abundance
0.1
0.2
0.3
0.4
0.5

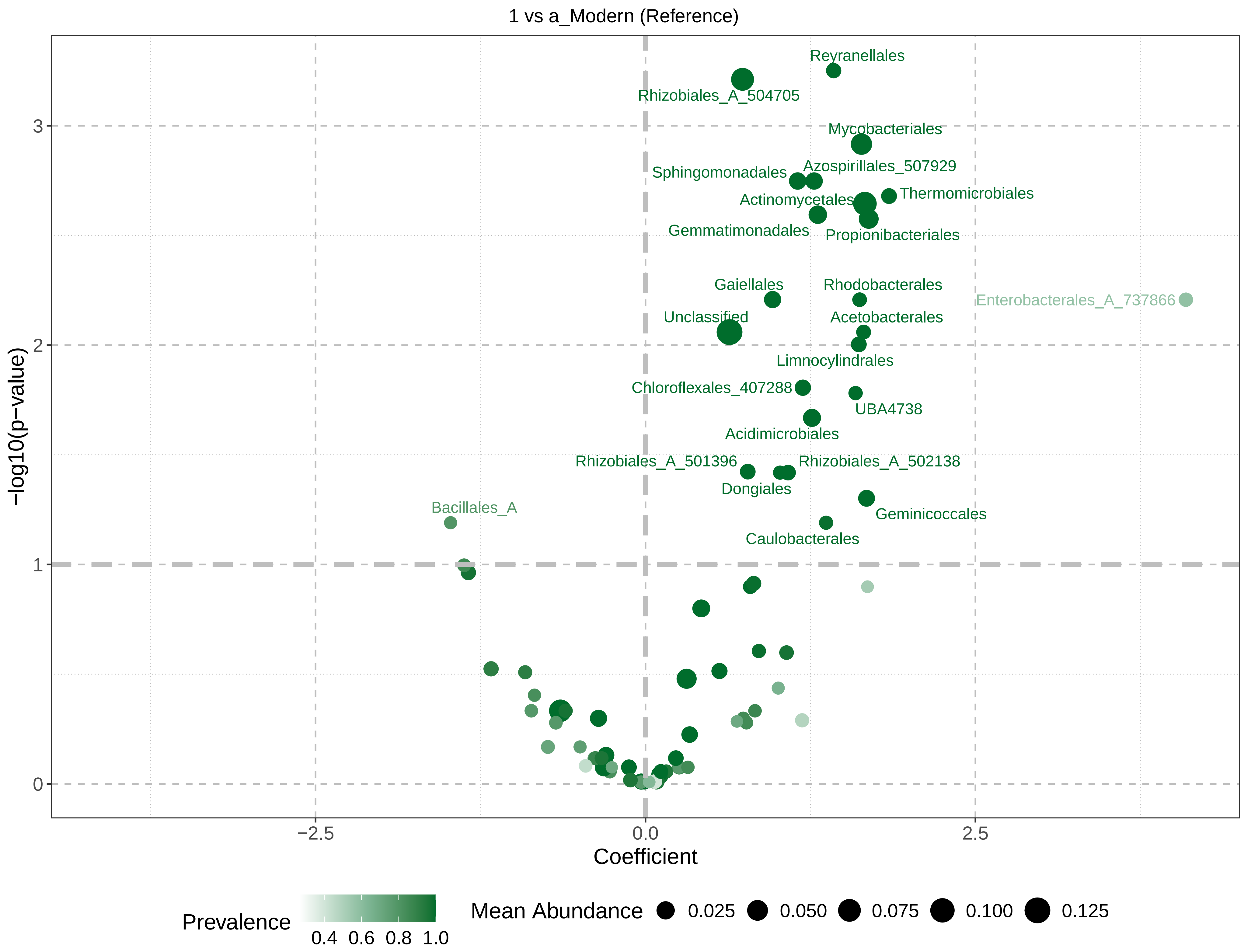

1 vs a_Modern (Reference)
Reyranellales
Rhizobiales_A_504705
Mycobacteriales
Sphingomonadales
Azospirillales_507929
Thermomicrobiales
Actinomycetales
Gemmatimonadales
Propionibacteriales
Gaiellales
Rhodobacterales
Enterobacterales_A_737866
Unclassified
Acetobacterales
Limnocylindrales
Chloroflexales_407288
UBA4738
Acidimicrobiales
Rhizobiales_A_501396
Rhizobiales_A_502138
Dongiales
Bacillales_A
Geminicoccales
Caulobacterales
−log10(p−value)
0
1
2
3
−2.5
0.0
2.5
Coefficient
Prevalence
0.4 0.6 0.8 1.0
Mean Abundance
0.025
0.050
0.075
0.100
0.125

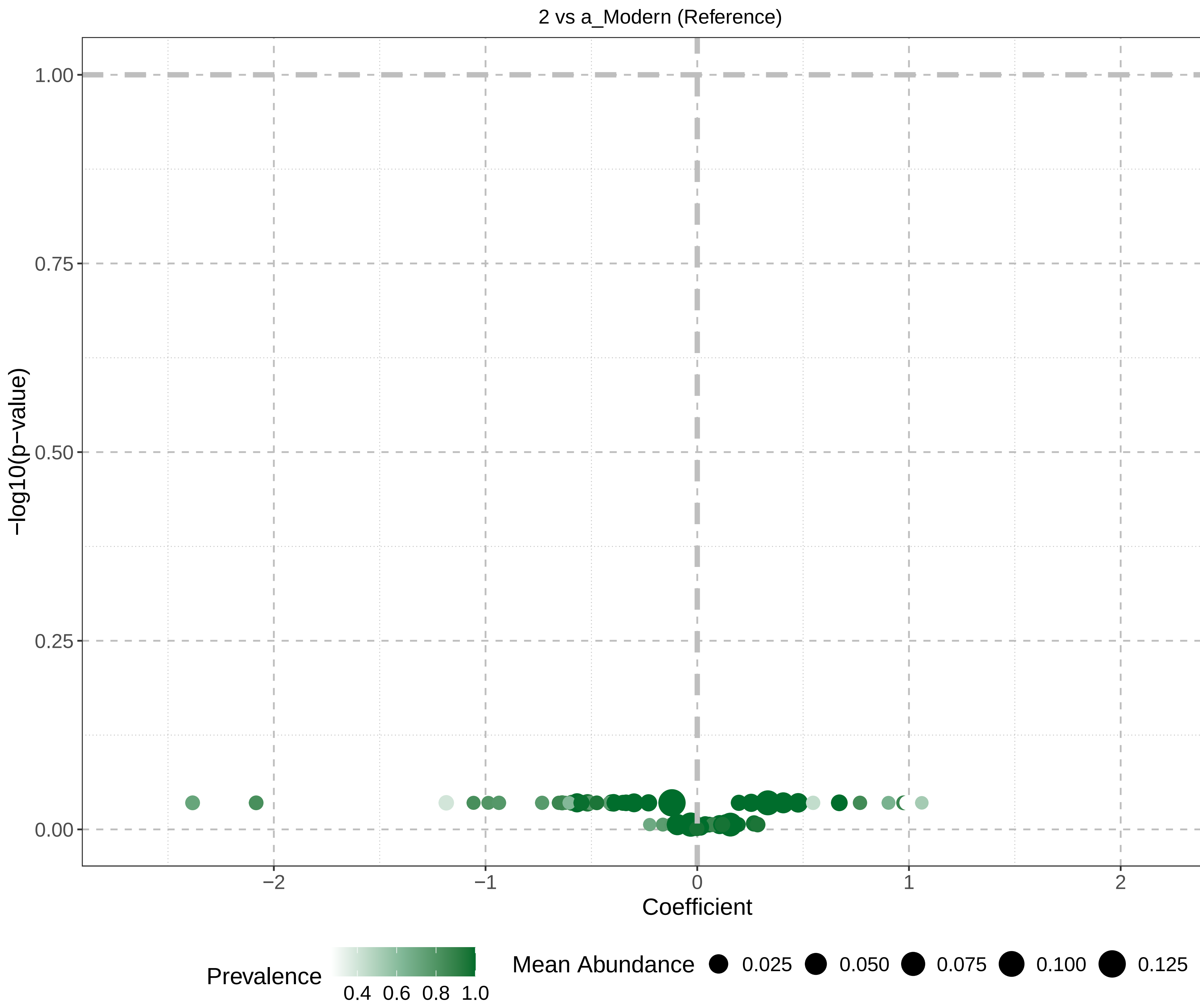
2 vs a_Modern (Reference)
1.00
0.75
0.50
0.25
0.00
−log10(p−value)
−2
−1
0
1
2
Coefficient
Prevalence
0.4 0.6 0.8 1.0
Mean Abundance
0.025
0.050
0.075
0.100
0.125

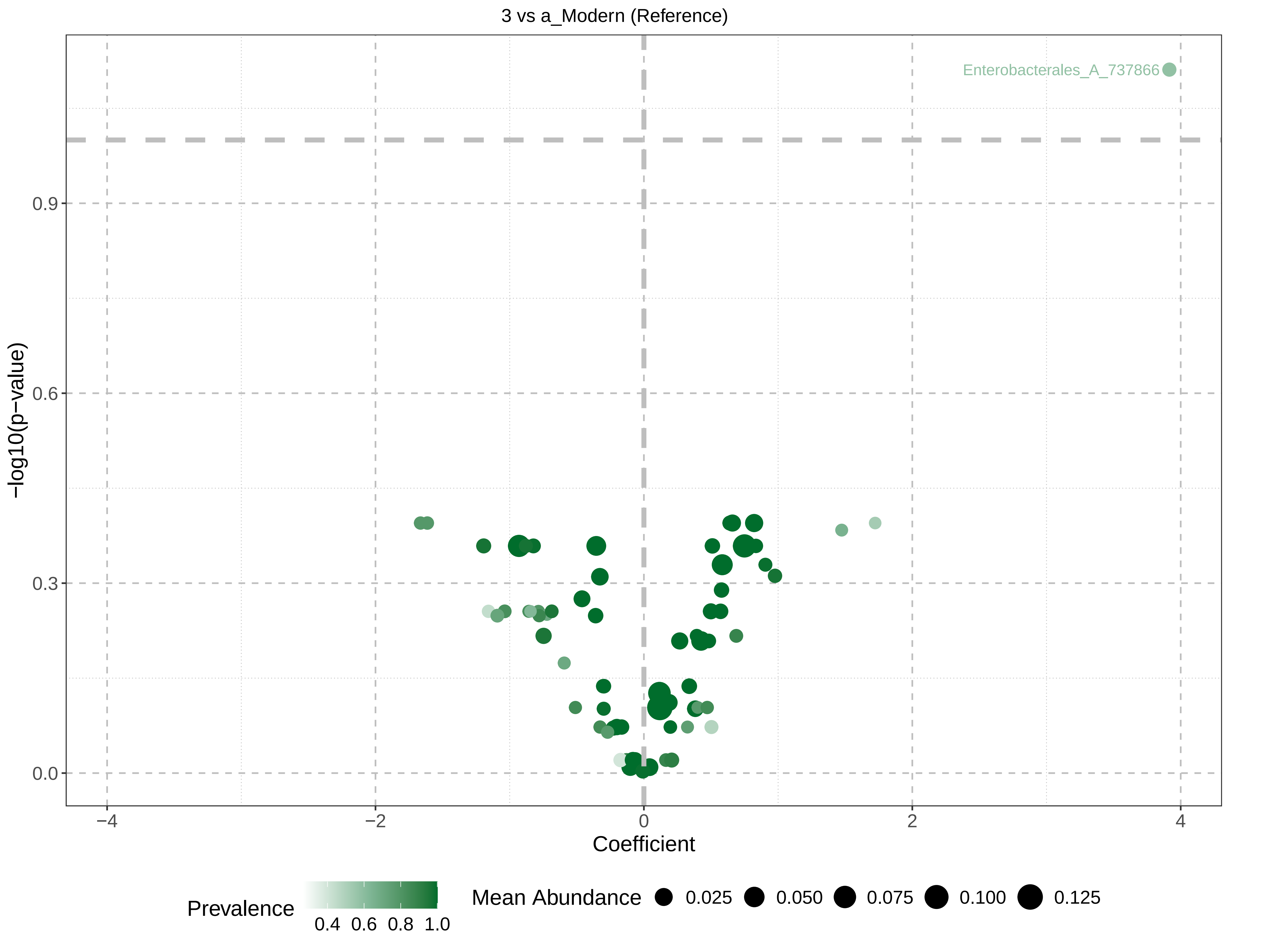

3 vs a_Modern (Reference)
Enterobacterales_A_737866
−log10(p−value)
0.9
0.6
0.3
0.0
−4
−2
0
2
4
Coefficient
Prevalence
0.4
0.6
0.8
1.0
Mean Abundance
0.025
0.050
0.075
0.100
0.125

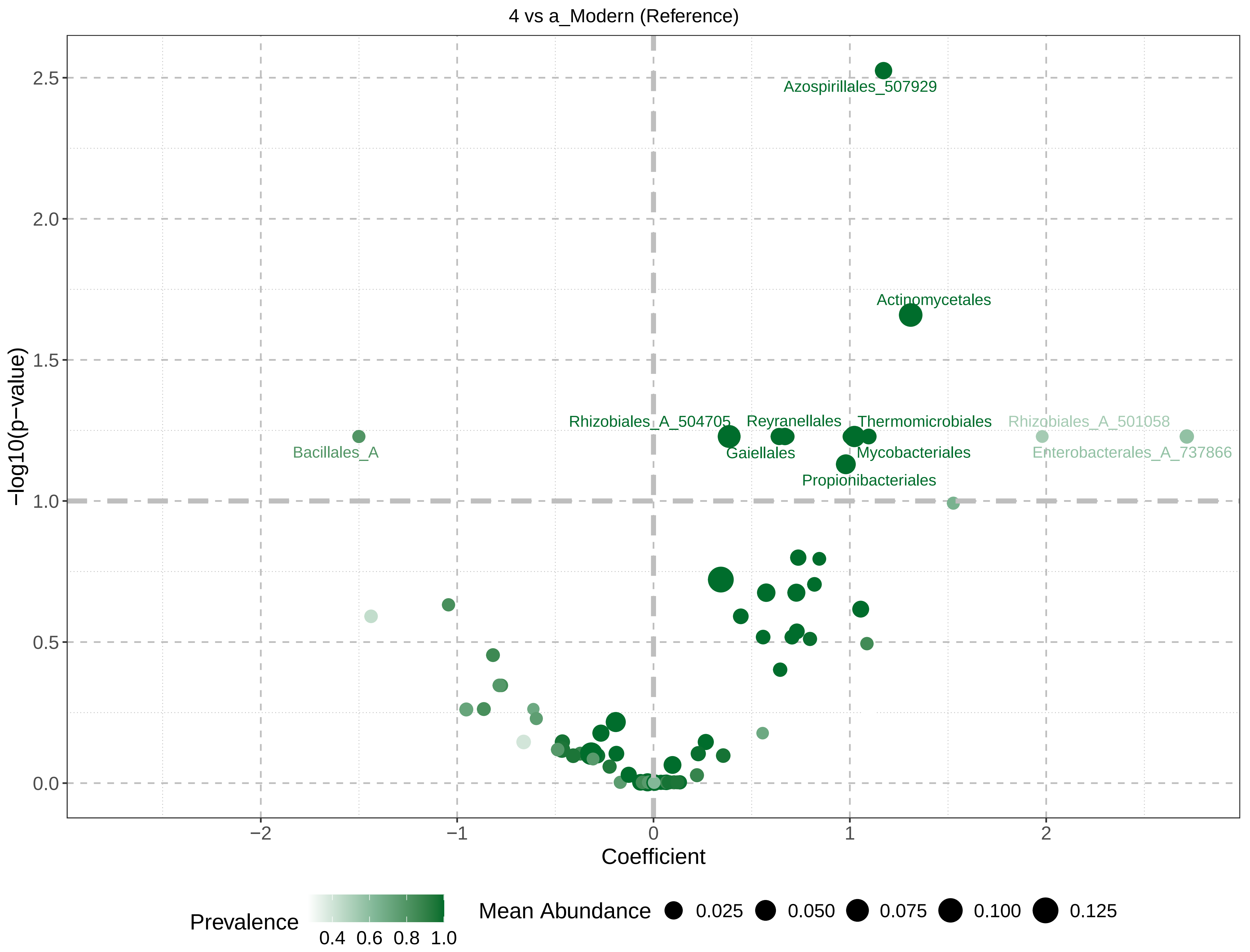
4 vs a_Modern (Reference)
Azospirillales_507929
Actinomycetales
Rhizobiales_A_504705
Reyranellales
Thermomicrobiales
Rhizobiales_A_501058
Bacillales_A
Gaiellales
Mycobacteriales
Enterobacterales_A_737866
Propionibacteriales
−log10(p−value)
2.5
2.0
1.5
1.0
0.5
0.0
−2
−1
0
1
2
Coefficient
Prevalence
0.4 0.6 0.8 1.0
Mean Abundance
0.025
0.050
0.075
0.100
0.125

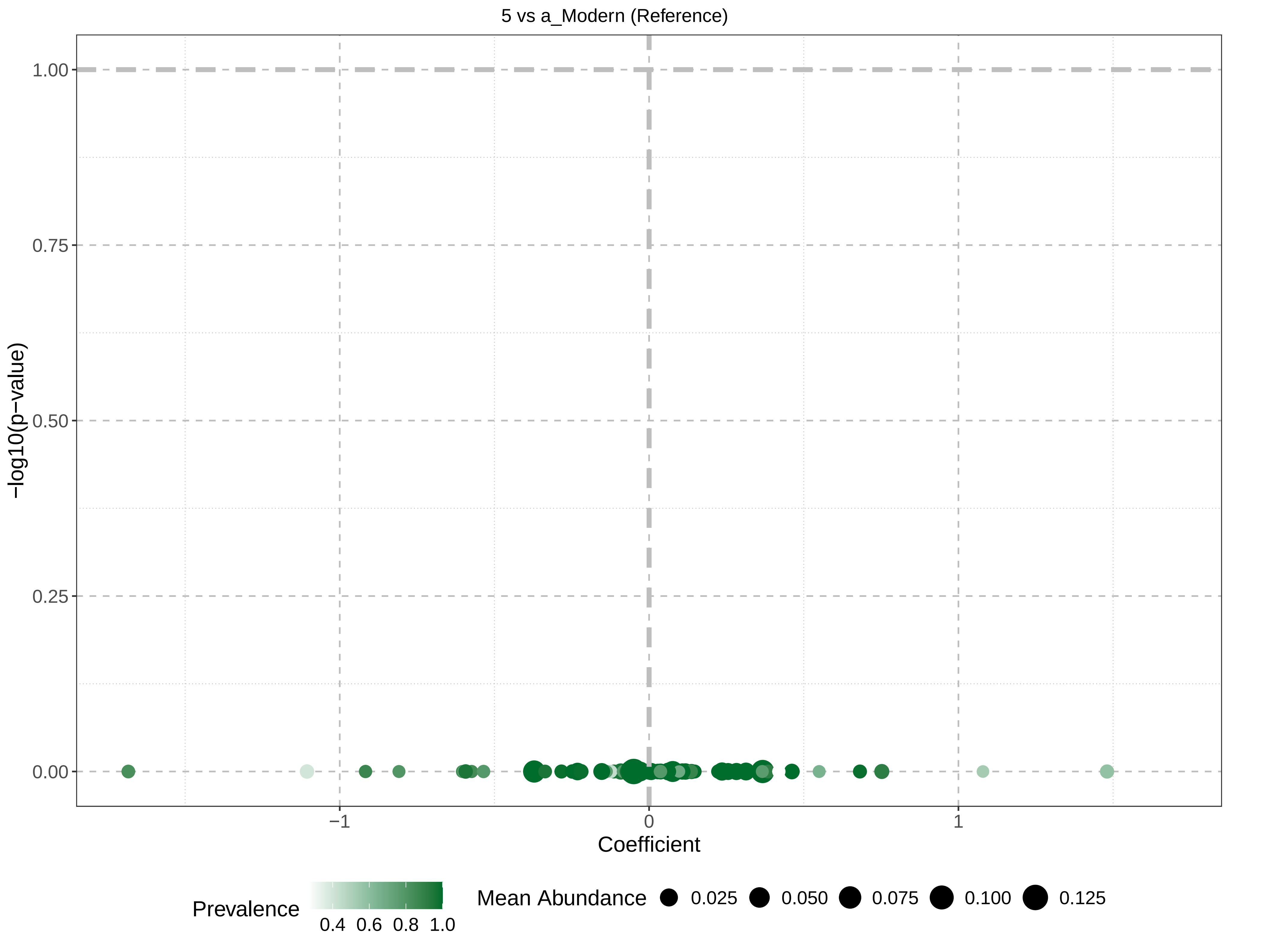
5 vs a_Modern (Reference)
−log10(p−value)
1.00
0.75
0.50
0.25
0.00
−1
0
1
Coefficient
Prevalence
0.4 0.6 0.8 1.0
Mean Abundance
0.025
0.050
0.075
0.100
0.125

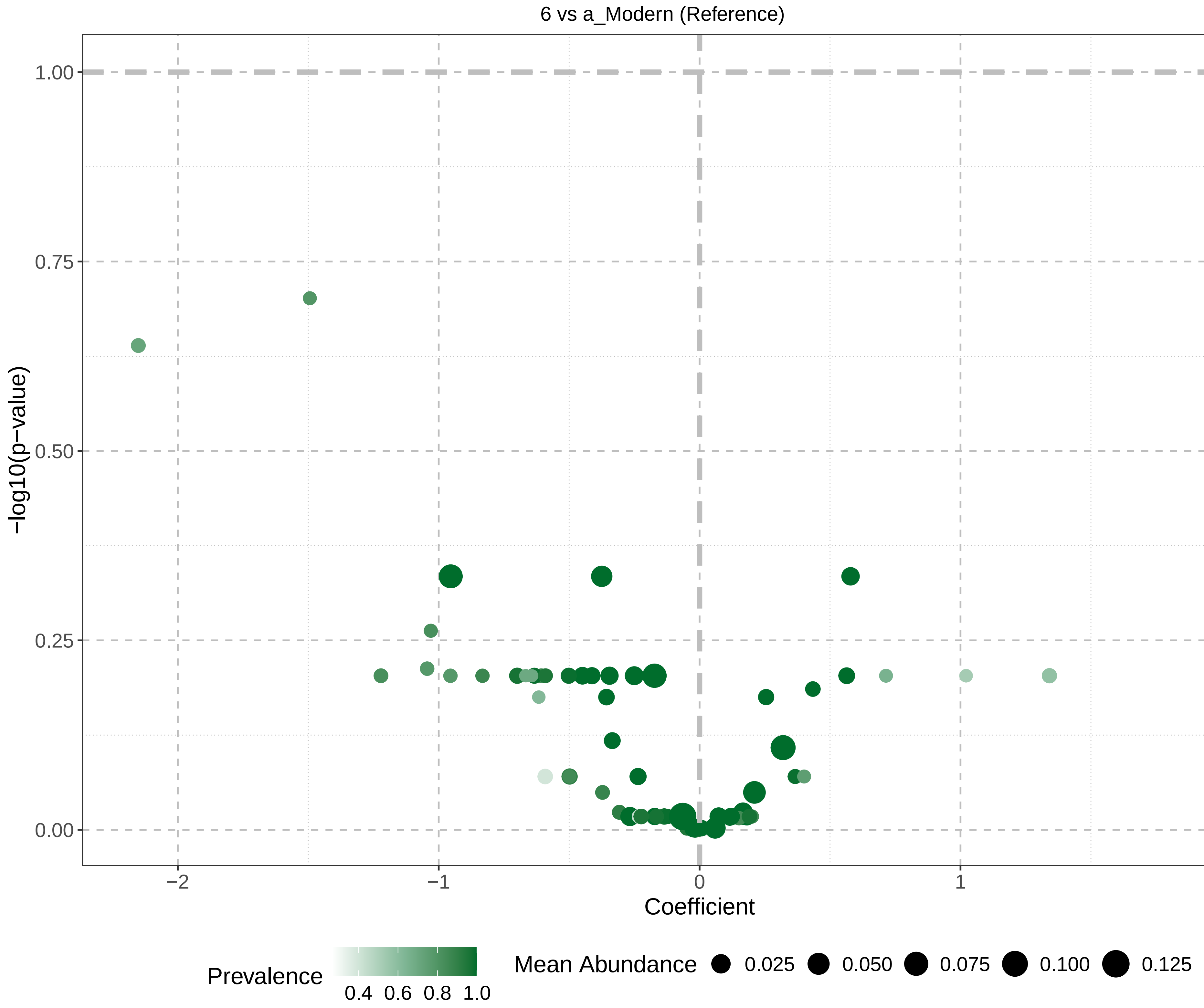

6 vs a_Modern (Reference)
1.00
0.75
0.50
0.25
0.00
−log10(p−value)
−2
−1
0
1
2
Coefficient
Prevalence
0.4 0.6 0.8 1.0
Mean Abundance
0.025
0.050
0.075
0.100
0.125

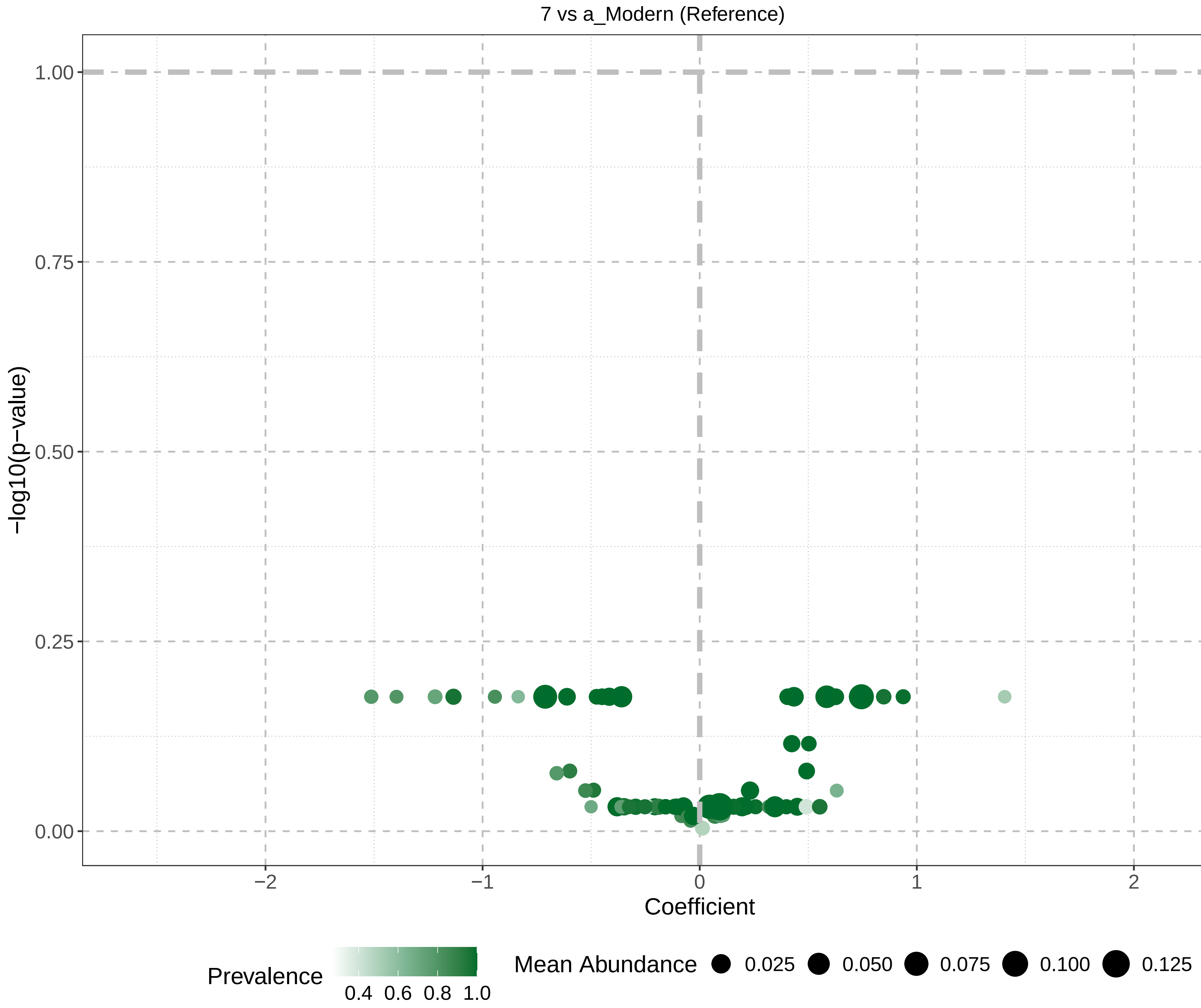
7 vs a_Modern (Reference)
−log10(p−value)
1.00
0.75
0.50
0.25
0.00
−2
−1
0
1
2
Coefficient
Prevalence
0.4 0.6 0.8 1.0
Mean Abundance
0.025
0.050
0.075
0.100
0.125

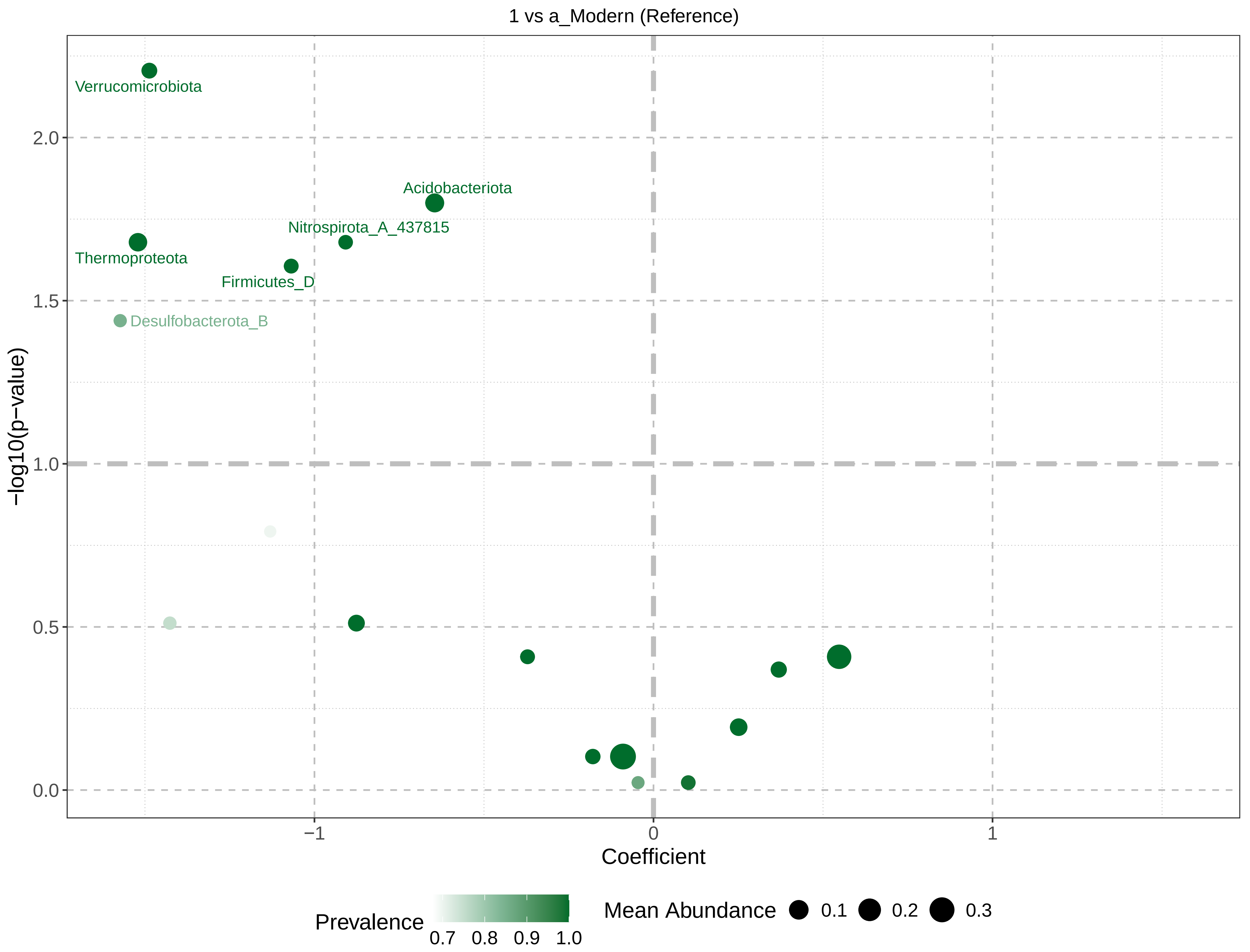
1 vs a_Modern (Reference)
Verrucomicrobiota
Acidobacteriota
Nitrospirota_A_437815
Thermoproteota
Firmicutes_D
Desulfobacterota_B
2.0
1.5
1.0
0.5
0.0
−log10(p−value)
−1
0
1
Coefficient
Prevalence
0.7
0.8
0.9
1.0
Mean Abundance
0.1
0.2
0.3

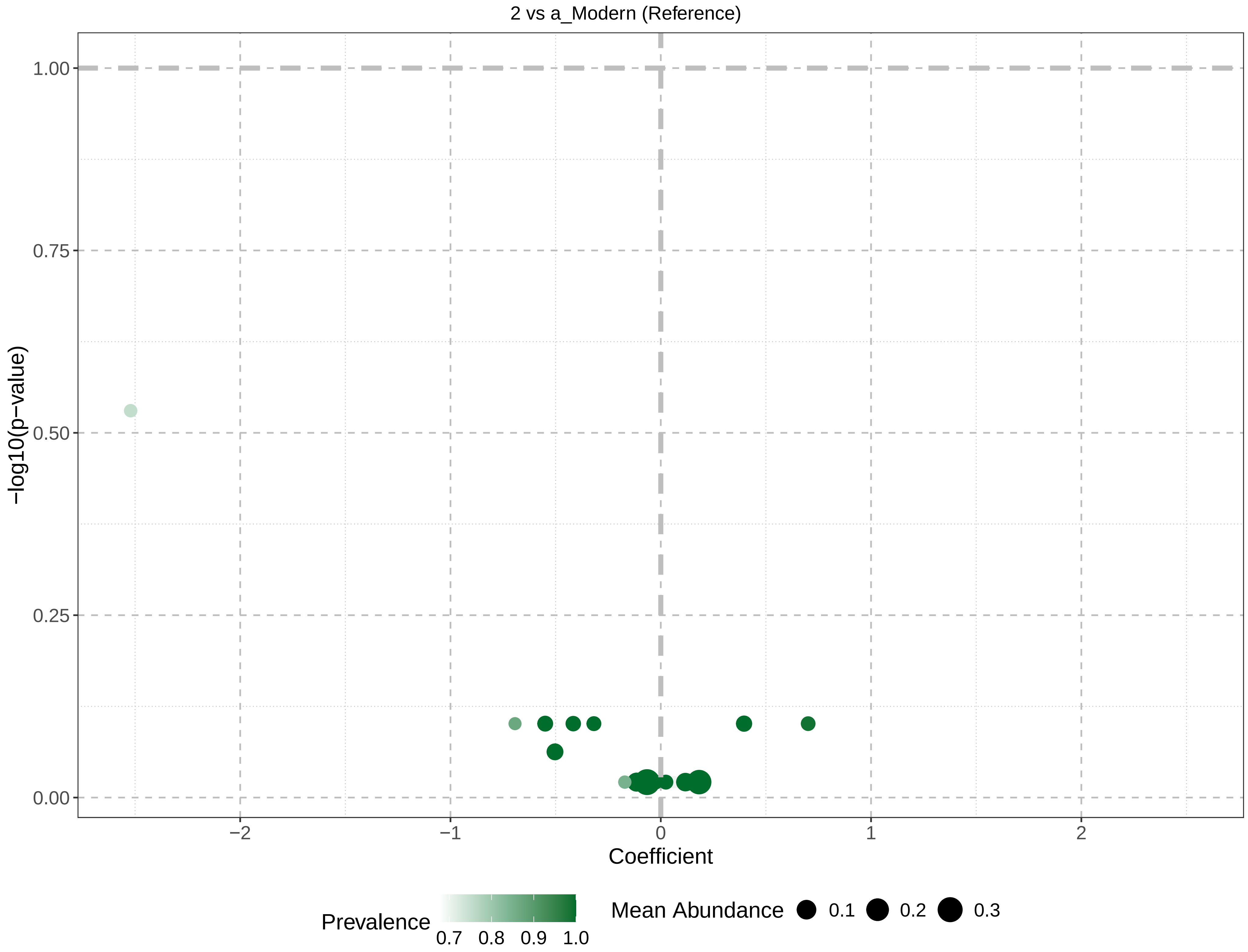
2 vs a_Modern (Reference)
1.00
0.75
0.50
0.25
0.00
−log10(p−value)
−2
−1
0
1
2
Coefficient
Prevalence
0.7
0.8
0.9
1.0
Mean Abundance
0.1
0.2
0.3

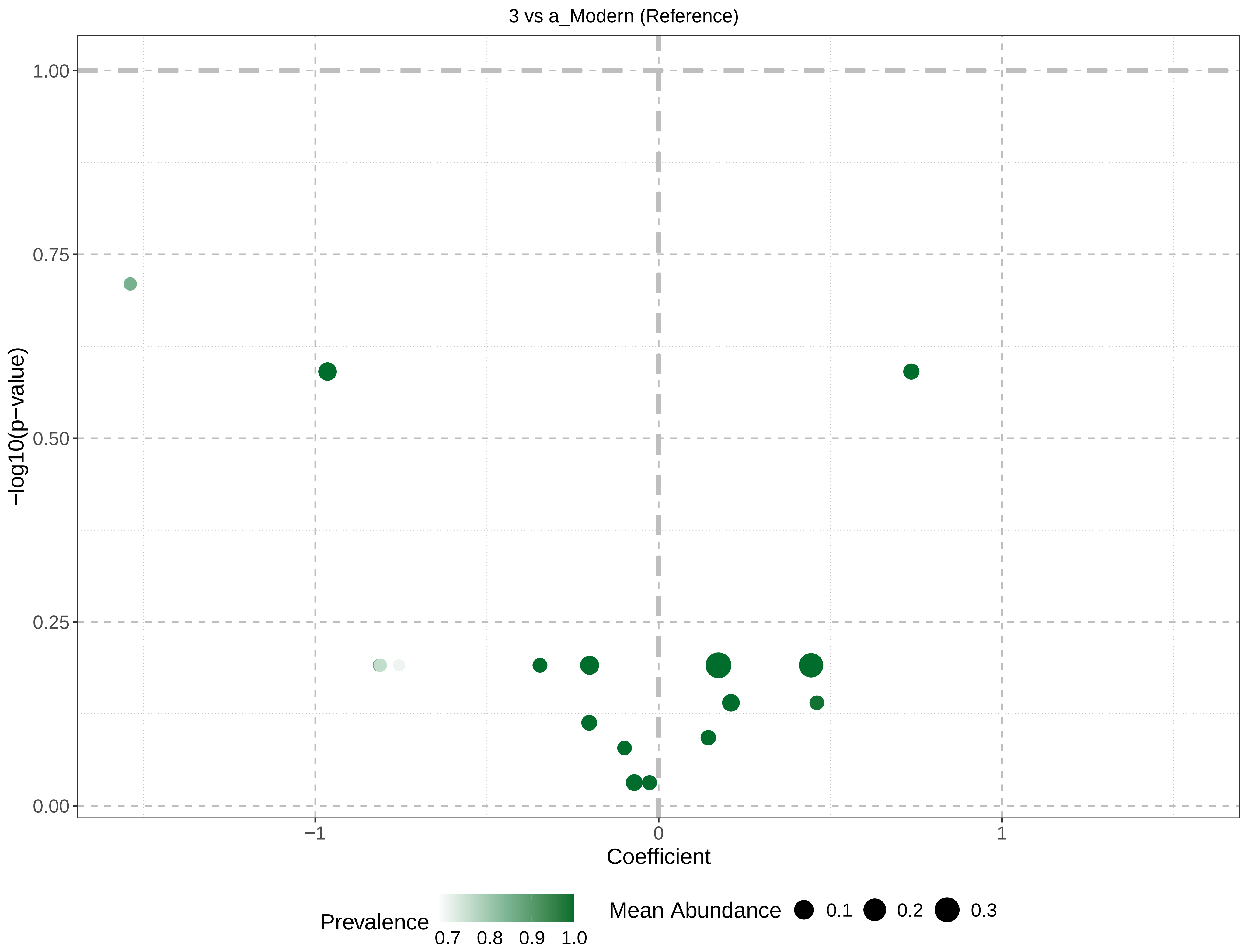

3 vs a_Modern (Reference)
1.00
0.75
0.50
0.25
0.00
−log10(p−value)
−1
0
1
Coefficient
Prevalence
0.7
0.8
0.9
1.0
Mean Abundance
0.1
0.2
0.3

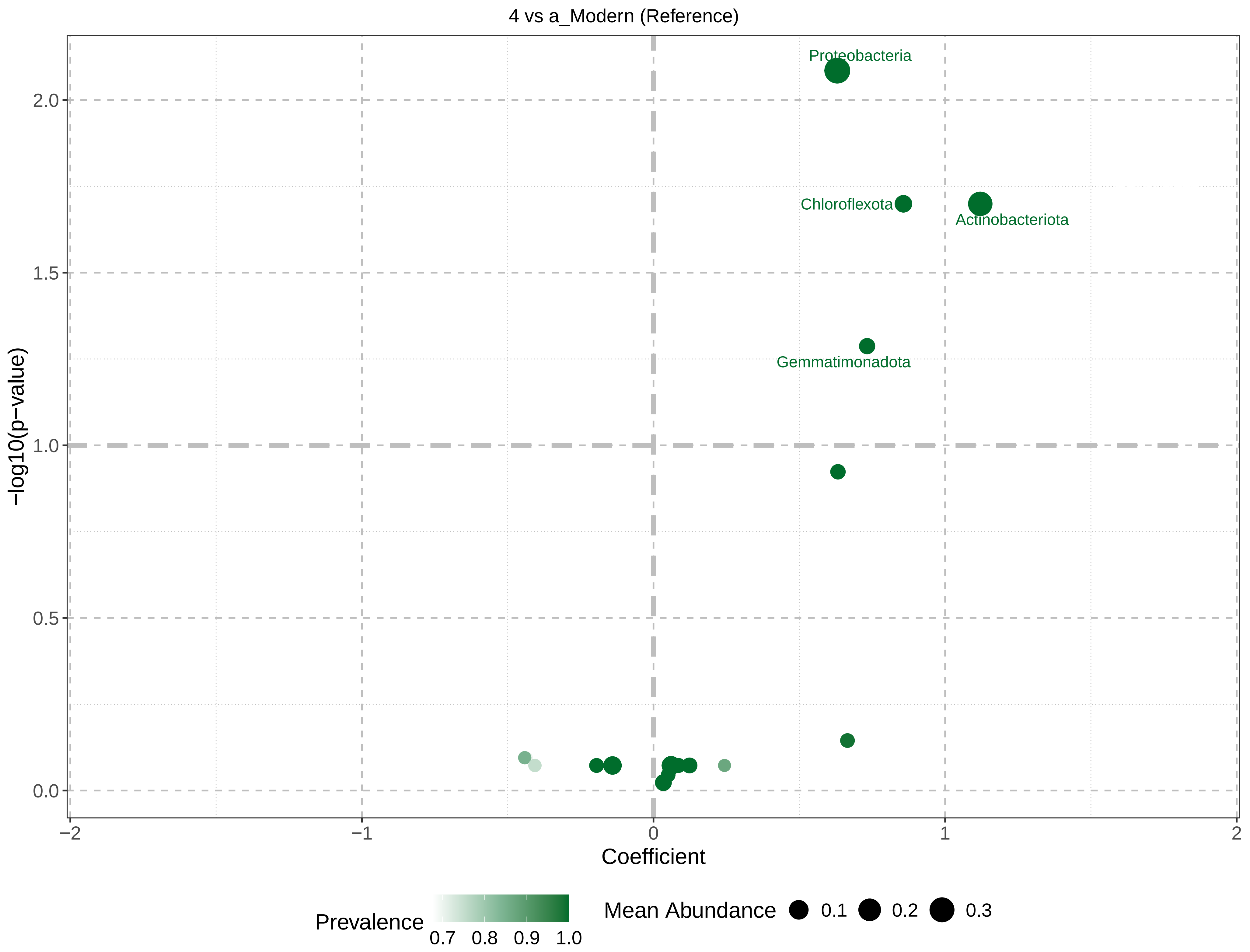
4 vs a_Modern (Reference)
Proteobacteria
Chloroflexota
Actinobacteriota
Gemmatimonadota
-log10(p-value)
2.0
1.5
1.0
0.5
0.0
-2
-1
0
1
2
Coefficient
Prevalence
0.7
0.8
0.9
1.0
Mean Abundance
0.1
0.2
0.3

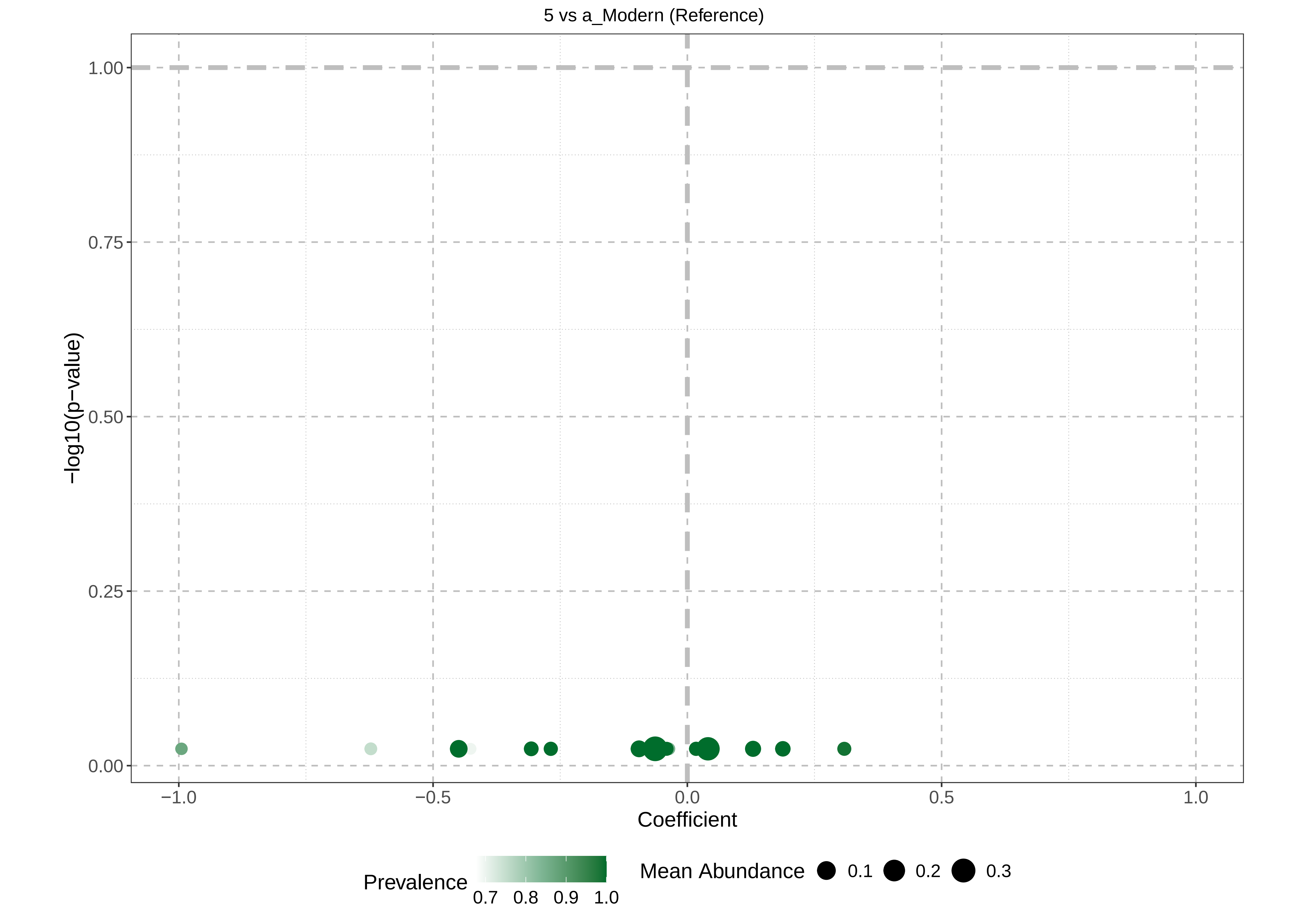

5 vs a_Modern (Reference)
-log10(p-value)
1.00
0.75
0.50
0.25
0.00
−1.0
−0.5
0.0
0.5
1.0
Coefficient
Prevalence
0.7
0.8
0.9
1.0
Mean Abundance
0.1
0.2
0.3

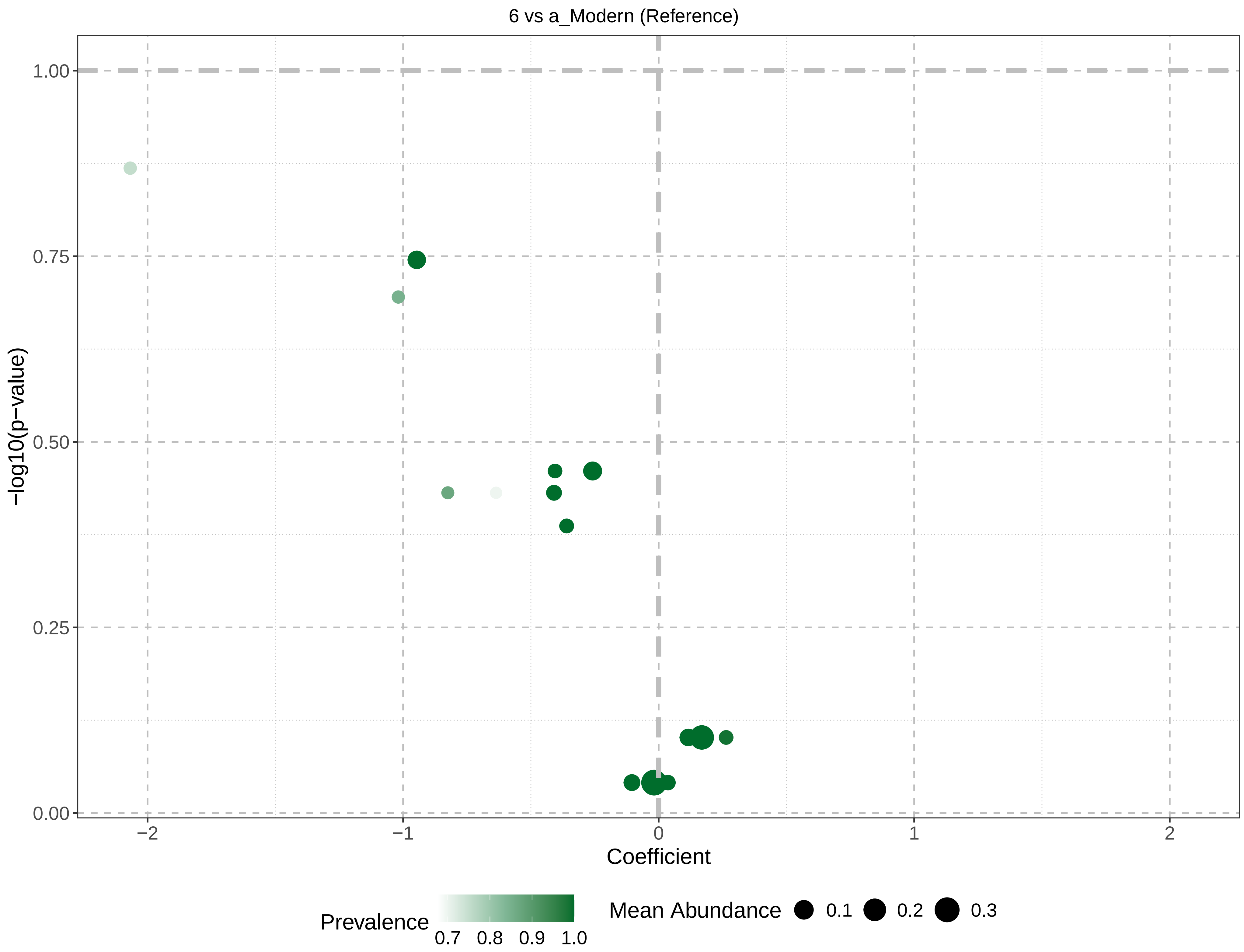
6 vs a_Modern (Reference)
1.00
0.75
0.50
0.25
0.00
−log10(p−value)
−2
−1
0
1
2
Coefficient
Prevalence
0.7
0.8
0.9
1.0
Mean Abundance
0.1
0.2
0.3

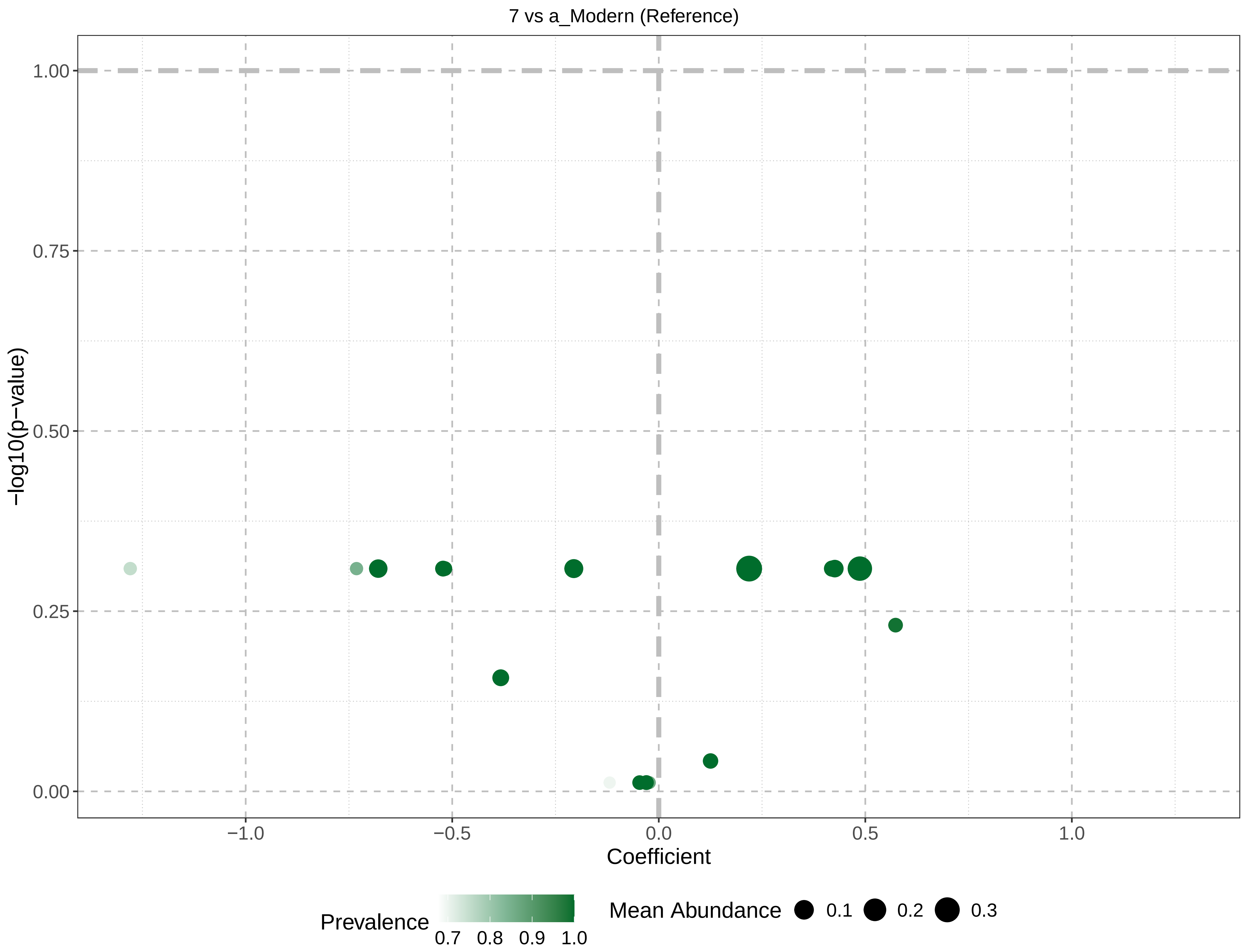
7 vs a_Modern (Reference)
1.00
0.75
0.50
0.25
0.00
−log10(p−value)
−1.0
−0.5
0.0
0.5
1.0
Coefficient
Prevalence
0.7
0.8
0.9
1.0
Mean Abundance
0.1
0.2
0.3

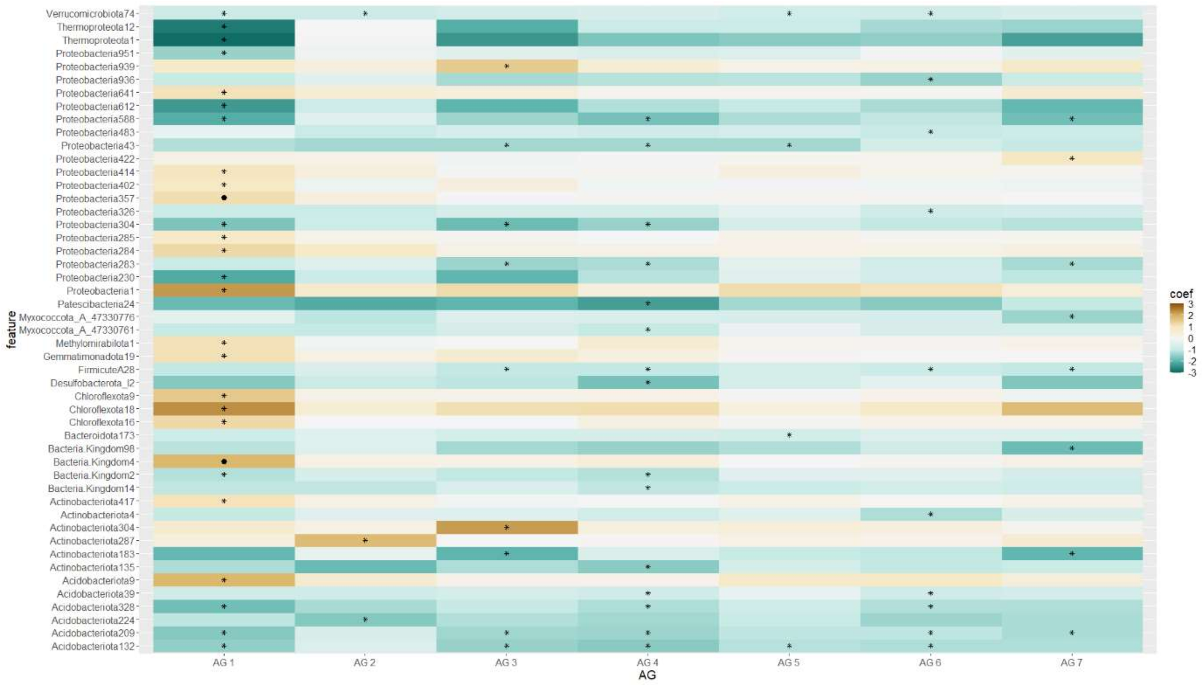


**Figure S8**. Heatmap indicating significant associations (asterisks indicate p-value <0.05, dots indicate q-value < 0.05) between the rhizosphere microbiome taxa abundance (phylum level) and the 81 wheat genotypes grouped into ancestral groups (AG) [2], closely related genetically and two elite (modern) cultivars. Associations were obtained using Multivariate Association with Linear Models (MaAsLin2 [11]). The list of taxa, coefficients, standard error, p-values, and q-values/FDR are provided in Supplementary Table 9.

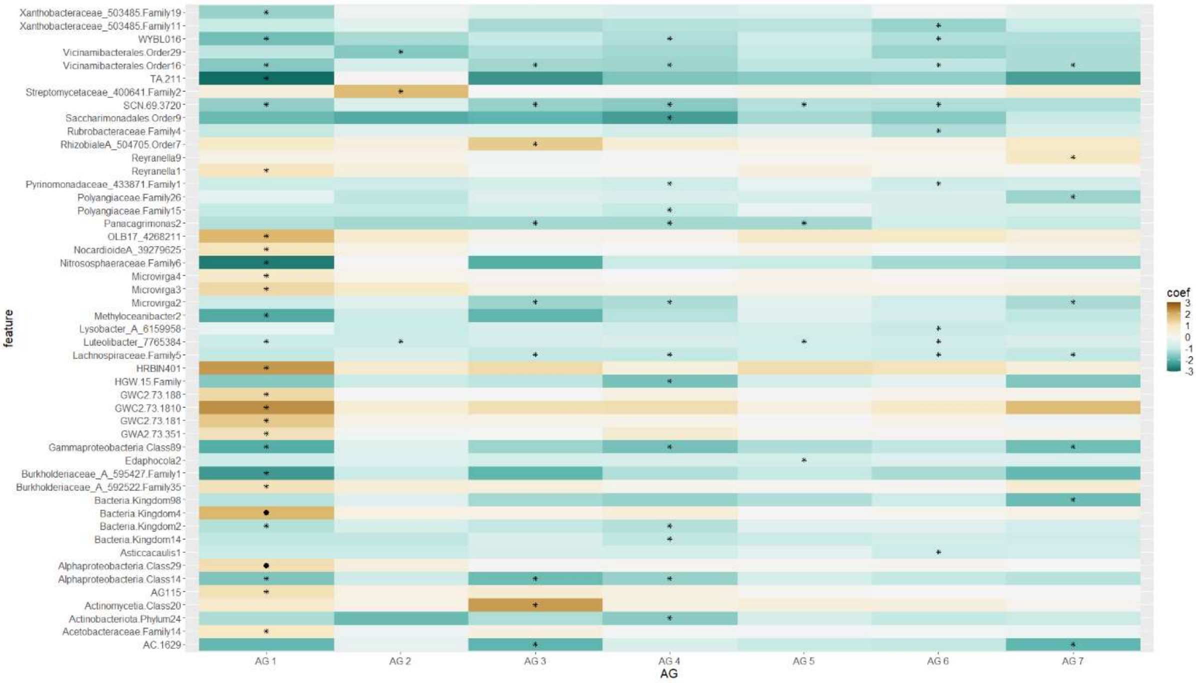


**Figure S9**. Heatmap indicating significant associations (asterisks indicate p-value <0.05, dots indicate q-value < 0.05) between the rhizosphere microbiome taxa abundance (genus level) and the 81 wheat genotypes grouped into ancestral groups (AG) [2], closely related genetically and two elite (modern) cultivars. Associations were obtained using Multivariate Association with Linear Models (R package MaAsLin2 [11]). The list of taxa, coefficients, standard error, p-values, and q-values/FDR are provided in Supplementary Table 10.

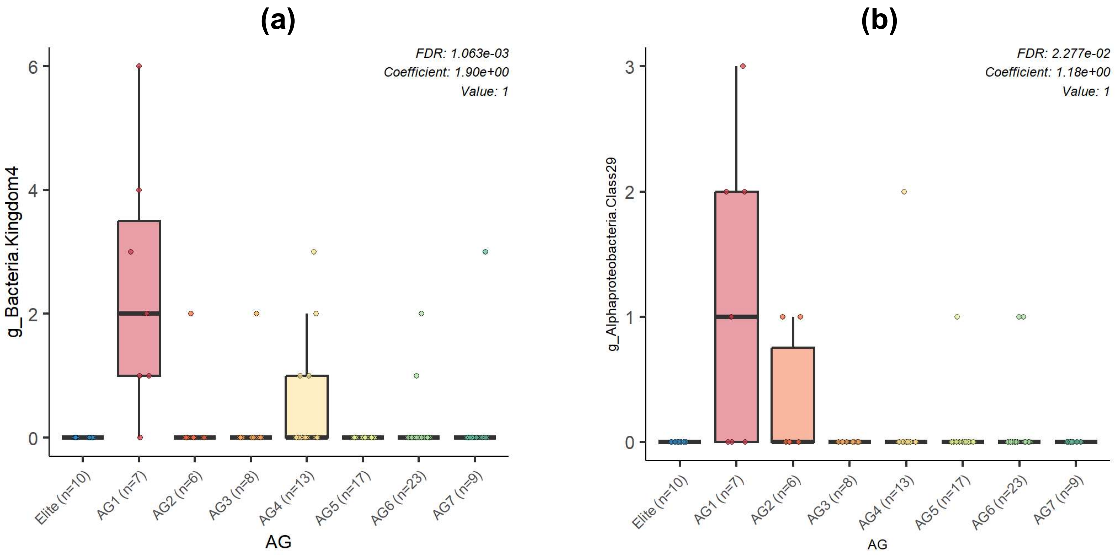


**Figure S10**. Box plots generated by Multivariate Association with Linear Models (R package MaAsLin2 [11]) show the differential abundance of microbial taxa (genus level) between ancestral groups (AG) [2], closely related genetically and elite (modern) cultivars. FDR p-values and regression coefficients are shown in the upper right corner of each plot. The analysis was performed with the following parameters: fixed_effects = c("AG"), reference = c(c("AG, Elite")), random_effects = c("Accessions"), max_significance = 0.05. The x-axis indicates the wheat genotype group (AG), while the y-axis indicates relative abundance. Two unresolved genera were identified as significantly different (q-value < 0.05) between **(a)** elite cultivars and AG1 and AG4, and **(b)** between elite cultivars and AG1 and AG2 at the genus level. The list of taxa, coefficients, standard error, p-values, and q-values/FDR are provided in Supplementary Table 10.

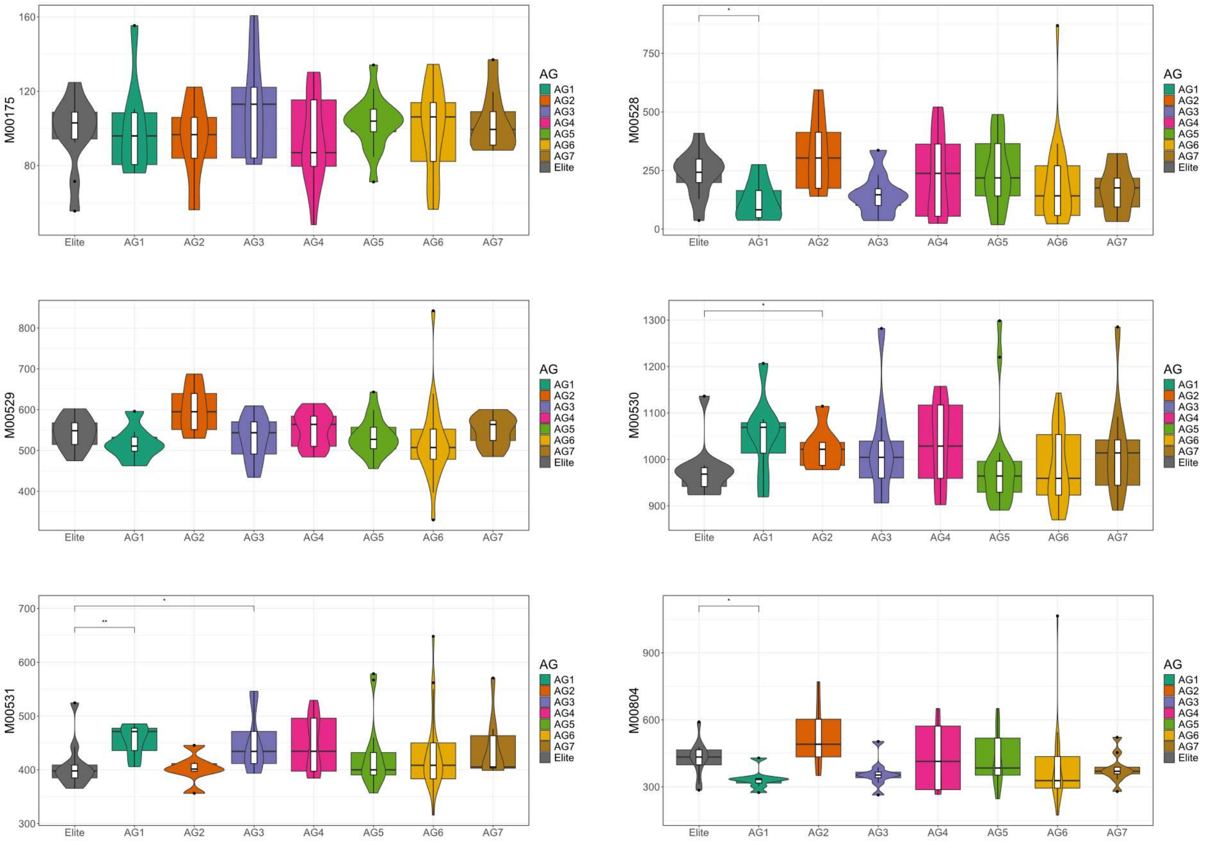


**Figure S11**. Functional analysis of microbial communities in the rhizosphere soil collected from 81 genotypes of hexaploid wheat grouped into ancestral groups (AG) [2], closely related genetically and two elite (modern) cultivars. KEGG [12,13] modules relative abundances for nitrogen metabolism pathways (Supplementary Table 12). **M00175** Nitrogen fixation, nitrogen => ammonia (4) (complete 1/1); **M00531** Assimilatory nitrate reduction, nitrate => ammonia (4) (complete 2/2); **M00530** Dissimilatory nitrate reduction, nitrate => ammonia (9) (complete 2/2); **M00529** Denitrification, nitrate => nitrogen (10) (complete 4/4); **M00528** Nitrification, ammonia => nitrite (4) (complete 2/2); **M00804** Complete nitrification, comammox, ammonia => nitrite => nitrate (6) (complete 3/3). Significant differences were calculated by pairwise comparison using the Wilcoxon test and the Benjamini-Hochberg method (R packages ggpubr [8] and ggplot2 [9]) for the adjusted p-value (*$p \le 0.05$, **$p \le 0.01$, ***$p \le 0.001$).

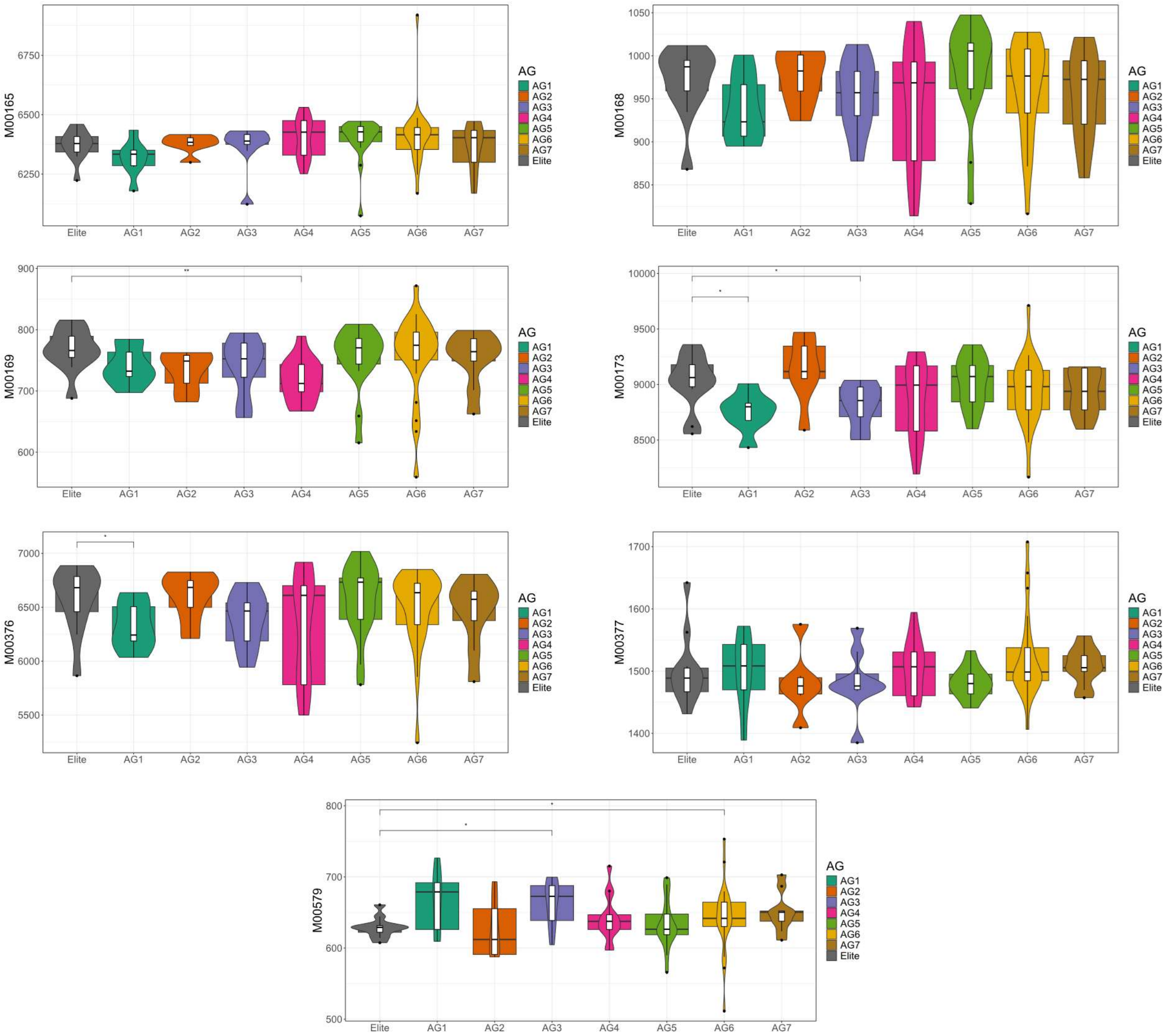


**Figure S12**. Functional analysis of microbial communities in the rhizosphere soil collected from 81 genotypes of hexaploid wheat grouped into ancestral groups (AG) [2], closely related genetically and two elite (modern) cultivars. KEGG [12,13] modules relative abundances for carbon fixation pathways (Supplementary Table 13). **M00165** Reductive pentose phosphate cycle (Calvin cycle) (17) (complete 11/11); **M00168** Carbon fixation by Calvin cycle; **M00169** Carbon fixation by Calvin cycle; **M00173** Reductive citrate cycle (Arnon-Buchanan cycle) (32) (complete 10/10); **M00376** 3-Hydroxypropionate bi-cycle (21) (complete 13/13); **M00377** Reductive acetyl-CoA pathway (Wood-Ljungdahl pathway) (10) (complete 7/7); **M00579** Phosphate acetyltransferase-acetate kinase pathway, acetyl-CoA => acetate (4) (complete 2/2). Significant differences were calculated by pairwise comparison using the Wilcoxon test and the Benjamini-Hochberg method (R packages ggpubr [8] and ggplot2 [9]) for the adjusted p-value (*p ≤ 0.05, **p ≤ 0.01, ***p ≤ 0.001).

**Figure S13.** Differential network analyses were constructed using the R package NetCoMi [5] to compare the network structure and taxa associations between 81 genotypes of hexaploid wheat grouped into ancestral groups (AG) [2], closely related genetically, AG1 **(a)**, AG2 **(b)**, AG3 **(c)**, AG4 **(d)**, AG5 **(e)**, AG6 **(f)** and AG7 **(g),** and two elite cultivars used as reference for modern genotypes. Nodes indicate taxa (genus level and edge colours indicate the direction of change in the strength or type of association between taxa for the groups compared.

**(a)**

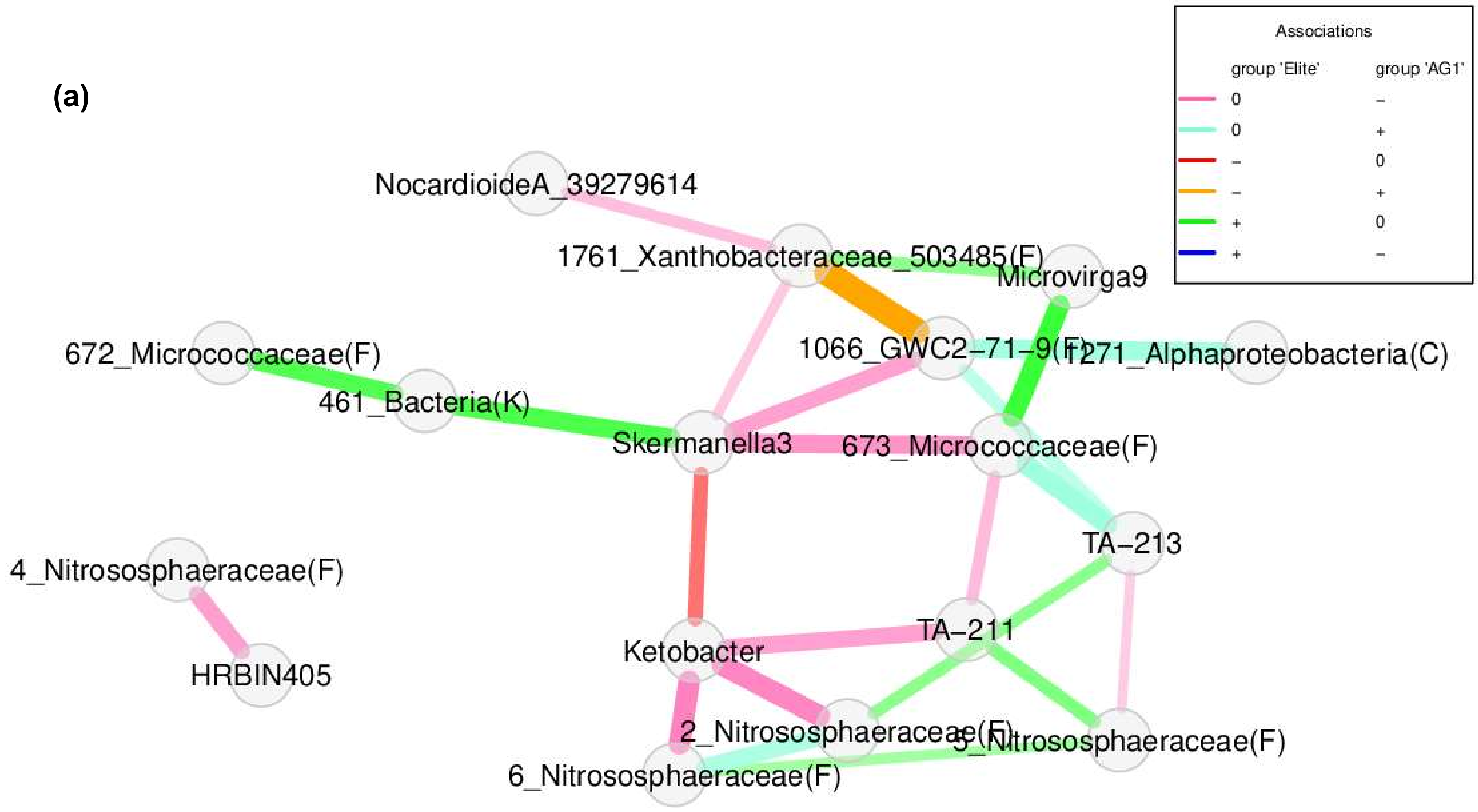

(b)

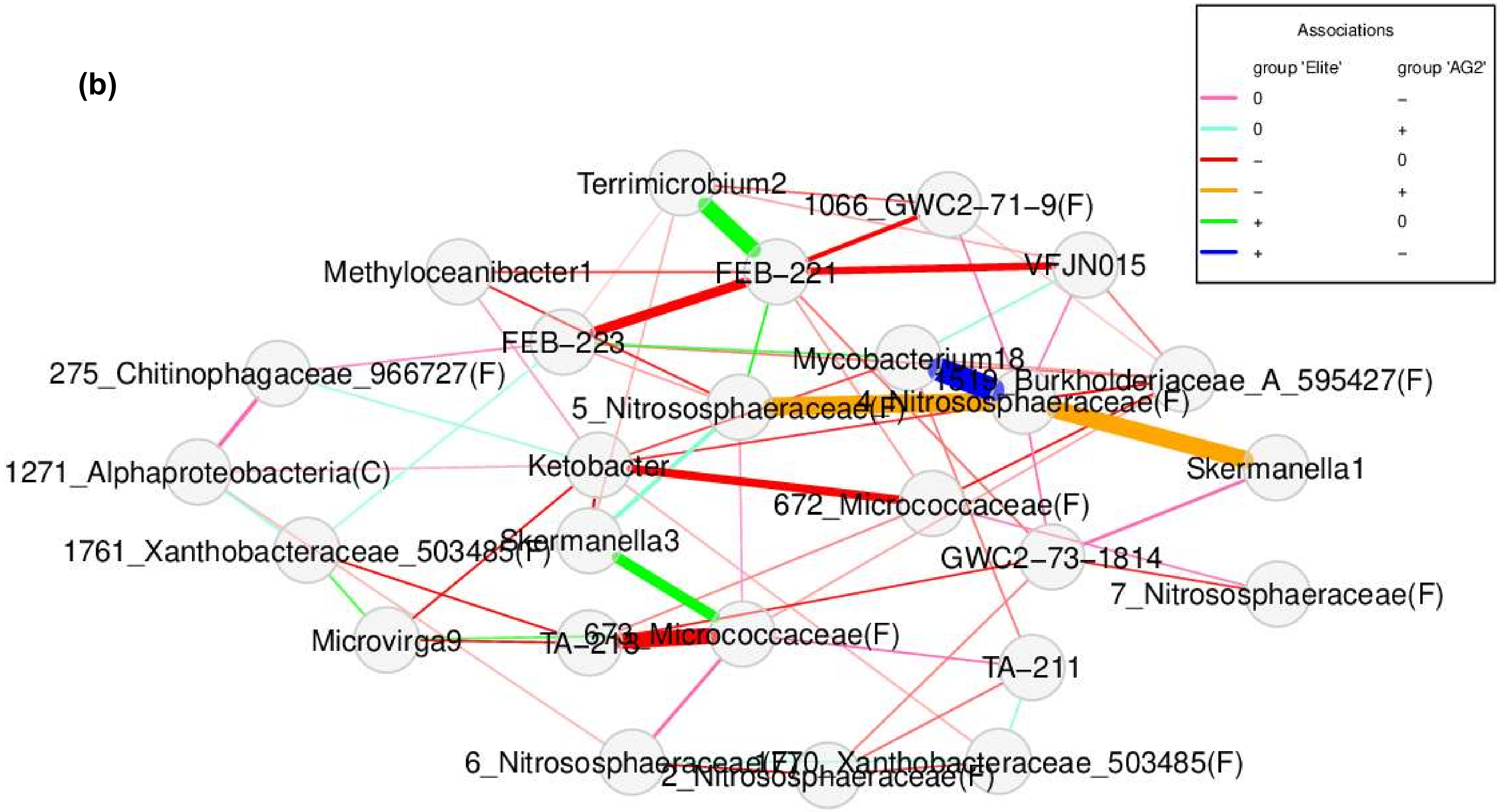
Associations
group 'Elite'
group 'AG2'
0 –
0 +
– 0
– +
+ 0
+ –
Terrimicrobium2
1066_GWC2–71–9(F)
Methyloceanibacter1
FEB–221
VFJN015
FEB–223
275_Chitinophagaceae_966727(F)
Mycobacterium18
Burkholderiaceae_A_595427(F)
5_Nitrososphaeraceae(F)
4_Nitrososphaeraceae(F)
Skermanella1
1271_Alphaproteobacteria(C)
Ketobacter
672_Micrococcaceae(F)
1761_Xanthobacteraceae_503485(F)
Skermanella3
GWC2–73–1814
7_Nitrososphaeraceae(F)
Microvirga9
673_Micrococcaceae(F)
TA–211
6_Nitrososphaeraceae(F)
2_Nitrososphaeraceae(F)
Xanthobacteraceae_503485(F)

(c)

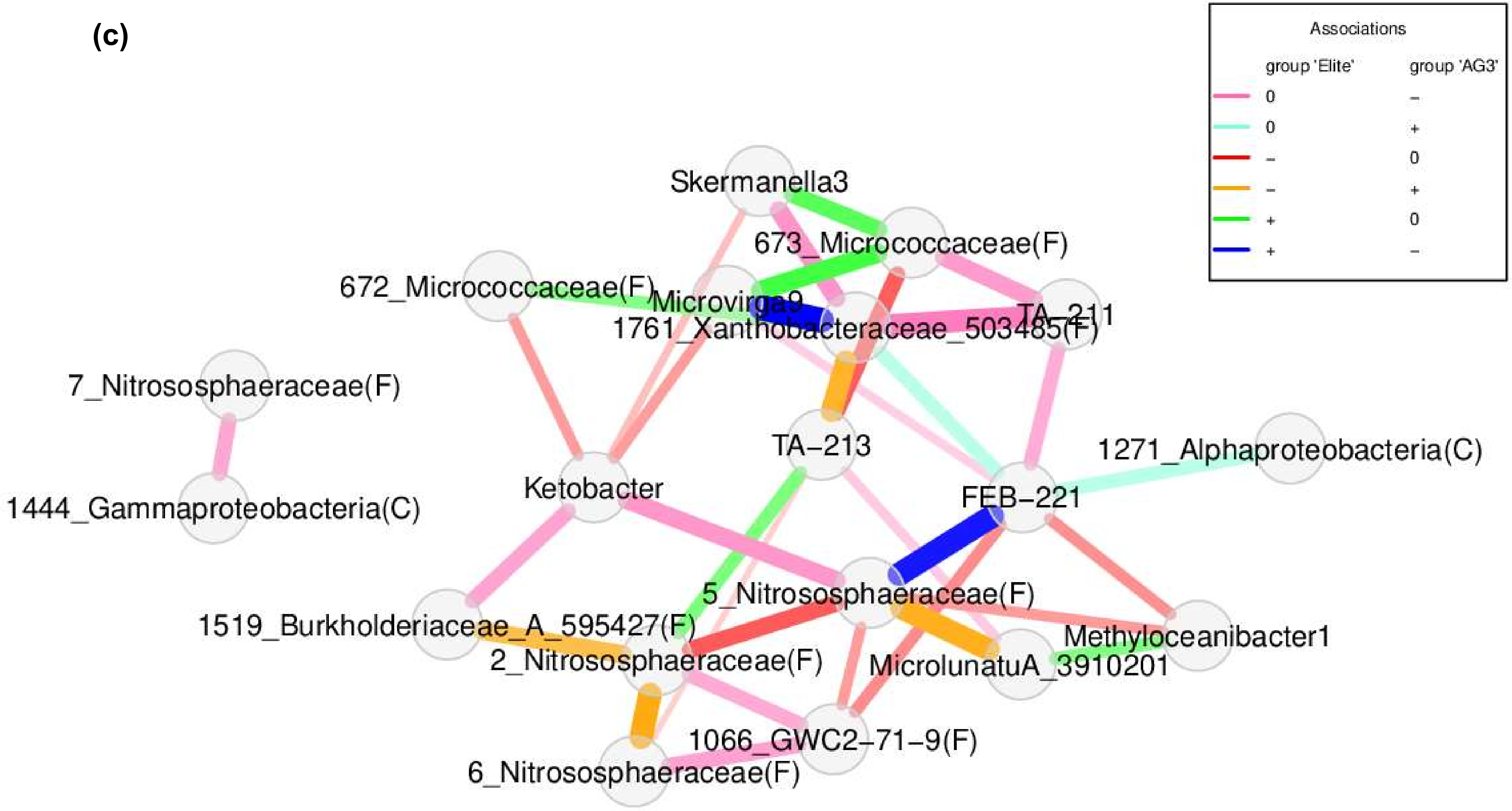
Associations
group 'Elite'
group 'AG3'
0 –
0 +
– 0
– +
+ 0
+ –
Skermanella3
673_Micrococcaceae(F)
672_Micrococcaceae(F)
Microvirga9
1761_Xanthobacteraceae_503485(F)
TA–211
7_Nitrososphaeraceae(F)
TA–213
1271_Alphaproteobacteria(C)
1444_Gammaproteobacteria(C)
Ketobacter
FEB–221
5_Nitrososphaeraceae(F)
1519_Burkholderiaceae_A_595427(F)
Methyloceanibacter1
2_Nitrososphaeraceae(F)
MicrolunatuA_3910201
1066_GWC2–71–9(F)
6_Nitrososphaeraceae(F)

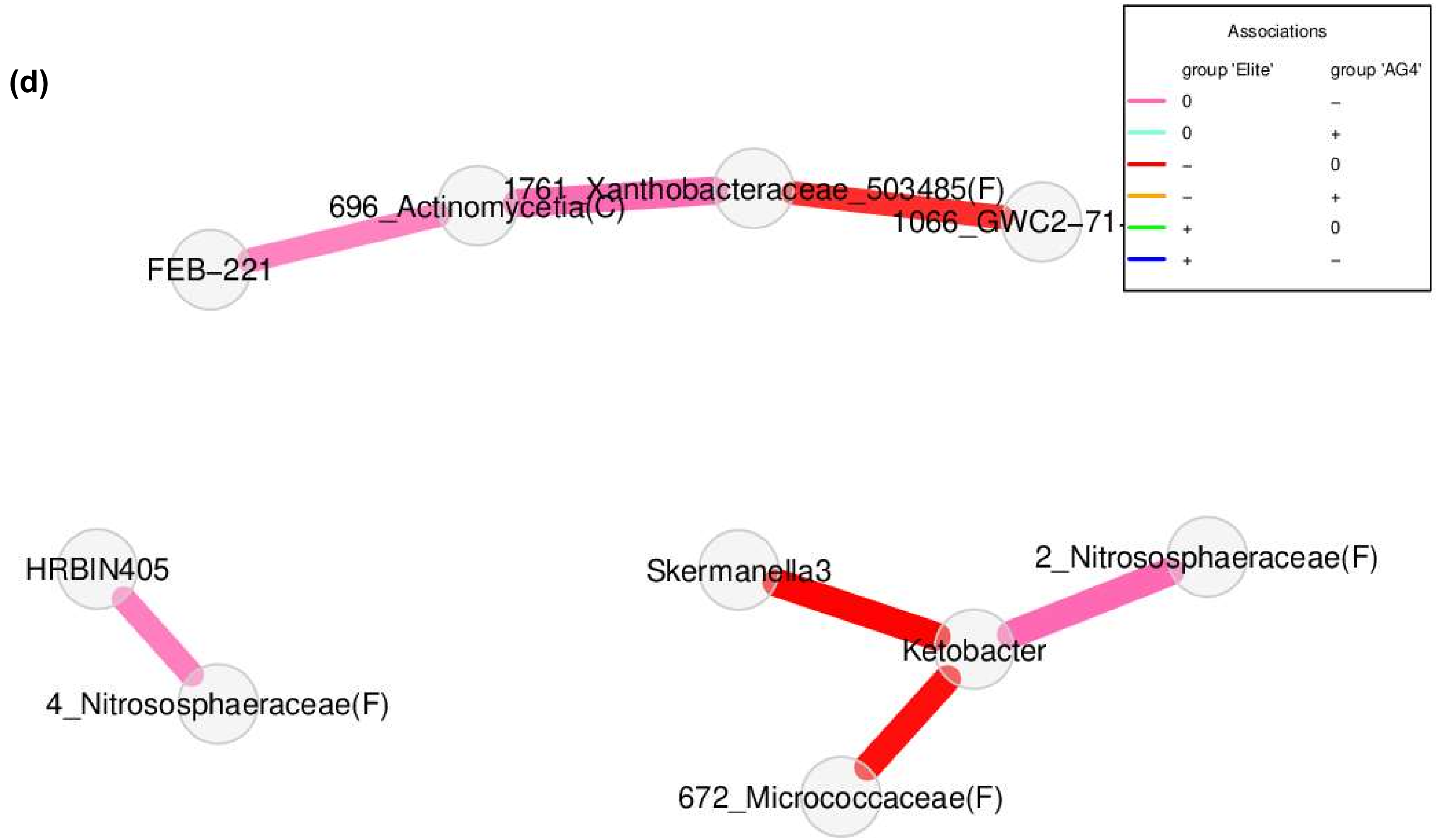
(d)
Associations
group 'Elite'
group 'AG4'
0 –
0 +
– 0
– +
+ 0
+ –
FEB–221
696_Actinomycetia(C)
1761_Xanthobacteraceae_503485(F)
1066_GWC2–71
HRBIN405
4_Nitrososphaeraceae(F)
Skermanella3
2_Nitrososphaeraceae(F)
Ketobacter
672_Micrococcaceae(F)

(e)

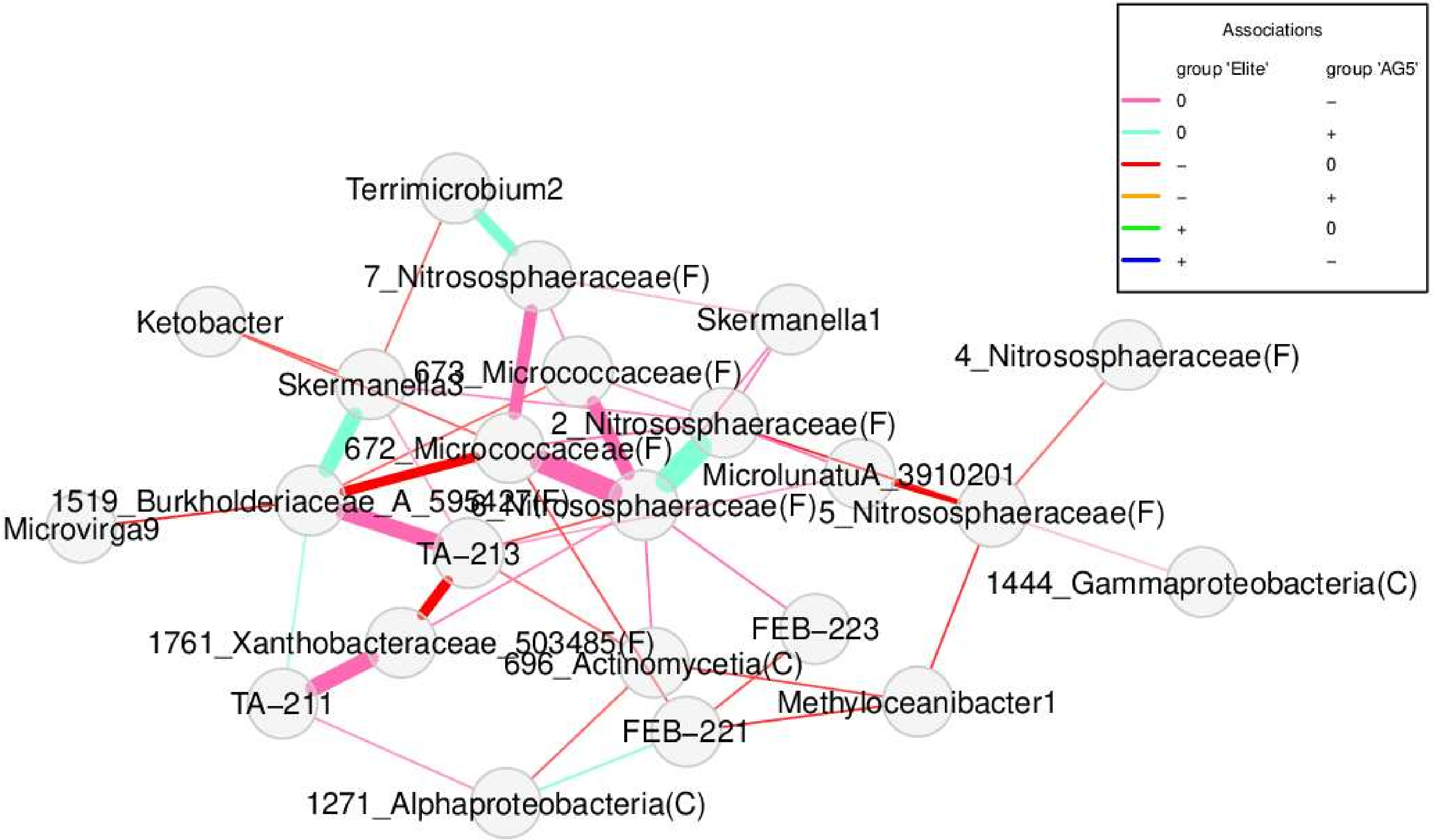
Associations
group 'Elite'
group 'AG5'
0 –
0 +
– 0
– +
+ 0
+ –
Terrimicrobium2
7_Nitrososphaeraceae(F)
Ketobacter
Skermanella1
4_Nitrososphaeraceae(F)
Skermanella3
673_Micrococcaceae(F)
2_Nitrososphaeraceae(F)
672_Micrococcaceae(F)
MicrolunatuA_3910201
1519_Burkholderiaceae_A_595427(F)
6_Nitrososphaeraceae(F)
5_Nitrososphaeraceae(F)
Microvirga9
TA–213
1444_Gammaproteobacteria(C)
1761_Xanthobacteraceae_503485(F)
FEB–223
696_Actinomycetia(C)
TA–211
Methyloceanibacter1
FEB–221
1271_Alphaproteobacteria(C)

(f)

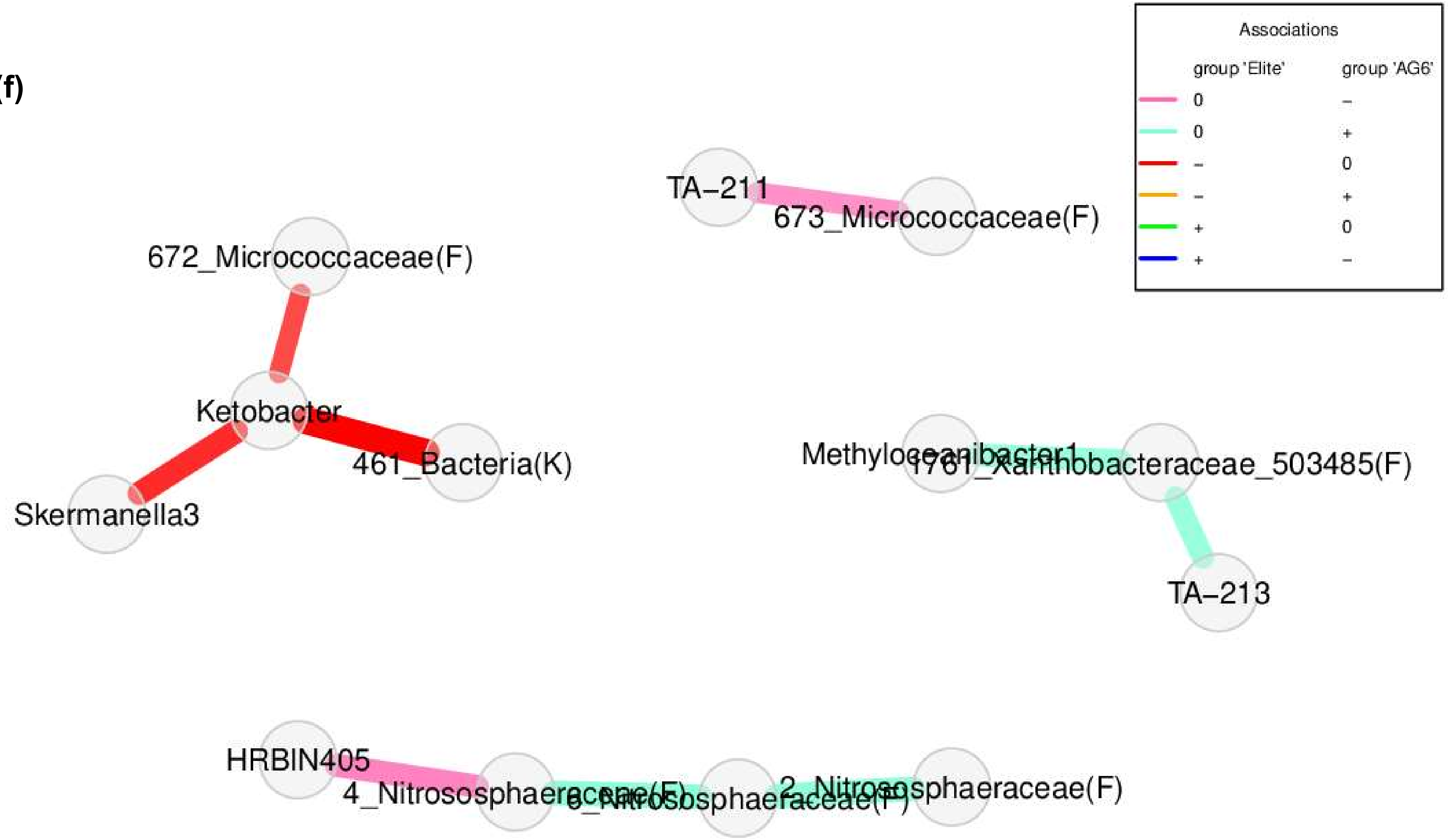
Associations
group 'Elite'
group 'AG6'
0
–
0
+
–
0
–
+
+
0
+
–
TA–211
673_Micrococcaceae(F)
672_Micrococcaceae(F)
Ketobacter
461_Bacteria(K)
Skermanella3
TA–213
HRBIN405
4_Nitrososphaeraceae(F)

(g)

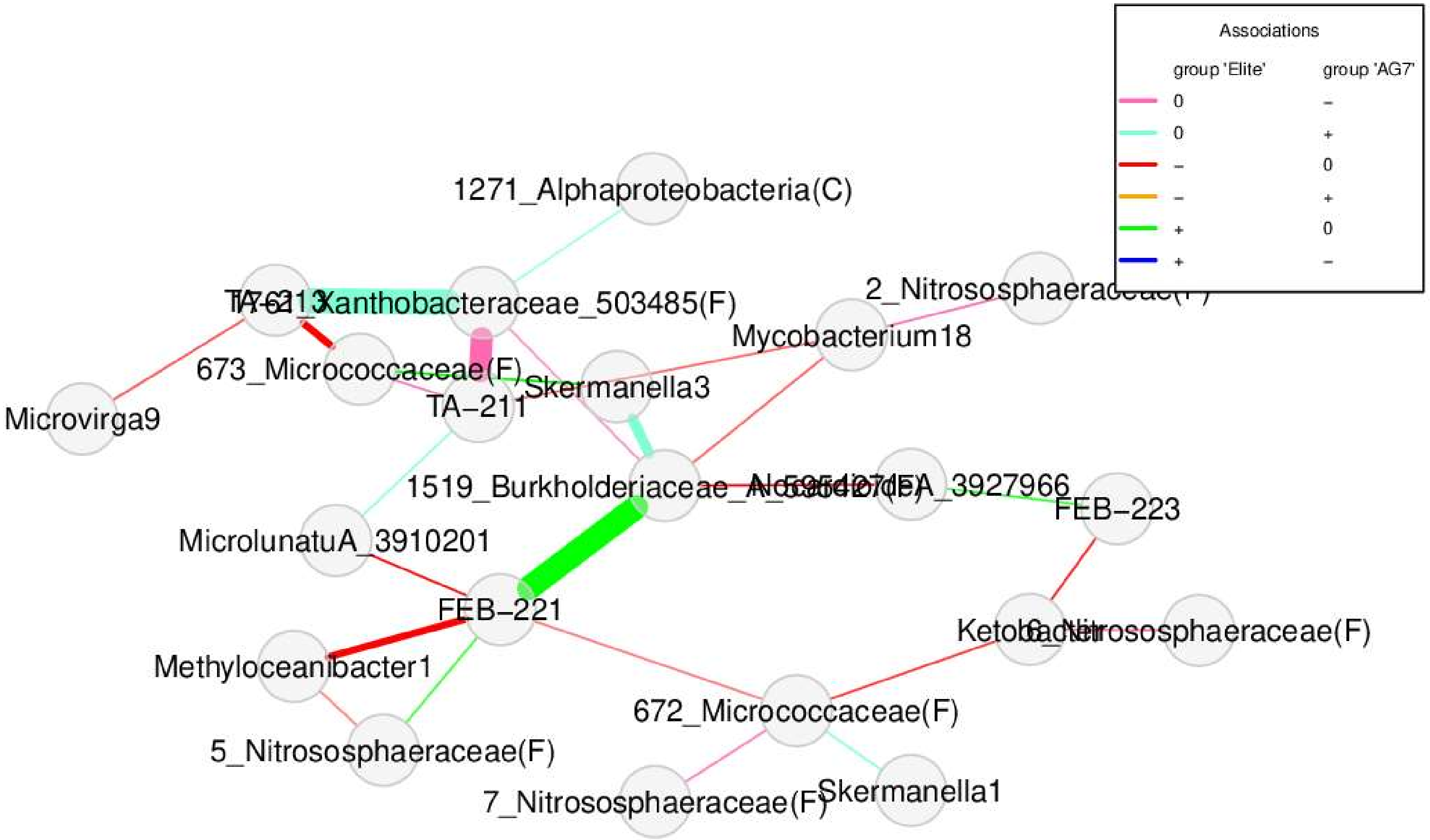

Associations
group 'Elite'
group 'AG7'
0
–
0
+
–
0
–
+
+
0
+
–
1271_Alphaproteobacteria(C)
Xanthobacteraceae_503485(F)
673_Micrococcaceae(F)
Skermanella3
TA–211
Microvirga9
Mycobacterium18
1519_Burkholderiaceae
FEB–223
MicrolunatuA_3910201
FEB–221
Methyloceanibacter1
5_Nitrososphaeraceae(F)
672_Micrococcaceae(F)
7_Nitrososphaeraceae(F)
Skermanella1

**Figure S14.** Association networks were constructed using the R package NetCoMi [5] to assess differences in taxa associations between 81 genotypes of hexaploid wheat grouped into ancestral groups (AG) [2], closely related genetically, AG1 **(a)**, AG2 **(b)**, AG3 **(c)**, AG4 **(d)**, AG5 **(e)**, AG6 **(f)** and AG7 **(g),** and two elite cultivars used as reference for modern genotypes. Networks only include taxa that are differentially associated between the two groups compared. Nodes correspond to taxa (genus level), and edges indicate significant interactions among nodes. The results of the comparison analyses are provided in Supplementary Tables 15 to 21. The Jaccard similarity coefficients obtained for the topological measures of the networks are also summarised in Supplementary Table 22.

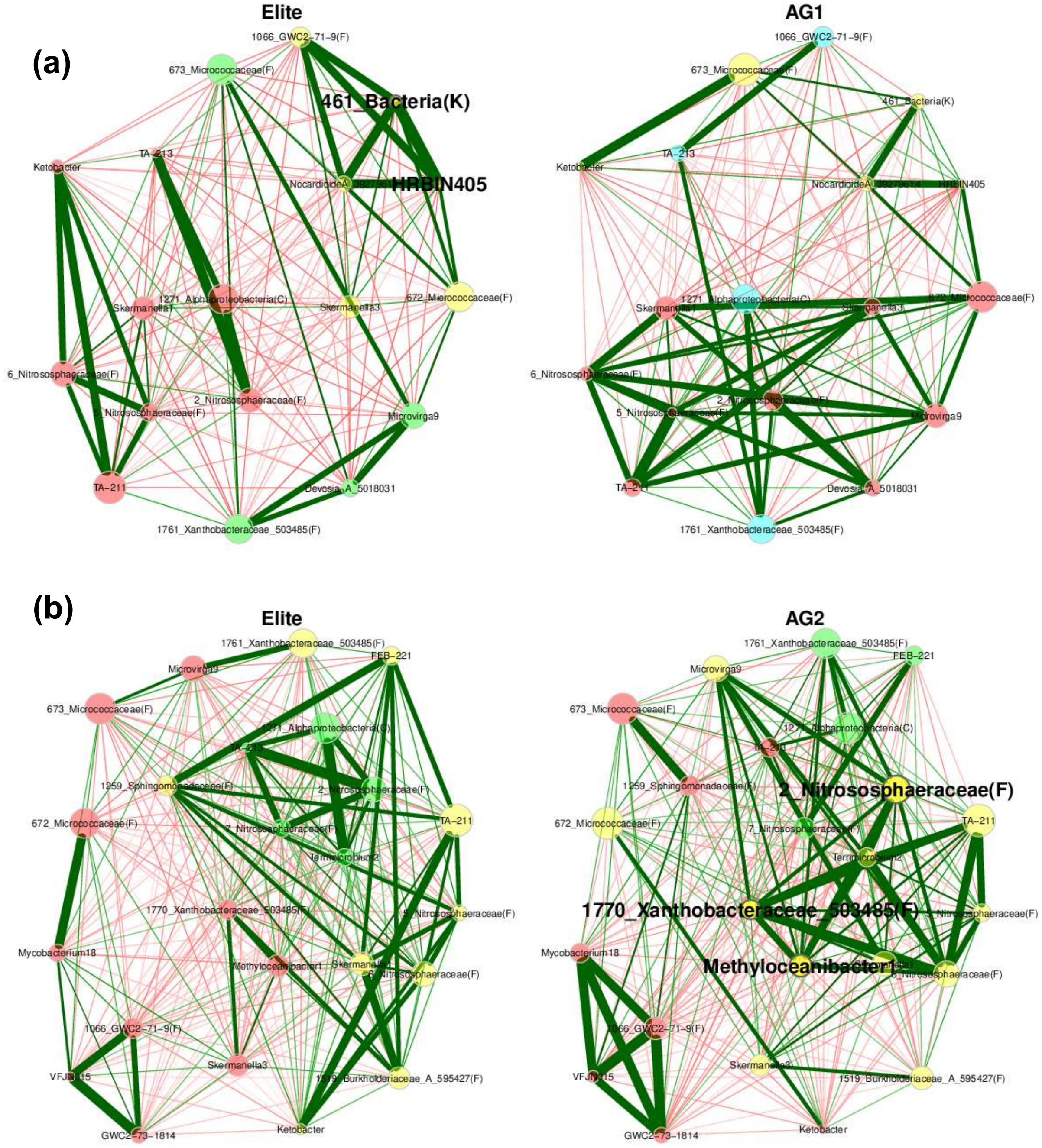

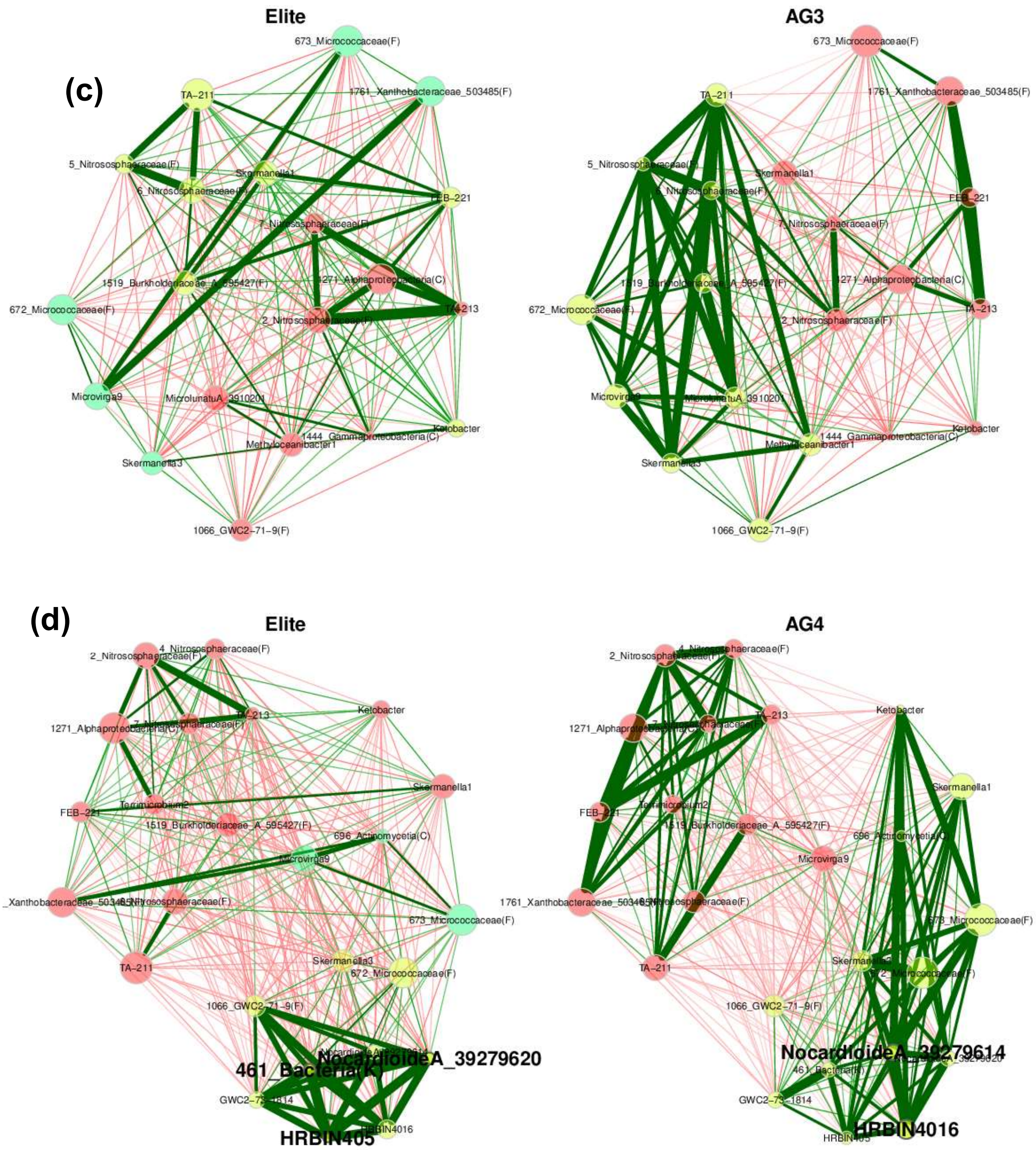
(c)
Elite
673_Micrococcaceae(F)
TA-211
1761_Xanthobacteraceae_503485(F)
5_Nitrososphaeraceae(F)
Skermanella1
6_Nitrososphaeraceae(F)
FEB-221
7_Nitrososphaeraceae(F)
1519_Burkholderiaceae_A_595427(F)
1271_Alphaproteobacteria(C)
672_Micrococcaceae(F)
TA-213
2_Nitrososphaeraceae(F)
Microvirga9
MicrolunatuA_391020
Ketobacter
1444_Gammaproteobacteria(C)
Methyloceanibacter1
Skermanella3
1066_GWC2-71-9(F)
AG3
673_Micrococcaceae(F)
TA-211
1761_Xanthobacteraceae_503485(F)
5_Nitrososphaeraceae(F)
Skermanella1
6_Nitrososphaeraceae(F)
FEB-221
7_Nitrososphaeraceae(F)
1519_Burkholderiaceae_A_595427(F)
1271_Alphaproteobacteria(C)
672_Micrococcaceae(F)
TA-213
2_Nitrososphaeraceae(F)
Microvirga9
MicrolunatuA_391020
Ketobacter
1444_Gammaproteobacteria(C)
Methyloceanibacter1
Skermanella3
1066_GWC2-71-9(F)
(d)
Elite
4_Nitrososphaeraceae(F)
2_Nitrososphaeraceae(F)
Ketobacter
1271_Alphaproteobacteria(C)
7_Nitrososphaeraceae(F)
TA-213
Skermanella1
FEB-221
Terrimicrobium2
1519_Burkholderiaceae_A_595427(F)
696_Actinomycetia(C)
Microvirga9
_Xanthobacteraceae_503485(F)
6_Nitrososphaeraceae(F)
673_Micrococcaceae(F)
TA-211
Skermanella3
672_Micrococcaceae(F)
1066_GWC2-71-9(F)
NocardioideA_39279620
461_Bacteria(K)
GWC2-73-1814
HRBIN4016
HRBIN405
AG4
4_Nitrososphaeraceae(F)
2_Nitrososphaeraceae(F)
Ketobacter
1271_Alphaproteobacteria(C)
TA-213
Skermanella1
FEB-221
Terrimicrobium2
1519_Burkholderiaceae_A_595427(F)
696_Actinomycetia(C)
Microvirga9
1761_Xanthobacteraceae_503485(F)
6_Nitrososphaeraceae(F)
673_Micrococcaceae(F)
TA-211
Skermanella3
672_Micrococcaceae(F)
1066_GWC2-71-9(F)
NocardioideA_39279614
NocardioideA_39279620
461_Bacteria(K)
GWC2-73-1814
HRBIN4016
HRBIN405

(e)

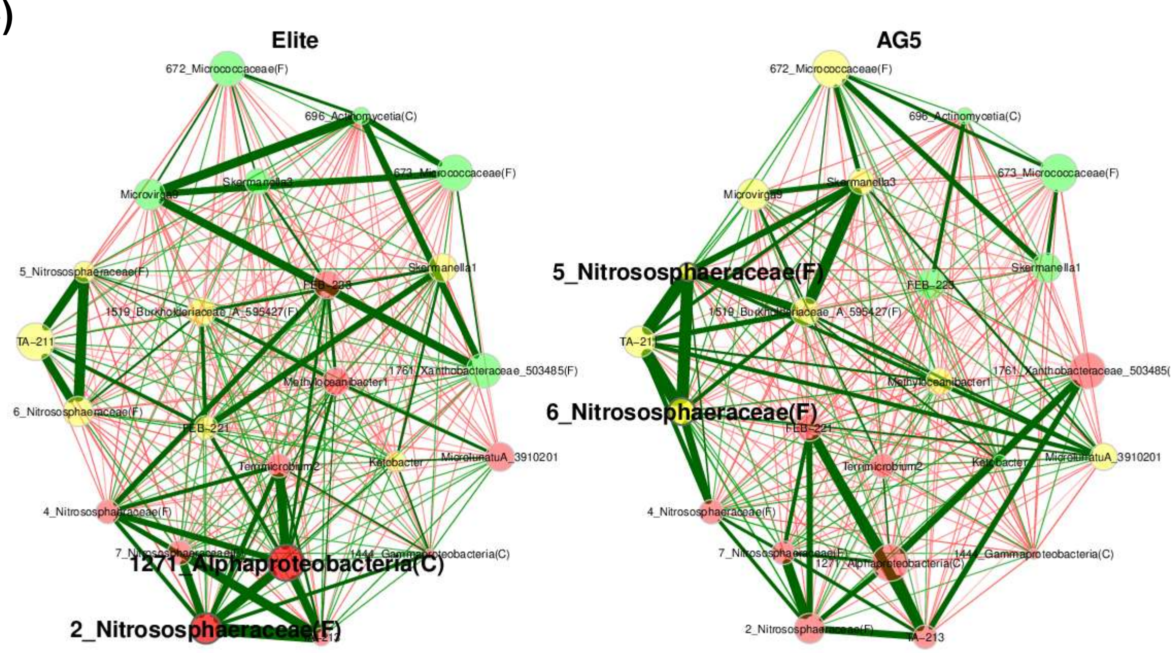
Elite
AG5
672_Micrococcaceae(F)
696_Actinomycetia(C)
673_Micrococcaceae(F)
Microvirga9
Skermanella3
5_Nitrososphaeraceae(F)
Skermanella1
FEB-223
1519_Burkholderiaceae_A_595427(F)
TA-211
1761_Xanthobacteraceae_503485(F)
Methyloceanibacter1
6_Nitrososphaeraceae(F)
FEB-221
Ketobacter
MicrolunatuA_3910201
Terrimicrobium2
4_Nitrososphaeraceae(F)
7_Nitrososphaeraceae(F)
1444_Gammaproteobacteria(C)
1271_Alphaproteobacteria(C)
2_Nitrososphaeraceae(F)
TA-213

(f)

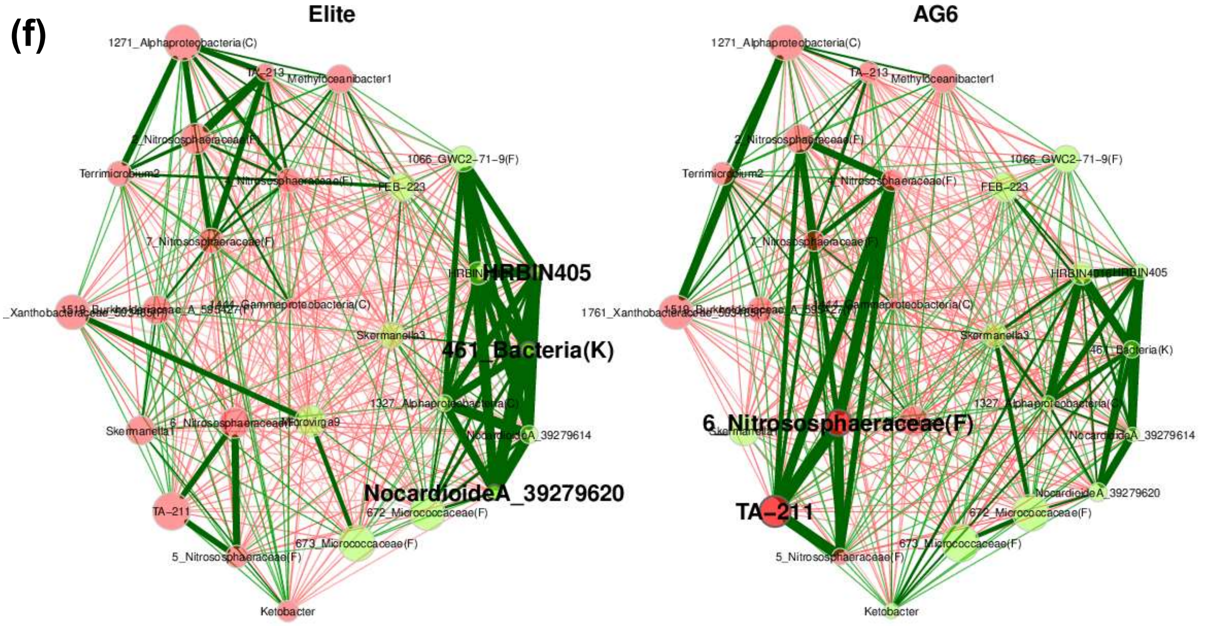
Elite
AG6
1271_Alphaproteobacteria(C)
TA-213
Methyloceanibacter1
2_Nitrososphaeraceae(F)
1066_GWC2-71-9(F)
Terrimicrobium2
4_Nitrososphaeraceae(F)
FEB-223
7_Nitrososphaeraceae(F)
HRBIN405
1761_Xanthobacteraceae_503485(F)
1444_Gammaproteobacteria(C)
Skermanella3
461_Bacteria(K)
1327_Alphaproteobacteria(C)
Skermanella1
6_Nitrososphaeraceae(F)
Microvirga9
NocardioideA_39279614
NocardioideA_39279620
TA-211
672_Micrococcaceae(F)
673_Micrococcaceae(F)
5_Nitrososphaeraceae(F)
Ketobacter

(g)

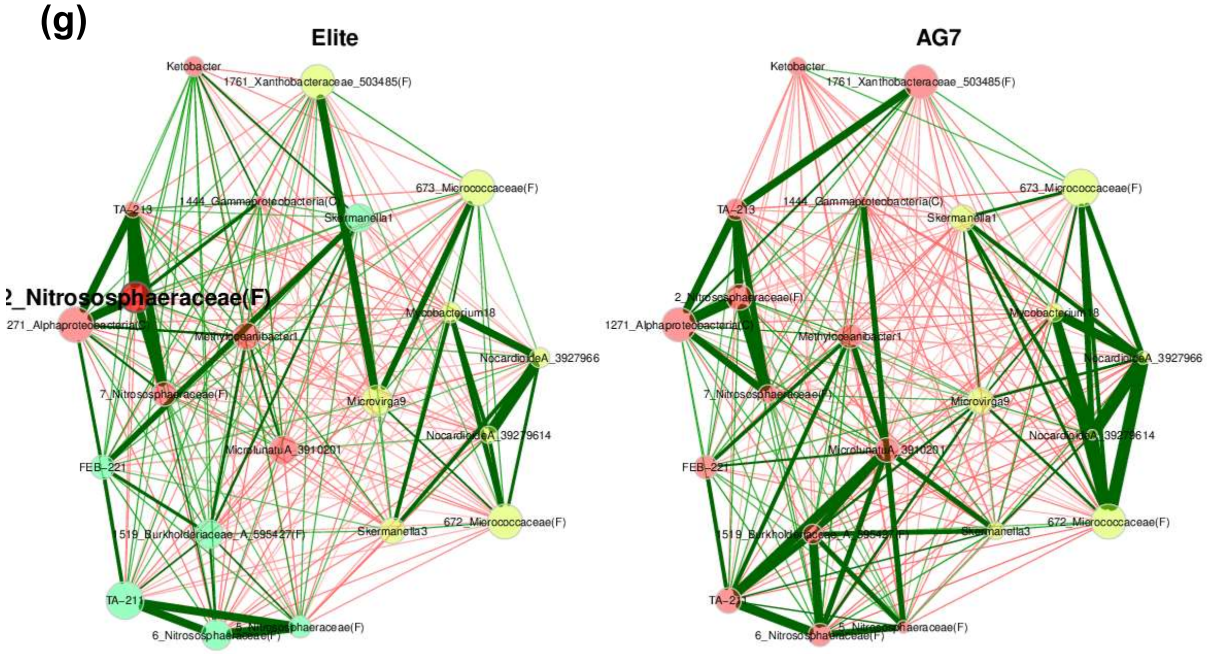
Elite
AG7
Ketobacter
1761_Xanthobacteraceae_503485(F)
673_Micrococcaceae(F)
TA-213
1444_Gammaproteobacteria(C)
Skermanella1
2_Nitrososphaeraceae(F)
1271_Alphaproteobacteria(C)
Methyloceanibacter1
Mycobacterium18
NocardioidesA_3927966
7_Nitrososphaeraceae(F)
Microvirga9
NocardioidesA_39279614
FEB-221
MicrolunatusA_3910201
672_Micrococcaceae(F)
1519_Burkholderiaceae_A_595427(F)
Skermanella3
TA-211
5_Nitrososphaeraceae(F)
6_Nitrososphaeraceae(F)

**Supplementary Tables Captions**

**Table S1**. Characterisation of the field soil properties where the 83 genotypes of hexaploid wheat were grown.

**Table S2**. Sample metadata used in this study, including the accession number of the A.E. Watkins landrace collection of bread wheat genotypes maintained at John Innes Centre, UK (https://www.jic.ac.uk/research-impact/germplasm-resource-unit/), ancestral groups (AGs) categories [2], genotype country and continent of origin and network cluster assignment by network dissimilarity analysis performed with the R package NetCoMi [5].

**Table S3**. Pairwise comparison between elite (modern) cultivars of hexaploid wheat and historic landraces grouped into ancestral groups (AG) [2], closely related genetically, using Bray-Curtis and Jaccard distances (R package micro4all [14]). The analysis assessed the dissimilarity among microbial communities in the rhizosphere soil of the wheat genotypes. Bray-Curtis considers both the presence and abundance of species, while Jaccard distance considers presence or absence. High coefficients and low p-values generally indicate strong statistical significance between the groups compared. The q-values are adjusted p-values using the false discovery rate approach.

**Table S4**. Alpha diversity indices (Shannon, Chao1, Inverse Simpson), Faith's phylogenetic diversity, species richness and evenness quantifying the variety and abundance of microbial taxa in the rhizosphere soil collected from accessions of hexaploid wheat genotypes of the A.E. Watkins landrace collection of bread wheat genotype maintained at John Innes Centre, UK (https://www.jic.ac.uk/research-impact/germplasm-resource-unit/) and two elite (modern) cultivars grown in the field. Alpha diversity indices were calculated using the R package phyloseq [6].

**Table S5**. Relative abundance of the top nine phyla identified in the rhizosphere soil of 81 genotypes of hexaploid wheat of the A.E. Watkins landrace collection of bread wheat genotypes maintained at John Innes Centre, UK (https://www.jic.ac.uk/research-impact/germplasm-resource-unit/), grouped into ancestral groups (AGs) [2], closely related genetically and two elite cultivars used as reference for modern genotypes. The relative abundance was obtained using the R package phyloseq [6].

**Table S6-S8**. Differential abundance in rhizosphere microbiome between hexaploid elite (modern) cultivars and historic landraces grouped into ancestral groups (AG) [2]. The LinDA (Linear models for Differential Abundance analysis, R package microbiomeStat [4]) output includes the list of differentially abundant taxa, their associated p-values, and adjusted p-values (using the false discovery rate approach) at phylum (Table S6), order (Table S7) and genus (Table S8) taxonomy levels.

**Table S9-S10**. Multivariate association between the abundance of microbial taxa in the rhizosphere and wheat genotypes grouped into ancestral groups (AG) [2], closely related genetically and elite (modern) cultivars. The Multivariable Association with Linear Models (R package Maaslin2 [11]) outputs include the list of differentially abundant taxa, their associated p-values, and adjusted p-values (using the false discovery rate approach) at phylum (Table S9) and genus (Table S10) taxonomy levels.

**Table S11**. Ecological roles of microbial taxa in the rhizosphere soil, for which we found distinct differences in abundance between 81 genotypes of hexaploid wheat grouped into ancestral groups (AG) [2] and two elite (modern) cultivars (see Fig. 3 and Supplementary Fig. 7).

**Table S12**. Functional analysis of microbial communities in the rhizosphere soil collected from 81 genotypes of hexaploid wheat genotypes grouped into ancestral groups (AG) [2], closely related genetically and two elite (modern) cultivars. KEGG [12,13] modules relative abundances for nitrogen metabolism pathways. **M00175** Nitrogen fixation, nitrogen => ammonia (4) (complete 1/1); **M00531** Assimilatory nitrate reduction, nitrate => ammonia (4) (complete 2/2); **M00530** Dissimilatory nitrate reduction, nitrate => ammonia (9) (complete 2/2); **M00529** Denitrification, nitrate => nitrogen (10) (complete 4/4); **M00528** Nitrification, ammonia => nitrite (4) (complete 2/2); **M00804** Complete nitrification, comammox, ammonia => nitrite => nitrate (6) (complete 3/3)

**Table S13**. Functional analysis of microbial communities in the rhizosphere soil collected from 81 genotypes of hexaploid wheat genotypes grouped into ancestral groups (AG) [2], closely related genetically and two elite (modern) cultivars. KEGG [12,13] modules relative abundances for carbon fixation pathways. **M00165** Reductive pentose phosphate cycle (Calvin cycle) (17) (complete 11/11); **M00168** Carbon fixation by Calvin cycle; **M00169** Carbon fixation by Calvin cycle; **M00173** Reductive citrate cycle (Arnon-Buchanan cycle) (32) (complete 10/10); **M00376** 3-Hydroxypropionate bi-cycle (21) (complete 13/13); **M00377** Reductive acetyl-CoA pathway (Wood-Ljungdahl pathway) (10) (complete 7/7); **M00579** Phosphate acetyltransferase-acetate kinase pathway, acetyl-CoA => acetate (4) (complete 2/2).

**Table S14**. Properties of the microbial co-ocurrence network constructed with NetComi [5] for the rhizosphere soil collected from 81 genotypes of hexaploid wheat of the A.E. Watkins landrace collection of bread wheat genotypes maintained at John Innes Centre, UK (https://www.jic.ac.uk/research-impact/germplasm-resource-unit/), grouped into ancestral groups (AGs) [2], closely related genetically, and two elite cultivars used as reference for modern genotypes (see Fig. 4a). The output of the network analysis includes global network properties, number of clusters, the identified hubs and the network centrality measures.

**Table S15-S21**. Properties of the network comparison performed with NetComi [5] for the rhizosphere soil collected from 81 genotypes of hexaploid wheat of the A.E. Watkins landrace collection of bread wheat genotypes maintained at John Innes Centre, UK (https://www.jic.ac.uk/research-impact/germplasm-resource-unit/), grouped into ancestral groups (AGs) [2], closely related genetically, and two elite cultivars used as reference for modern genotypes. The outputs of the network analysis microbiome include global network properties, number of clusters, the identified hubs, the network centrality measures and Jaccard comparison.

**Table S22**. Summary of Jaccard test results obtained from the network comparisons performed with NetCoMi [5] as presented in Supplementary Tables 15-21.

**Table S23**. Summary differences in hub taxa extracted from the network comparisons. performed with NetCoMi [5] as presented in Supplementary Tables 15-21.

**Table S24**. Properties of the dissimilatory network constructed with NetComi [5] for the rhizosphere soil collected from 81 genotypes of hexaploid wheat of the A.E. Watkins landrace collection of bread wheat genotypes maintained at John Innes Centre, UK (https://www.jic.ac.uk/research-impact/germplasm-resource-unit/), grouped into ancestral groups (AGs) [2], closely related genetically, and two elite cultivars used as reference for modern genotypes (see Fig. 4b). The output of the network analysis includes global network properties, number of clusters, the identified hubs and the network centrality measures.

### References


1. Barnett, D., Arts, I. & Penders, J. microViz: an R package for microbiome data visualization and statistics. *J. Open Source Softw.* **6**, 3201 (2021).
2. Cheng, S. *et al.* Harnessing landrace diversity empowers wheat breeding. *Nature* **632**, 823–831 (2024).
3. Nelkner, J. *et al.* Abundance, classification and genetic potential of Thaumarchaeota in metagenomes of European agricultural soils: a meta-analysis. *Environ Microbiome* **18**, 26 (2023).
4. Zhang, X., Chen, J. & Zhou, H. MicrobiomeStat: statistical methods for microbiome compositional data. (2022).
5. Peschel, S., Müller, C. L., von Mutius, E., Boulesteix, A.-L. & Depner, M. NetCoMi: network construction and comparison for microbiome data in R. *Brief. Bioinform.* **22**, bbaa290 (2021).
6. McMurdie, P. J. & Holmes, S. phyloseq: an R package for reproducible interactive analysis and graphics of microbiome census data. *PLoS One* **8**, e61217 (2013).
7. Kembel, S. W. *et al.* Picante: R tools for integrating phylogenies and ecology. *Bioinformatics* **26**, 1463–1464 (2010).
8. Kassambara, A. ggpubr:'ggplot2'based publication ready plots. *R package version* 2 (2018).
9. Wickham, H., Chang, W. & Wickham, M. H. Package 'ggplot2'. *Create elegant data visualisations using the grammar of graphics. Version* **2**, 1–189 (2016).
10. Lahti, L. & Shetty, S.
11. Mallick, H. *et al.* Multivariable association discovery in population-scale meta-omics studies. *PLoS Comput. Biol.* **17**, e1009442 (2021).
12. Kanehisa, M., Sato, Y., Kawashima, M., Furumichi, M. & Tanabe, M. KEGG as a reference resource for gene and protein annotation. *Nucleic Acids Res.* **44**, D457–62 (2016).
13. Kanehisa, M., Sato, Y. & Morishima, K. BlastKOALA and GhostKOALA: KEGG tools for

functional characterization of genome and metagenome sequences. *J. Mol. Biol.* **428**, 726–731 (2016).

14. Wentzien, M. N. Micro4all: microbiome for all. R package version 0.0.0.9000. https://rdrr.io/github/nuriamw/ (2023).
